# Ongoing genome doubling promotes evolvability and immune dysregulation in ovarian cancer

**DOI:** 10.1101/2024.07.11.602772

**Authors:** Andrew McPherson, Ignacio Vázquez-García, Matthew A. Myers, Matthew Zatzman, Duaa Al-Rawi, Adam Weiner, Samuel Freeman, Neeman Mohibullah, Gryte Satas, Marc J. Williams, Nicholas Ceglia, Allen W. Zhang, Jun Li, Jamie L.P. Lim, Michelle Wu, Seongmin Choi, Eliyahu Havasov, Diljot Grewal, Hongyu Shi, Minsoo Kim, Roland Schwarz, Tom Kaufmann, Khanh Ngoc Dinh, Florian Uhlitz, Julie Tran, Yushi Wu, Ruchi Patel, Satish Ramakrishnan, DooA Kim, Justin Clarke, Hunter Green, Emily Ali, Melody DiBona, Nancy Varice, Ritika Kundra, Vance Broach, Ginger J. Gardner, Kara Long Roche, Yukio Sonoda, Oliver Zivanovic, Sarah H. Kim, Rachel N. Grisham, Ying L. Liu, Agnes Viale, Nicole Rusk, Yulia Lakhman, Lora H. Ellenson, Simon Tavaré, Samuel Aparicio, Dennis S. Chi, Carol Aghajanian, Nadeem R. Abu-Rustum, Claire F. Friedman, Dmitriy Zamarin, Britta Weigelt, Samuel F. Bakhoum, Sohrab P. Shah

**Author notes:** equal contribution (Andrew McPherson, Ignacio Vázquez-García). Corresponding authors: Sohrab P. Shah Andrew McPherson.

## Abstract

Whole-genome doubling (WGD) is a critical driver of tumor development and is linked to drug resistance and metastasis in solid malignancies. Here, we demonstrate that WGD is an ongoing mutational process in tumor evolution. Using single-cell whole-genome sequencing, we measured and modeled how WGD events are distributed across cellular populations within tumors and associated WGD dynamics with properties of genome diversification and phenotypic consequences of innate immunity. We studied WGD evolution in 65 high-grade serous ovarian cancer (HGSOC) tissue samples from 40 patients, yielding 29,481 tumor cell genomes. We found near-ubiquitous evidence of WGD as an ongoing mutational process promoting cell-cell diversity, high rates of chromosomal missegregation, and consequent micronucleation. Using a novel mutation-based WGD timing method, `doubleTime`, we delineated specific modes by which WGD can drive tumor evolution: (i) unitary evolutionary origin followed by significant diversification, (ii) independent WGD events on a pre-existing background of copy number diversity, and (iii) evolutionarily late clonal expansions of WGD populations. Additionally, through integrated single-cell RNA sequencing and high-resolution immunofluorescence microscopy, we found that inflammatory signaling and cGAS-STING pathway activation result from ongoing chromosomal instability and are restricted to tumors that remain predominantly diploid. This contrasted with predominantly WGD tumors, which exhibited significant quiescent and immunosuppressive phenotypic states. Together, these findings establish WGD as an evolutionarily ‘active’ mutational process that promotes evolvability and dysregulated immunity in late stage ovarian cancer.

## INTRODUCTION

Whole-genome doubling (WGD) is found in >30% of solid cancers, leading to increased rates of metastasis, drug resistance and poor outcomes^1^. Often observed on a background of *TP53* mutation, genome doubling leads to increased chromosomal instability (CIN) and karyotypic diversification^2^. Several studies have reported that the fitness advantage of genome-doubled cells is conferred through its buffering effect on deleterious mutations^2–4^. *In vitro* studies indicate that genome doubling also leads to major phenotypic consequences, such as chromatin and epigenetic changes^5^, replication stress^6^ and cellular quiescence^5,6^. Previous studies of WGD in patient tumors used bulk whole-genome sequencing (WGS), which requires computational reconstruction of somatic evolutionary histories^7^, making cellular diversity difficult to infer. These analyses have tended to cast WGD as an early event in tumor evolution^8^, restricting its occurrence and mechanistic significance to an etiologic role^7^. However, live-cell analysis has previously suggested that errors in chromosome segregation often lead to cytokinesis failure and the ongoing generation of polyploid cells^9^, suggesting WGD might be an ongoing process during tumor evolution. Recent reports from *in vitro* and PDX models have demonstrated that temporal and evolutionary dynamics of genome doubling can be captured at single-cell resolution^10,11^. However, in the patient setting, how dynamical properties of genome doubling drive evolution and phenotypic state changes at the time of clinical presentation remains understudied. We contend that applying single cell approaches to clinical samples therefore opens the opportunity to ask new questions of how genome doubling evolution drives genomic diversity and phenotypic cellular states in the patient context.

We used single-cell whole-genome sequencing to study WGD in the context of high-grade serous ovarian cancer (HGSOC), a tumor type with ubiquitous *TP53* mutation and frequent WGD. We analyzed 65 untreated HGSOC samples at the time of diagnosis from 40 patients with single-cell whole-genome sequencing (29,481 tumor cell genomes) and site-matched immunofluorescent staining for markers of micronuclei and DNA sensing, known byproducts of chromosome segregation defects. Using this multi-modal approach, we conclude WGD is an ongoing mutational process which promotes evolvability through cell-cell diversity, high rates of CIN, and pervasive co-occurrence of cells with heterogeneous ploidy states within the same tumor. We delineated three modes of WGD evolution: (i) early fixation of a single event, (ii) late fixation of multiple independent WGD events, and (iii) emergence of late WGD clones. By linking genomic measurements with cellular phenotypes in previously generated site-matched single-cell RNA sequencing data^13^ we found that microenvironmental inflammatory signaling remains active in tumors that remain predominantly diploid in contrast to enriched quiescent and immunosuppressive states in predominantly WGD tumors. Our findings therefore point to WGD as a critical co-variate of inflammatory signaling and immunosuppression. Given our findings are derived from clinical samples at disease presentation, we suggest our study should further motivate and inform novel therapeutic targeting of WGD and CIN^12,13^.

## RESULTS

### Ovarian cancer patient cohort

Surgical specimens (*n*=65) from treatment-naive HGSOC patients (*n*=40) were collected from multiple sites during primary debulking surgery or laparoscopy, as previously described^14^ (**Fig. 1A**, **Extended Data Fig. 1A, Methods**). Patients were confirmed as advanced HGSOC by gynecologic pathologists. Sampled sites included adnexa (i.e. ovary and fallopian tube), omentum, peritoneum, bowel, and other intraperitoneal sites (**Extended Data Fig. 1B**). Clinical characteristics of all patients are summarized in **Extended Data Fig. 1B** and **Supp. Tab. 1**. Somatic and germline driver mutations were determined by MSK-IMPACT clinical sequencing, including ubiquitous somatic *TP53* loss, somatic and germline *BRCA1/2* loss, somatic *CDK12* mutation and somatic *CCNE1* amplification (**Extended Data Fig. 1B**)^14^. Mutational signatures derived from whole-genome sequencing included homologous recombination-deficient (HRD)-Dup (*BRCA1* mutant-like) and HRD-Del (*BRCA2* mutant-like) cases, as well as HR-proficient foldback inversion-bearing (FBI) and tandem duplicator (TD) tumors (18 HRD-Dup, 8 HRD-Del, 13 FBI, 1 TD) using integrated point mutation and structural variations, as previously described^11,14,15^.

**Figure 1.**
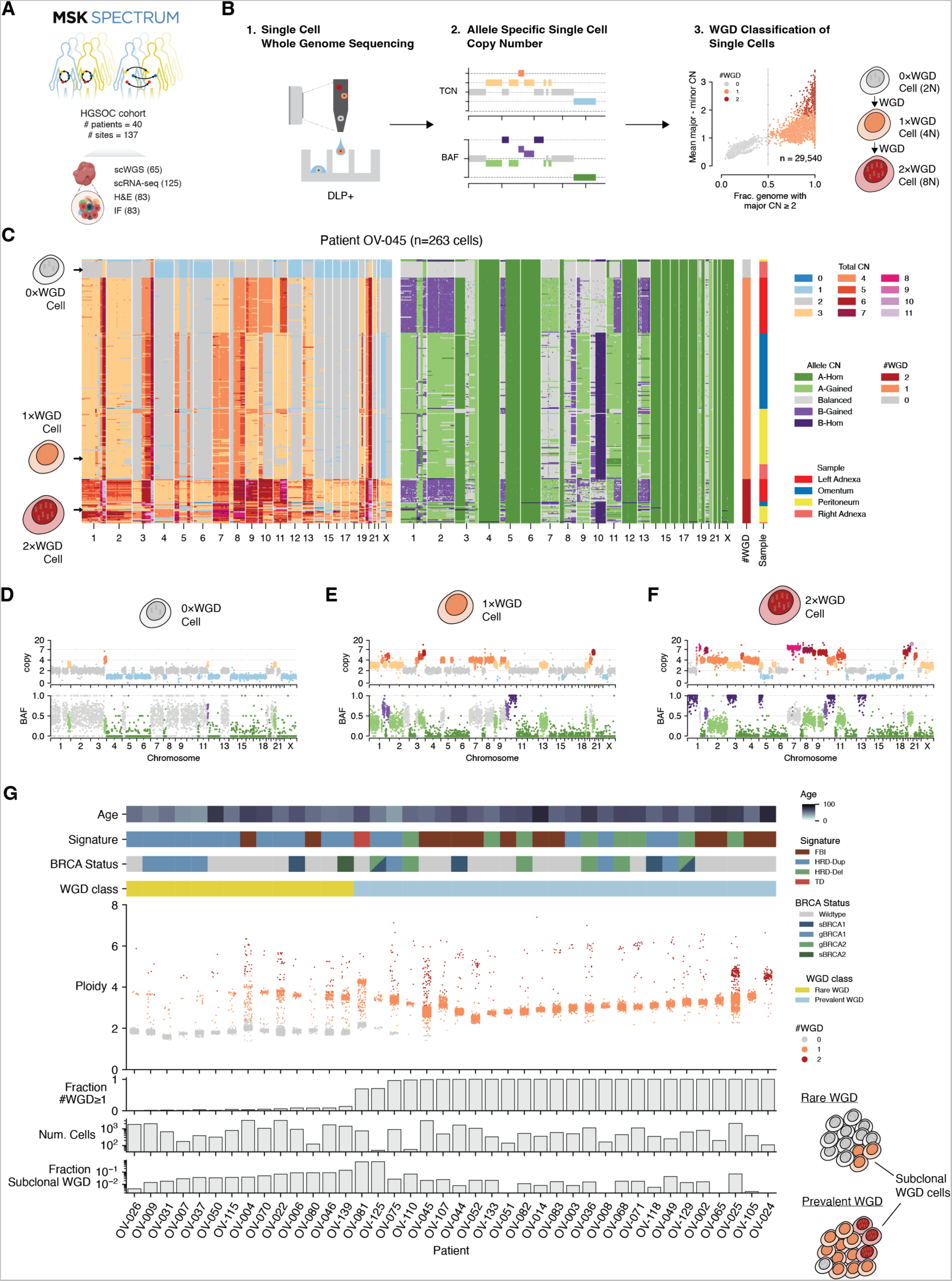
Whole genome doubling is a dynamic mutational process. **A.** Overview of the MSK SPECTRUM cohort and specimen collection workflow. **B.** Study design for analyzing cellular ploidy and WGD in single cells using scWGS with the DLP+ protocol. Right-hand plot shows classification of WGD multiplicity in cancer cells (# WGD=0, 1, or 2) using fraction of the genome with major CN ≥ 2 (x-axis) vs mean allele CN difference (y-axis). **C.** Heatmap of total (left) and allele specific (right) copy number for patient 045, with predicted #WGD and site of resection for each cell annotated. The dominant 1×WGD population was downsampled from 1,857 to 200 cells, and the full 0×WGD and 2×WGD populations numbering 18 and 44 cells respectively are shown. **D-F.** Example 0×WGD, 1×WGD, and 2×WGD cells from patient 045. **G.** Distribution of cell ploidy (middle y-axis) of individual cells for each tumor, colored by # WGD. Age at diagnosis, mutation signature, *BRCA1/2* mutation status, and WGD class are annotated at top; % WGD and number of cells per patient are annotated at bottom.

### Single-cell whole-genome sequencing and orthogonal phenotypic assays

Tumor-derived single-cell suspensions were flow-sorted to remove CD45^+^ immune cells, then subject to single-cell whole-genome sequencing (scWGS) using the Direct Library Preparation protocol^16^ (DLP+, **Methods**, **Supp. Tab. 2**). A total of 53,005 single-cell whole-genomes (median 1,345 per patient) were generated with median coverage depth of 0.078 per cell and median coverage breadth of 0.073 (**Extended Data Fig. 2A-B**) with 29,481 genomes admitted into analysis following quality control (**Methods**). In addition, whole-slide H&E and immunofluorescence (IF) images from adjacent formalin-fixed paraffin-embedded (FFPE) tissue sections were obtained for 37 out of 40 patients (**Supp. Tab. 2**). IF sections were assessed for DNA sensing mechanisms and genome sequencing-independent readouts of chromosomal instability (DAPI, cGAS) (**Methods**). In addition, we leveraged previously-generated single-cell RNA sequencing (scRNA-seq) data from both CD45^+^ and CD45^-^ compartments of 32 patients (52 scRNA samples site matched to scWGS)^14^, enabling genotype-phenotype analyses of these tissues. Together the dataset comprises a single-cell resolution multi-modal measurement of aneuploidies, genomic and chromosomal instability, and their cell-intrinsic and tumor microenvironment phenotypic readouts in HGSOC patient samples.

### Whole-genome doubling at single-cell resolution

Using the 29,481 high-quality single cell cancer genomes, we first investigated the distribution of WGD states across our cohort and within each tumor. We inferred the number of WGD events in the evolutionary history of each cell based on allele-specific copy-number profiles^17,18^ (**Fig. 1B**). The distribution of allele-specific copy number features showed clear separation between WGD states, permitting assignment of per-cell WGD multiplicities of 0 (0×WGD, 46% of cells), 1 (1×WGD, 53%) and 2 (2×WGD, 1%) (**Extended Data Fig. 2H-I**). The number of WGD events per cell correlated with both cell size measured through the optical components of DLP+ (**Extended Data Fig. 2J**), and mitochondrial copy number (**Extended Data Fig. 2K**), providing orthogonal validation based on known correlates of nuclear genome scaling^16,19^.

We then analyzed intra-patient WGD states at single-cell resolution, finding pervasive heterogeneity in WGD states within tumors. For example, patient 045 (**Fig. 1C**) simultaneously harbored a minority of 0×WGD (1%, **Fig. 1D**), a majority of 1×WGD (97%, **Fig. 1E**), and a minor fraction of 2×WGD cells (2%, **Fig. 1F**). Surprisingly, tumors with co-existing WGD states were present in 36/40 patients, including 31/32 of the patients with >200 tumor cells (**Fig. 1G**). In total, 5% of all tumor cells (*n*=1,481 cells), were part of non-dominant WGD states across the cohort (median of 2.6% of cells per patient; **Extended Data Fig. 2L**). As each patient’s tumor was typically dominated by a single WGD state (the dominant state comprising >85% of cells for 38/40 patients), we dichotomized each tumor as either Prevalent WGD: harboring >50% of 1×WGD or 2×WGD cells (26/40 patients); or Rare WGD: with ≥50% 0×WGD cells (14/40 patients). Prevalent WGD patients comprised 60% of the cohort, were older, and were enriched for FBI and HRD-Del mutation signature patients, consistent with previous bulk genome sequencing studies^7,14^ (**Extended Data Fig. 2M-P**). Thus, while average signals corroborate previous bulk estimates of WGD prevalence across patients^17^, single cell analysis established that WGD exists as a distribution over 0×WGD, 1×WGD, 2×WGD cells within tumors, with at least 2 co-existing WGD states observed in the majority of patients.

### Evolutionary histories of WGD clones

We next analyzed evolutionary histories of WGD and non-WGD clones including timing the origin of single or multiple WGD expansions within each patient. We developed doubleTime, a multi-step computational approach that (i) uses a scWGS mutation caller to predict SNVs (Articull, manuscript in prep.), (ii) identifies SNV clones using SBMClone^20^, (iii) constructs a clone phylogeny, (iv) places WGD events on branches of the phylogeny, and (v) infers mutation timing including the relative timing of WGDs on the WGD branches (**Fig. 2A, Methods**). For 23 out of 25 Prevalent WGD patients, a single ancestral WGD event was common to the dominant 1×WGD population of cells (**Fig. 2B**). For two patients (025 and 045) we observed coexisting WGD clones from distinct WGD events, consistent with lineage divergence in the ancestral diploid population followed by expansion of independent WGD clones (**Fig. 2C, Extended Data Fig. 3A-B**). Within both patients, the multiple WGD events were predicted to occur at approximately the same mutation time in the tumor’s life history. Remarkably, for both of these patients, WGD clones coexisted in multiple anatomic sites. All WGD clones were present in both right adnexa and omentum of 025. For patient 045, the left adnexa harbored one of the three of the WGD clones whereas the right adnexa, omentum, and peritoneal tumors were mixtures of all three WGD clones. The remaining 14 patients did not show evidence of expanded WGD clones through SNV analysis (**Fig. 2D**), although all harbored small populations of 1×WGD cells (**Fig. 2D**).

**Figure 2.**
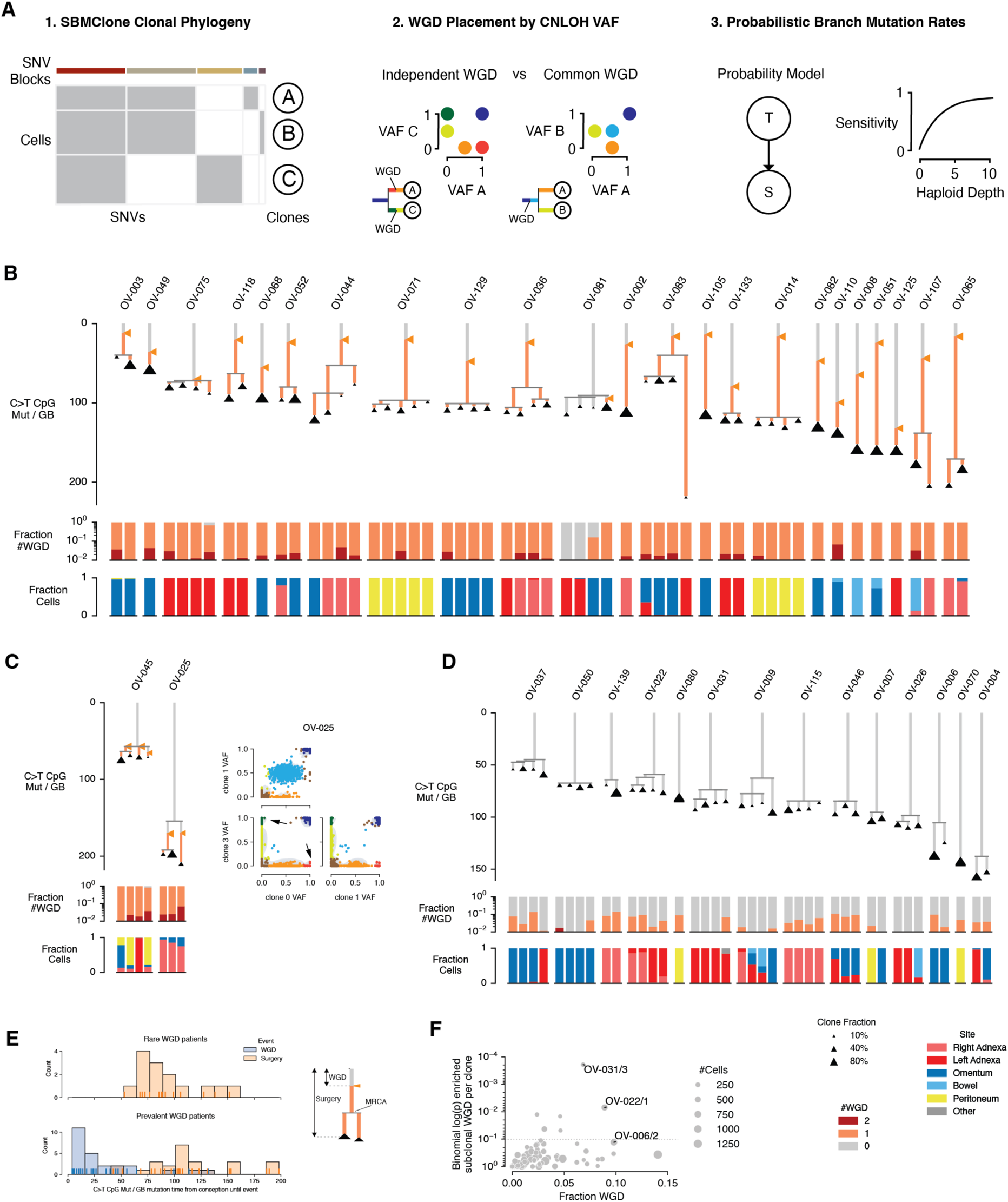
Evolutionary timing of WGD events from single nucleotide variants. **A.** Schematic of the approach for timing WGDs in SNV clones. SBMClone is used to infer clones based on SNVs, and a phylogeny is constructed from presence/absence patterns of SNVs across SNV clones (left). For each pair of WGD clones, independence of the WGD is determined through analysis of the SNV VAF in clonal cnLOH regions (center). Predictions of independent vs common WGD are used to place WGD events in the tree. A probabilistic method is used to assign SNVs to the tree including placing SNVs before or after WGD events (right). The method models the relationship between depth of coverage and SNV sensitivity to account for clones of differing size. **B-D.** Clone phylogenies for the 39 patients for which the SNV based method could be applied. Length of branches show the number of age-associated SNVs (C to T at CpG) assigned to each branch, adjusted for coverage-depth-related reduction in SNV sensitivity. Clone sizes as a fraction of the patient’s total sequenced cells are shown by the size of the triangle for each leaf. Clonal WGD events are represented as orange triangles at the predicted location along WGD branches, and branches are colored according to the number of WGD at that point in the evolutionary history. The fractions of each clone with each #WGD state and from each sampled site are shown below each clone tree. Each patient is annotated with mutation signature and age at diagnosis. **E.** Histogram and rug plot showing the sensitivity-adjusted age-associated SNV count for WGD and diagnosis events for rare WGD (top) and prevalent WGD (bottom) patients. **F.** Fraction of +1 WGD cells within each clone (x axis) and log binomial *p*-value for the test that a clone has a greater fraction of +1 WGD cells than the overall +1 WGD fraction for the patient.

We then investigated the evolutionary timing of WGD clonal expansions to determine if they fixed early in tumorigenesis, or whether clonal expansions of WGD cells occurred throughout their life histories. Prevalent WGD patients exhibited increased mutation time from fertilization to surgical resection (total C>T CpG burden) vs Rare WGD patients, similar to WGD vs non-WGD patients in previous bulk WGS analyses^7^ (**Fig. 2E**). However, while bulk genome sequencing studies of ovarian tumors reported early acquisition of WGD^7^, we found that later WGD clonal expansions inferred from scWGS were common, with 7/25 WGD events occurring more than 50% of the way through the tumors life history. For three of the late WGD patients (045, 075 and 081), WGD events approximately coincided with the most recent common ancestor (MRCA). (**Fig. 2B-C,E**). These same three patients, in addition to late WGD patient 125, all exhibited extant populations of 0×WGD cells: 17 (0.9%) 0×WGD cells for 045 (**Extended Data Fig. 3A**), 34 (3.9%) 0×WGD cells for 075 (**Extended Data Fig. 3B**), 14 (27%) 0×WGD cells for 125 (**Extended Data Fig. 3C**), and 38 0×WGD cells in the omentum sample of 081 which contained the site-specific WGD expansion (**Fig. 2B**). Thus, in these four patients, more recently expanded WGD clones co-existed with extant cells from the 0×WGD population from which they were derived.

The existence of a substantial fraction of subclonal WGD subpopulations (1×WGD in 0×WGD clones, 2×WGD in 1×WGD clones) across multiple clones was consistent with parallel and ongoing WGD. We investigated if these rare subclonal WGD cells shared common mutations, indicative of late WGD-associated clonal expansions (**Fig. 2B-D,F**). In patient 025, a small subpopulation of 43 2×WGD cells harbored 325 SNVs specific to the 2×WGD cells (**Extended Data Fig. 3D**). Subclonal WGD expansions in patients 031 (7 cells) and 006 (27 cells) were too small to be detected by SNV analysis, but could be identified by copy-number events shared across multiple subclonal WGD cells (**Extended Data Fig. 3E-F**). For other patients, subclonal WGD cells were evenly distributed across multiple clones, indicative of continual WGD across clonal populations. Together, quantifying the evolutionary history of WGD and chromosomal instability in single cells revealed distinct modes of ongoing WGD evolution: (i) diploid tumors with a background rate of unexpanded WGD cells, (ii) tumors with evolutionary late WGD expansions including parallel expansion of multiple WGD clones, and (iii) evolutionary early WGD tumors with a single dominant WGD clone.

### Post WGD genomic diversification

Leveraging single-cell-resolution measurements, we next asked how WGD promotes genomic diversification and evolvability. We first quantified cell-to-cell genomic heterogeneity using pairwise nearest-neighbor copy-number distance (NND) for each cell (**Methods**, **Extended Data Fig. 4A**). Mean NND increased with WGD multiplicity and was highest for subclonal WGD populations, with 1×WGD populations in Rare WGD patients exhibiting higher mean NND than 1×WGD populations in Prevalent WGD patients, and 2×WGD cells in predominantly 1×WGD tumors exhibiting the highest cell-cell diversity (**Fig. 3A**). Some Prevalent WGD patients exhibited surprising levels of diversity: in 8 patients, cells were on average different for 10% of the genome when compared with the most similar cell. The empirical distribution of NND values had a heavy tail (**Extended Data Fig. 4B**) with unexpected enrichment for highly divergent cells with very distinct copy-number profiles. We defined divergent cells as those with NND greater than the 99th percentile of a Beta distribution fit to the NND values (**Fig. 3B**). The CN profiles of divergent cells resembled the previously reported ‘hopeful monsters’ found in colorectal cancer organoids^21^, suggesting they may be the product of unstable tetraploid cells undergoing multipolar mitosis (**Fig. 3C**). When compared with clonal CN profiles, divergent cells harbored elevated rates of whole chromosome and chromosome arm loss, uniformly distributed across the genome (**Extended Data Fig. 4C-D**), accompanied by a significant rate of arm and chromosome nullisomy for both Rare and Prevalent WGD patients (**Extended Data Fig. 4E**). Notably, these divergent cells were present in 39/40 patients (mean 2.8% of cells), with higher rates in Prevalent WGD patients suggesting an increased propensity for abnormal mitoses (**Fig. 3D**). Interestingly, the three patients with large clonal expansion of late WGD (081, 045 and 025), ranked first, fourth and seventh highest in divergent cell fraction. Furthermore, patient 049 had the second-highest divergent cell fraction and the third most recent clonal WGD behind two of the three WGDs in 045. Overall, our data is concordant with previous evidence suggesting WGD cell populations sustain a period of instability following WGD, which can result from an increase in the number of centrosomes^22^.

**Figure 3:**
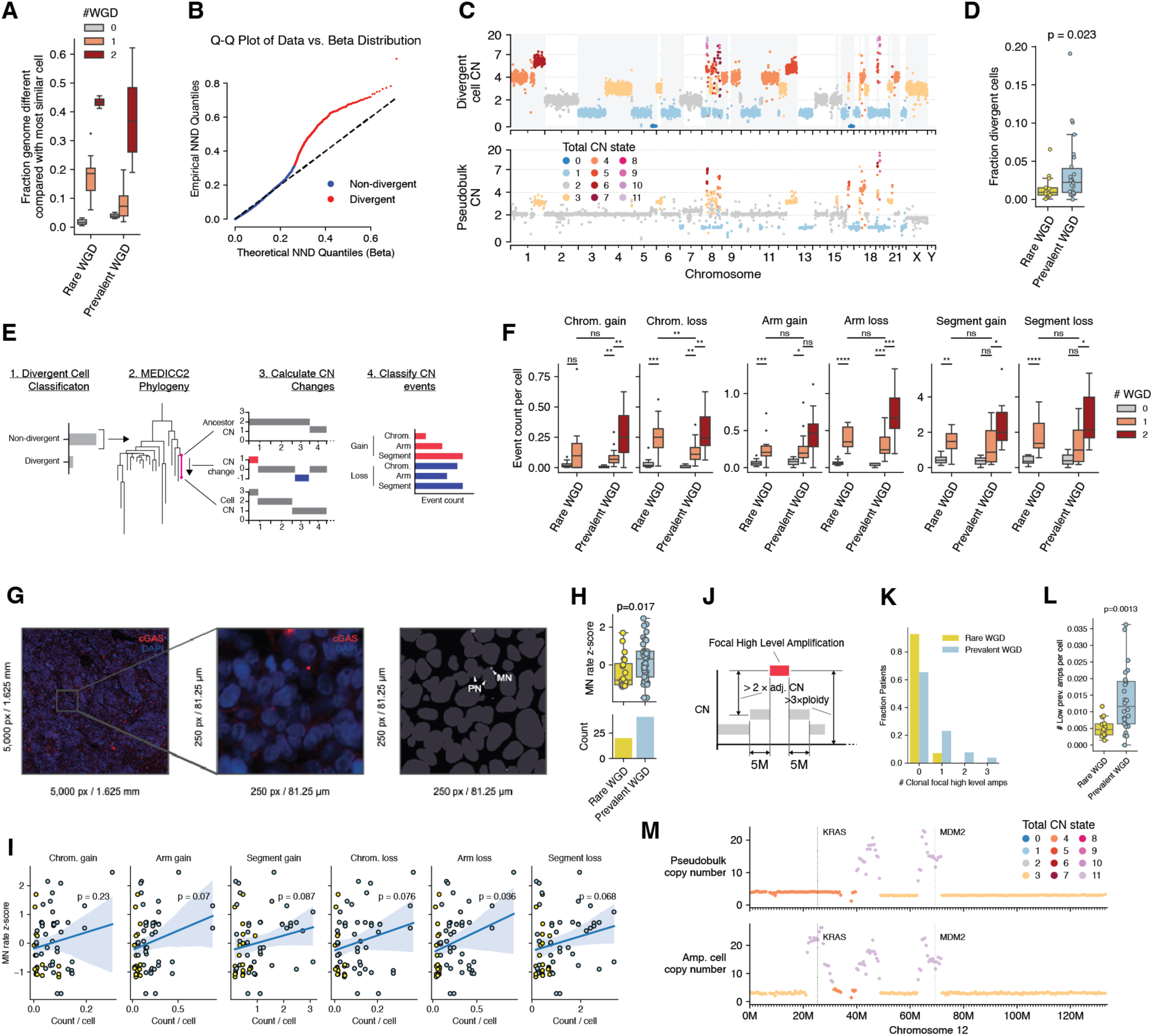
Impact of WGD on rates of chromosomal instability at single-cell resolution. **A.** Divergence as measured by nearest neighbor distance, where distance is represented as the fraction of the genome with different CN. NND is calculated for each population of cells within each patient. Boxplots show the mean NND for each WGD population within each patient. **B.** QQ plot of a beta fit (x-axis) vs empirical (y-axis) quantiles of NND values for all cells in the cohort. Divergent cells, defined as outliers (>99 percentile) of the beta distribution, are shown in red. **C.** CN profile of an example divergent cell from patient 004 (top) compared to pseudobulk CN of all cells for that patient (bottom). Shaded regions show differences between cell and pseudobulk CN. **D.** Fraction of divergent cells in Rare vs Prevalent WGD patients. **E.** Method for computing cell specific events in non-divergent cells. **F.** Event counts per cell for loss and gain of chromosomes, arms, and large segments, split by #WGD state and Prevalent vs Rare WGD patient status. Mann-Whitney U test significance is annotated as ‘ns’: 5.0×10^-2^ < *p* ≤ 1.0, ‘*’: 1.0×10^-2^ < *p* ≤ 5.0×10^-2^, ‘**’: 1.0×10^-3^ < *p* ≤ 1.0×10^-2^, ‘***’: 1.0×10^-4^ < *p* ≤ 1.0×10^-3^, ‘****’: *p* ≤ 1.0×10^-4^. **G.** Left: Low-magnification IF image of FFPE tumor section from a representative HGSOC patient, stained with DAPI (DNA) and anti-cGAS antibody. Middle: High-magnification inset. Right: cGAS segmentation mask of MN in the foreground and DAPI segmentation mask of PN in the background. **H.** Z-scored MN rate split by Prevalent vs Rare WGD patient status. **I.** Patient CN event counts per cell for loss and gain of chromosomes, arms, and large segments (x-axis) compared with slide specific Z-scored MN rate (y-axis). Points are colored by Prevalent vs Rare WGD patient status. **J.** Diagram defining focal high-level amplification. **K.** Count of clonal focal high-level amplifications per patient split by Prevalent vs Rare WGD. **L.** Count per cell of low-prevalence focal high-level amplifications split by Prevalent vs Rare WGD. Low prevalence was defined as occurring in 2 or more cells but less than 10% of the patient cell population. **M.** Example low prevalence focal high-level amplification found in patient 002 (bottom) and not detectable in the pseudobulk copy number of the same patient.

To investigate rates of chromosome missegregation events, we computed copy number alterations in each cell (excluding divergent cells) accrued since its immediate ancestor in a phylogeny inferred for each patient (**Fig. 3E, Extended Data Fig. 4F, Methods**). This enabled inference of rates of cell-specific (and therefore most recent) copy number changes. The rate (counts per cell) of gains and losses of whole chromosomes, chromosome arms, and segments (>15MB) increased with WGD multiplicity across all event types (**Fig. 3F**). We recomputed a ploidy-adjusted version of the gain and loss rates that accounted for the increased opportunity for copy-number events in higher ploidy cells. The ploidy-adjusted rates (counts per cell per GB) showed similar increases, highlighting that the rate differences were not merely a function of increased chromosome number but were instead indicative of systemic changes in post-WGD cells (**Extended Data Fig. 4G, Methods**). 1×WGD subpopulations had higher ploidy-adjusted rates of chromosome losses in Rare WGD patients than Prevalent WGD patients, suggesting more recently emerging WGD cells were more prone to chromosome loss events, or that early WGD populations had stabilized. For instance, ploidy-adjusted chromosome losses were 4.1 times more abundant in Rare WGD 1×WGD cells compared to Rare WGD 0×WGD cells (*p*=5.6×10^-4^, Mann-Whitney U test), 2.3 times more abundant in Prevalent WGD 1×WGD cells compared to Rare WGD 0×WGD cells (*p*=5.6×10^-4^, Mann-Whitney U test), and 1.8 times more abundant in Rare WGD 1×WGD cells compared to Prevalent WGD 1×WGD cells (*p*=0.016, Mann-Whitney U test).

One of the phenotypic consequences of chromosome segregation errors is the formation of micronuclei, which are chromosome or chromosome arm containing structures that are distinct from the primary nucleus during interphase. Micronuclear envelopes are rupture-prone, often exposing their enclosed genomic double-stranded DNA (dsDNA) to the cytoplasm^23–25^, leading to activation of innate immune signaling driven by the cytosolic dsDNA sensing cGAS-STING pathway^26^. We asked whether the propensity of chromosome missegregation correlates with micronuclei formation via high-resolution immunofluorescence microscopy of cGAS and DAPI staining on FFPE sections, site-matched to scWGS datasets. We used a deep learning approach to automatically detect primary nuclei (PN) and cGAS^+^ ruptured micronuclei (MN), enabling whole-slide quantification of MN-to-PN ratios at scale (1,779,351 PN and 83,352 ruptured MN from 61 quality-filtered IF images of slides obtained for 31 patients, **Fig. 3G**, **Methods**). Ruptured micronuclei per primary nucleus (MN rate) ranged from 0.005 to 0.31 (median 0.05) and was 2 times higher in Prevalent WGD patients (*p*<0.01, **Fig. 3H, Methods**), with MN rate showing modest correlation with rates of cell specific copy number change (**Fig. 3I**). Thus, WGD-related copy number change associates with the formation of micronuclei and raises the possibility that micronuclei are a vehicle for losses and segmental amplifications in HGSOC^27^.

Abnormalities in micronuclei have been proposed as a mechanism for the formation of complex chromosomal rearrangements, chromothripsis and extrachromosomal DNA, all of which can lead to elevated rates of oncogene amplification. We found that clonal (>90% cells) high-level amplification (HL Amps, **Methods**, **Fig. 3J**) were more frequent in Prevalent WGD patients (*p*=0.028 Mann-Whitney U test, **Fig. 3K**), including events amplifying *MDM2* (002), *CCNE1* (105), *ERBB2* (044, 051), *CCND1* (065), and *CCND3* (083). Only 1 of the 14 clonal HL Amps was found in a Rare WGD patient, involving the *CCNE1* gene in patient 004, corroborated by bulk sequencing^7^ (**Extended Data Fig. 4H**). We classified HL Amps as low cancer cell fraction (CCF) if they occurred in less than 10% of the patient cell population. Prevalent WGD was associated with a significantly higher number of low-CCF HL Amps per cell compared to Rare WGD (*p*=0.022 Mann-Whitney U test, **Fig. 3L**), highlighting that the mutational process that generates HL Amps may itself be increased in Prevalent versus Rare WGD patients. Many of the low prevalence HL Amps are undetectable at a bulk level, highlighting the need for single cell data to identify these events (**Fig. 3M**).

Taken together, multiple forms of cell-to-cell genome diversification, including chromosomal missegregations, multipolar mitoses, ruptured micronuclei and HL Amps, all exhibited elevated rates in Prevalent WGD patients, firmly linking WGD to increased cellular genomic diversification in HGSOC.

### Evolutionary dynamics of WGD and non-WGD clones

Given the increased rate of chromosomal instability associated with WGD, we next used a phylogenetic approach to investigate the impact of this instability on tumor evolution and the extent to which WGD promotes punctuated vs gradual evolutionary change (**Fig. 4A**, **Methods**). We first focused on events predicted to be on the ancestral branches of each patient. These events were divided into those inferred to occur (i) after WGD in the ancestral branches of prevalent WGD patients (post-WGD) (ii) before WGD in ancestral branches of rare WGD patients (pre-WGD) or (iii) on the ancestral branches of rare WGD patients (non-WGD) (**Fig. 4B**). Rates of losses and gains of arms and chromosomes were significantly higher post-WGD relative to pre-WGD or non-WGD ancestral branches. Thus WGD was associated not only with increased rates of CIN, but also increased propensity for fixation of the changes resulting from CIN. Gains of arms and especially whole chromosomes were rare pre-WGD or on non-WGD ancestral branches, and were significantly more prevalent post-WGD (**Fig. 4B**). This highlights that commonly observed pseudo-triploid karyotypes are unlikely to arise through incremental gains on a diploid background. Instead, triploidy in HGSOC most likely results from WGD and both pre- and post-WGD losses.

**Figure 4.**
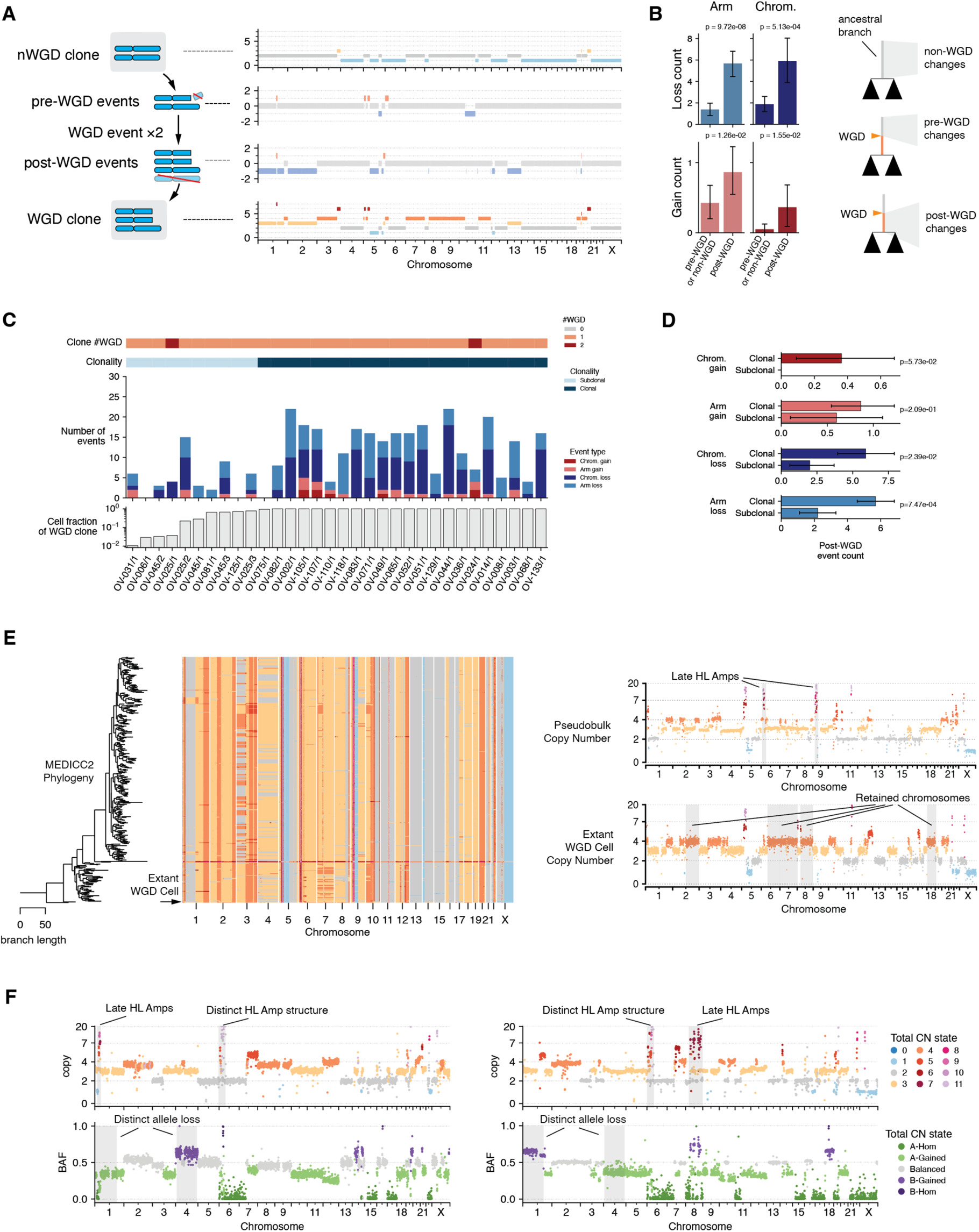
Modes of evolution post WGD. **A.** Pre- and post-WGD events illustrated for the ancestral branch of patient 044. **B.** Counts of ancestral arm and chromosome events detected across the cohort, grouped into pre- or post-WGD. **C.** Counts of arm and chromosome events occurring post-WGD for all high-confidence clonal and subclonal WGD events detected across the cohort, split by clonality of the WGD (cell fraction threshold 0.99). **D.** Boxplots summarizing C annotated with *p*-values (Mann-Whitney U test). **E.** MEDICC2 tree and copy number for patient 014 (left). The outgroup cell is shown with missing focal HL Amps (bottom right) compared with the majority of cells represented as a pseudobulk (upper right) for which there have been additional losses and focal HL Amps. **F.** CN (top) and allelic imbalance (bottom) for two divergent WGD clones from patient 083 (left, right) with shared WGD origin. Late and divergent HL Amps and losses of distinct alleles are highlighted.

To investigate punctuated vs gradual evolution post-WGD, we interrogated the post-WGD events for all high-confidence clonal (>95% of cells) and subclonal (<95% of cells) WGD clones (**Fig. 4C**). Comparing post-WGD events between clonal and subclonal WGD clones, we found clonal WGDs to have significantly more post-WGD events of all types (**Fig. 4D**). For instance, clonal WGDs accrued on average 3 times as many whole chromosome losses compared to subclonal WGDs. The number of events post-WGD for some subclonal WGD was surprisingly low. For example, the large WGD subclone (70% of cells) in patient 081 exhibited only two arm loss events post-WGD, while the many cells with more highly divergent genomes post-WGD were unique, possibly indicative of their comparable lack of fitness and inability to expand. In clonal WGDs, the number of chromosome losses and arm gains and losses were significantly correlated with the age of the WGD as estimated using mutations (**Methods**, **Extended Data Fig. 4J**). Patient 014 with a clonal WGD exemplifies post-WGD evolution (**Fig. 4E**). In this patient, a single cell distinct from the majority of the cells in the patient shared several post-WGD copy number changes with the majority population, but lacked 2 focal HL Amps common to the remaining cells, and retained 4 copies post-WGD of 2q, 6q, 7, 8q, and 18, all of which had 3 copies in the majority of cells. Thus, this outlying cell represented an intermediate stage of evolution post-WGD, likely outcompeted by the other cells with additional focal HL Amps and arm and chromosome losses. Post-WGD divergent evolution with clone specific HL Amps and parallel allele-specific losses was also observed, as exemplified by patient 083 (**Fig. 4F**).

### WGD-specific cell intrinsic and tumor microenvironment phenotypes

We next investigated phenotypic associations with WGD states to understand the cancer cell-intrinsic, stromal, and immune activation states found in HGSOC, leveraging previously published patient and site matched scRNA-seq data^14^. We first compared the fraction of cancer cells in the G1, S and G2/M phase of the cell cycle. Prevalent WGD samples exhibited a lower proportion of S-phase cells and a higher proportion of G1-phase cells, consistent with a slower proliferation rate and elongated G1 progression through the cell cycle (**Fig. 5A**, **Methods**)^28^. Pseudotime inference of cell cycle trajectories revealed divergent cell cycle progression in Prevalent vs Rare WGD tumors (**Fig. 5B,C**, **Extended Data Fig. 5A-E, Methods**). In particular, MCM complex genes involved in licensing of DNA replication origins at the G1/S transition (*MCM2*, *MCM6*) were expressed earlier in the cell cycle in Prevalent WGD tumors, together with factors involved in MCM complex loading such as *CDC6* (**Fig. 5D-E**), thus facilitating the replication of larger genomes. Mitotic cyclins (*CCNE1*) and genes involved in DNA repair (*BRCA2*, *MSH2*) had altered temporal order in association with WGD. We also observed correlation between cell cycle distribution and chromosomal missegregation event rates in a WGD-specific manner (**Fig. 5F**), where the fraction of cells in G1 was highly correlated with rates of chromosome losses and arm losses and gains in Rare WGD patients, but not in Prevalent WGD patients (**Fig. 5G, Extended Data Fig. 5G**). This might be due to the well-documented G1 cell cycle arrest that occurs upon chromosome missegregation and which must be overcome for cells to tolerate CIN^29,30^, an evolutionary milestone that is likely achieved by clones that have undergone WGD.

**Figure 5.**
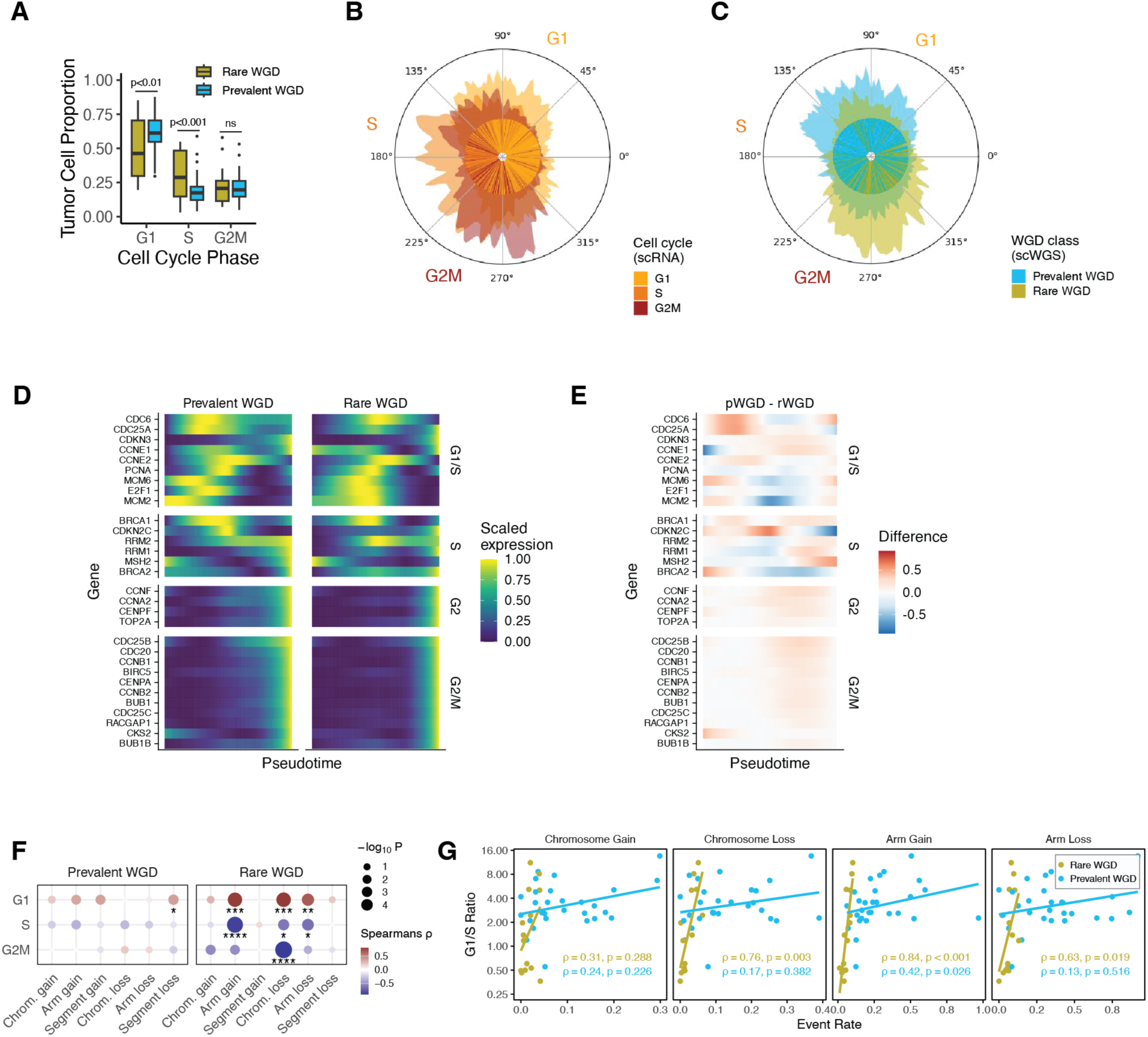
Cell cycle progression in the context of whole genome doubling. **A.** Proportion of cancer cells (y-axis) grouped by cell cycle phase (x-axis) in Prevalent WGD vs Rare WGD tumors (color). **B-C.** Cell cycle pseudotime inference in cancer cells. Inner ring shows cell cycle pseudotime in cancer cells and outer ring shows smoothed density estimate. B: Cell cycle assignment. C: Pseudotime grouped by Prevalent WGD (cyan) and Rare WGD (yellow) tumors. **D.** Scaled expression of phase-specific genes in Prevalent vs Rare WGD tumors as a function of cell cycle pseudotime. **E.** Differences in scaled gene expression of phase-specific genes in Prevalent vs Rare WGD tumors as a function of cell cycle pseudotime. **F.** Dotplot of correlations between missegregation rates derived from scWGS and cell cycle phase from scRNA in site-matched samples. **G.** Scatter plot of G1/S cell count ratios (y-axis) by rates (counts per cell) of large chromosomal changes (x-axis) split by Rare and Prevalent WGD (color). Regression coefficients and significance results are shown separately for Rare and Prevalent WGD.

We next proceeded to investigate the association between WGD and cancer cell-intrinsic immune signaling. Cells in Prevalent WGD tumors showed a significant decrease in Type I (IFN-α/IFN-β) and Type II (IFN-γ) interferon, inflammatory pathways, complement and TNFa/NF-κB signaling (**Fig. 6A**). We investigated how chromosomal instability phenotypes encoded in a CIN gene expression signature^31^ related to WGD state and found this was significantly higher in Prevalent WGD (**Fig. 6A**), likely due to the elevated missegregation rates as observed in scWGS. Interestingly, STING (*TMEM173*), an innate immune response gene activated by the presence of cytosolic DNA, was expressed at significantly lower levels in Prevalent WGD (**Fig. 6B**), suggesting STING expression may be repressed in Prevalent WGD tumors to evade the immunostimulatory effects of CIN^32–34^. In the context of Rare WGD, STING expression showed strong positive correlation with rates of missegregation, especially chromosome losses (**Fig. 6C**). In addition, expression of E2F target genes showed strong negative correlation with chromosome losses in Rare WGD (**Fig. 6D**). We validated our findings in an hTERT-immortalized retinal pigment epithelial (RPE1) cell line (**Methods**). In diploid RPE1 cells (**Extended Data Fig. 6D-E**), treatment with nocodazole and reversine resulted in increasing levels of chromosome and arm losses and gains (**Extended Data Fig. 6F**), in addition to concomitant increases in G1 cell fraction (**Extended Data Fig. G**) and STING expression (**Extended Data Fig. 6H**). Next we compared non-WGD cells in a later passage with a spontaneously arising WGD clone present in the same sample (**Extended Data Fig. 6i-J, Methods**). We found no difference in cell cycle fractions (**Extended Data Fig. 6K**), but STING expression was decreased in the WGD clone (**Extended Data Fig. 6L**). In the Rare WGD context, our results are concordant with the hypothesis that CIN-associated cytosolic DNA activates NF-κB, which promotes transcription of STING^35^, and suppresses E2F targets^36,37^, ultimately leading to G1 arrest or delay. Critically, we note that this cascade only appears to hold in Rare WGD tumors, suggesting signal re-wiring in Prevalent WGD tumors that enables highly chromosomally unstable tumor cells to adapt to ongoing chromosome missegregation events, thereby evading anti-tumor immune surveillance, as recently proposed^38^.

**Figure 6.**
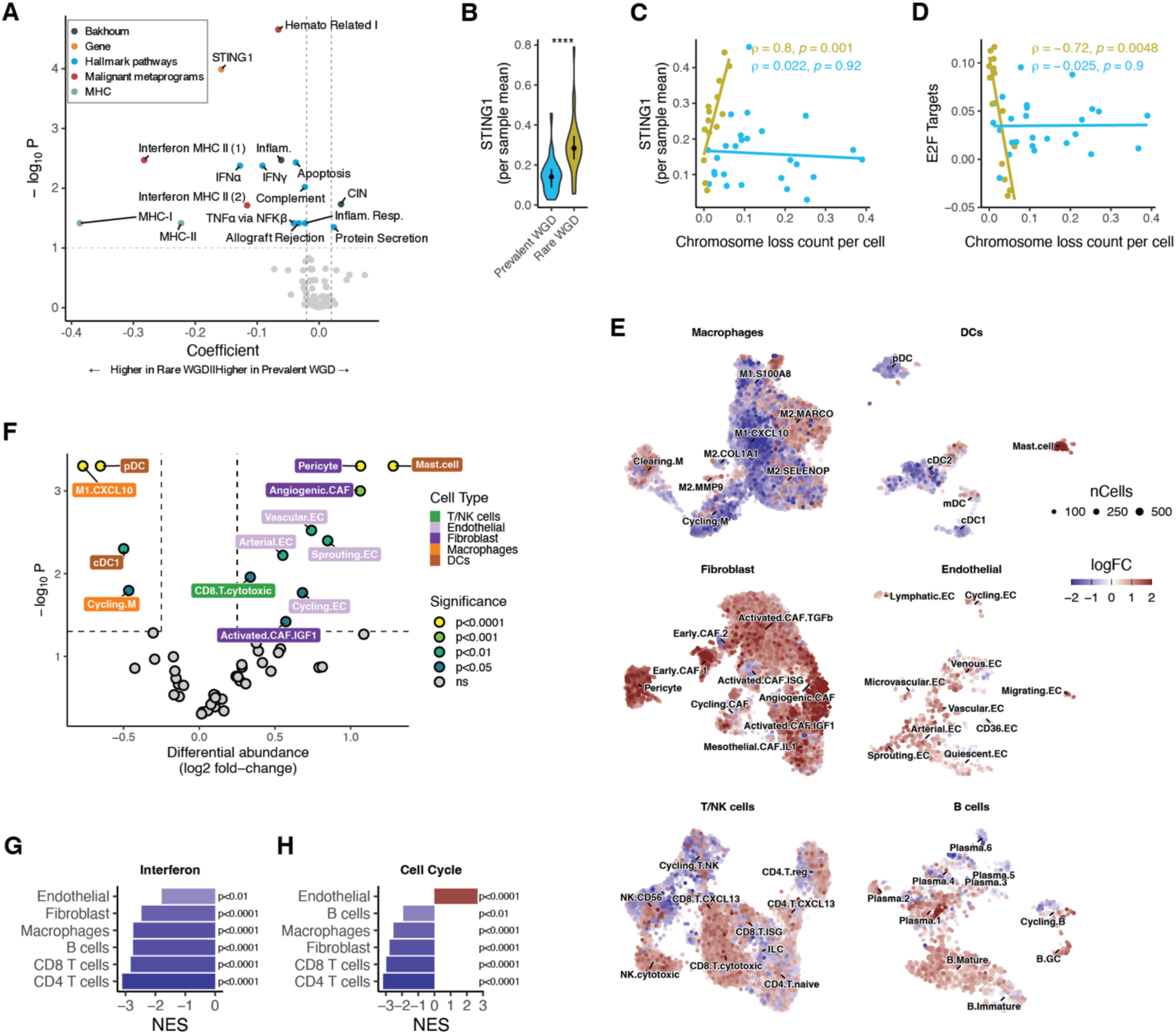
Tumor cell phenotypes and microenvironment remodeling in the context of whole genome doubling. **A.** Scatter plot depicting regression coefficients (x-axis) and significance (y-axis) for selected genes and pathways in Prevalent versus Rare WGD tumor cells. **B.** Per-sample mean expression of STING1 in Prevalent and Rare WGD samples. **C.** Scatter plot of STING1 gene expression (y-axis) by rate (counts per cell) of chromosomal losses (x-axis) split by Rare and Prevalent WGD (color). Regression coefficients and significance results are shown separately for Rare and Prevalent WGD patients. **D.** Scatter plot of hallmark E2F module score (y-axis) by rate (counts per cell) of chromosomal losses (x-axis) split by Rare and Prevalent WGD (color). Regression coefficients and significance are shown separately for Rare and Prevalent WGD patients. **E.** UMAP showing differential cell state enrichment in Prevalent versus Rare WGD samples in different TME cell types. **F.** Differential cell-type abundance testing results for cell types in Prevalent versus Rare WGD samples. **G.** Normalized enrichment scores (NES) for the interferon pathway across TME cell types. **H.** Normalized enrichment scores (NES) for the cell cycle pathway across TME cell types.

We next analyzed the impact of WGD and chromosome missegregation on the tumor immune microenvironment (TME). Consistent with the increased expression of IFN-stimulated genes (ISGs) observed in the cancer cells of Rare WGD patients, we observed enrichment of CXCL10^+^CD274^+^ macrophages (M1.CXCL10), IFN-producing plasmacytoid (pDCs), and activated dendritic cells (cDC1) in the microenvironments of Rare WGD tumors (**Fig. 6E-F**). All major cell types surveyed showed significant enrichment of ISGs in Rare WGD tumors, indicating a pro-inflammatory immune response (**Fig. 6G**). By contrast, the TME of Prevalent WGD tumors showed enrichment for endothelial cells, pericytes, and cancer-associated fibroblasts (CAFs) (**Fig. 6F**), along with suppression of ISG expression. Prevalent WGD tumors also showed slight enrichment of cytotoxic CD8^+^ T cells, possibly due to mutually exclusivity between cytotoxic CD8^+^ T cells and CXCL10^+^CD274^+^ macrophages across the cohort (**Extended Data Fig. 5C**). Notably, all major cell types in the TME of Prevalent WGD tumors, except for endothelial cells, exhibited marked depletion in cell cycle related gene expression, consistent with a pro-angiogenic yet immunosuppressive microenvironment in WGD tumors (**Fig. 6H**).

## DISCUSSION

Using single-cell whole-genome sequencing matched with scRNA and tissue-based quantification of ruptured micronuclei, we illuminate the significant impact of WGD on tumor evolvability and identify associations with cell cycle regulation, inflammatory signaling, angiogenesis and immunosuppressive phenotypes. Through evolutionary timing of WGD, we find variation in clonal WGD expansions from very early to late^7^, in addition to both subclonal WGD expansions and multiple independent WGD expansions that would be difficult to identify from bulk sequencing data^18^. Interestingly, for two patients with independent WGD clones, the WGD events were approximately synchronous in the tumor’s life history. The remaining patients could be classified into three groups: those with small fractions of non-clonal WGD cells, those with a late-emerging non-dominant WGD clone, and those with a dominant WGD clone. Importantly, we did not observe coexisting clones with varying WGD timing. These findings suggest that the evolutionary history of WGD in HGSOC is characterized by the rapid expansion of WGD clones, likely driven by changes in the fitness landscape that favor their proliferation.

The established relationship between WGD and genomic diversification is especially evident in our data, wherein we find ubiquitous presence of minor populations that have undergone additional doublings, an increased rate of cell-specific aneuploidies post-WGD, and profoundly divergent cells for which WGD has led to extreme instability. Evolutionary analysis of our data indicates that gradual losses, rather than punctuated evolution, shape the post-WGD evolution of many WGD clones, suggesting historical adaptation and tolerance for the high CIN levels associated with WGD. While WGD was associated with increased cGAS^+^ ruptured micronuclei, as expected given the higher levels of CIN, prevalent WGD tumors showed decreased cell-intrinsic and cell-extrinsic interferon signaling. In Rare WGD tumors, strong correlations between CIN and tumor phenotypes were consistent with CIN-dependent G1 elongation and increased STING transcription indicating an active cGAS-STING response in Rare WGD patients. This is in contrast with Prevalent WGD, which did not exhibit CIN-associated cell cycle alterations or STING expression increases. Thus STING transcriptional repression may be a prerequisite for clonal expansion of WGD. Furthermore, given the very early timing of WGD in some patients, our results also suggest that deactivation of STING may also be an early event in the evolutionary history of some HGSOC tumors, and may predate WGD. Future investigations into therapeutic targeting of the cGAS-STING pathway should consider WGD-specific abrogation of this pathway, as well as heterogeneity in WGD states.

Our results rely in part on the ability to accurately identify ploidy at single cell resolution. Several lines of evidence support the veracity of our results. As shown, many of the cells that make up the small fraction of subclonal WGD cells in each tumor are highly divergent and/or harbor homozygous regions, and would be unlikely to be either a miscalled doublet or poor quality copy number. The less divergent cells still show copy number changes in addition to a perfect doubling, given the requirement of at least one 10MB or larger segment with copy number state 1, 3 or 5. Given the expectation that WGD should be associated with additional post-WGD instability, we were surprised to find that some of the low-prevalence and late-emerging WGD clones did not have large amounts of post-WGD copy number change. This suggests that we may in fact be underestimating the number of small WGD clones, as some of those clones may not be marked by common post-WGD changes. Individual cells that have sustained perfect doublings and non-aberrant G2 phase cells would be detected as half their true ploidy in this study. Future methods may allow isolation of these cell populations, providing further insight into the dynamics of WGD.

Critically, we show that potentially targetable therapeutic vulnerabilities such as high-level oncogene amplification preferentially occur on a WGD background, and therefore arise in the context of a low-inflammation and immunosuppressive tumor microenvironment. Those tumors were primarily composed of 1×WGD and 2×WGD cells with increased immunosuppressive properties. We speculate that even if selective targeting of focal oncogene amplification^39^ were successful, immunosuppressive states may persist. As therapeutic stratification of patients by genomic properties gains traction in HGSOC, our data introduces a critical covariate given that nearly every tumor harbors WGD cells and multiple co-existent WGD states. Even with the modest cohort size presented here, we anticipate that studying how expanded WGD clones intersect with homologous recombination deficiency and impact responsiveness to anti-angiogenic therapies such as bevacizumab, will advance the rational administration of therapeutic strategies for HGSOC^40,41^. Moreover, targeting the WGD process itself may be required to prevent emergence of newly acquired WGD clones. The relevance of our findings to other tumor types remains unclear, although breast PDX^11^, *in vitro*^10^ and pancreatic cancer mouse^42^ studies suggest that WGD dynamics may be pervasive across *TP53* mutant cancers with implications for diverse mechanisms of therapeutic resistance^43^. Thus studying the role of WGD throughout the life history of a tumor should be prioritized as a determinant of therapeutic response.

## Data availability

Publicly accessible and controlled access data generated and analyzed in this study are documented in Synapse (accession number syn25569736). Raw sequencing data for scWGS data will be available from the NCBI Sequence Read Archive prior to publication. 10x 3’ scRNA-seq is available from the NCBI Gene Expression Omnibus (accession number GSE180661). scWGS copy number heatmaps can be visualized on Synapse (https://www.synapse.org/#!Synapse:syn51769919/datasets/). In addition, MEDICC2 trees and SBMClone results are provided as supplementary file spectrum-trees.html. IF images will be available from Synapse prior to publication.

## Code availability

The pipeline to process DLP+ scWGS is available at https://github.com/mondrian-scwgs. SIGNALS^11^ was used for most plotting and scWGS analysis and is available at https://github.com/shahcompbio/signals. `doubleTime` is available at https://github.com/shahcompbio/doubleTime.

## Acknowledgements

This project was funded in part by Cycle for Survival supporting Memorial Sloan Kettering Cancer Center and the Halvorsen Center for Computational Oncology. S.P.S. holds the Nicholls Biondi Chair in Computational Oncology and is a Susan G. Komen Scholar. This work was funded in part by awards from the Ovarian Cancer Research Alliance (OCRA) Collaborative Research Development Grant [648007] and NIH R01 CA281928-01 to S.P.S., OCRA Ann Schreiber Mentored Investigator Award to I.V.-G. [650687], OCRA Liz Tilberis Award to D.Z., the Department of Defense Congressionally Directed Medical Research Programs to S.P.S., D.Z. and B.W [W81XWH-20-1- 0565], the Seidenberg Family Foundation and the Cancer Research UK Cancer Grand Challenges Program to S.P.S. [C42358/A27460], the Marie-Josée and Henry R. Kravis Center for Molecular Oncology and the National Cancer Institute (NCI) Cancer Center Core Grant [P30-CA008748]. S.F.B. is funded by NIH/NCI grants [P50CA247749, DP5OD026395, R01CA256188, P30-CA008748], the Department of Defense Congressionally Directed Medical Research Program [BC201053], Burroughs Wellcome Fund (BWF), Josie Robertson Foundation, Pershing Square Sohn Alliance for Cancer Research, and Mary Kay Ash Foundation. B.W. is funded in part by Breast Cancer Research Foundation and NIH/NCI P50 CA247749 01 grants. D.Z. is funded by NIH grant R01 CA269382. A.C.W. is supported by NCI Ruth L. Kirschstein National Research Service Award for Predoctoral Fellows F31-CA271673. R.F.S. is a Professor at the Cancer Research Center Cologne Essen (CCCE) funded by the Ministry of Culture and Science of the State of North Rhine-Westphalia. R.F.S. was partially funded by the German Ministry for Education and Research as BIFOLD-Berlin Institute for the Foundations of Learning and Data [01IS18025A and 01IS18037A].

## Competing interests

B.W. reports grant funding by Repare Therapeutics paid to the institution, outside the submitted work, and employment of a direct family member at AstraZeneca. C.A. reports grants from Clovis, Genentech, AbbVie and AstraZeneca and personal fees from Tesaro, Eisai/Merck, Mersana Therapeutics, Roche/Genentech, Abbvie, AstraZeneca/Merck and Repare Therapeutics, outside the scope of the submitted work. C.A. reports clinical trial funding to the institution from Abbvie, AstraZeneca, and Genentech/Roche; participation on a data safety monitoring board or advisory board: AstraZeneca, Merck; unpaid membership of the GOG Foundation Board of Directors and the NRG Oncology Board of Directors. C.F. reports research funding to the institution from Merck, AstraZeneca, Genentech/Roche, Bristol Myer Squibb, and Daiichi; uncompensated membership of a scientific advisory board for Merck and Genentech; and is a consultant for OncLive, Aptitude Health, Bristol Myers Squibb and Seagen, all outside the scope of this manuscript. D.S.C. reports membership of the medical advisory board of Verthermia Acquio Inc and Biom’up, is a paid speaker for AstraZeneca, and holds stock of Doximity, Moderna, and BioNTech. D.Z. reports institutional grants from Merck, Genentech, AstraZeneca, Plexxikon, and Synthekine, and personal fees from AstraZeneca, Xencor, Memgen, Takeda, Astellas, Immunos, Tessa Therapeutics, Miltenyi, and Calidi Biotherapeutics. D.Z. own a patent on use of oncolytic Newcastle Disease Virus for cancer therapy. N.A.-R. reports grants to the institution from Stryker/Novadaq and GRAIL, outside the submitted work. R.N.G. reports funding from GSK, Novartis, Mateon Therapeutics, Corcept, Regeneron, Clovis, Context Therapeutics, EMD Serono, MCM Education, OncLive, Aptitude Health and Prime Oncology, outside this work. S.F.B. owns equity in, receives compensation from, and serves as a consultant and the Scientific Advisory Board and Board of Directors of Volastra Therapeutics Inc. He also serves on the SAB of Meliora Therapeutics Inc. S.P.S. reports research funding from AstraZeneca and Bristol Myers Squibb, outside the scope of this work; S.P.S. is a consultant and shareholder of Canexia Health Inc. Y.L.L. reports research funding from AstraZeneca, GSK/Tesaro, Artios Pharma, and Tesaro Therapeutics outside this work. Y.L. reports serving as a consultant for Calyx Clinical Trial Solutions outside this work.

## METHODS

### Experimental methods

#### Sample collection

All enrolled patients were consented to an institutional biospecimen banking protocol and MSK-IMPACT testing^44^, and all analyses were performed per a biospecimen research protocol. All protocols were approved by the Institutional Review Board (IRB) of Memorial Sloan Kettering Cancer Center. Patients were consented following the IRB-approved standard operating procedures for informed consent. Written informed consent was obtained from all patients before conducting any study-related procedures. The study was conducted in accordance with the Declaration of Helsinki and the Good Clinical Practice guidelines (GCP).

We collected fresh tumor tissues from 40 HGSOC patients at the time of upfront diagnostic laparoscopic or debulking surgery. Ascites and tumor tissue from multiple metastatic sites, including bilateral adnexa, omentum, pelvic peritoneum, bilateral upper quadrants, and bowel were procured in a predetermined, systemic fashion (median of 4 primary and metastatic tissues per patient) and were placed in cold RPMI for immediate processing. Blood samples were collected pre-surgery for the isolation of peripheral blood mononucleated cells (PBMCs) for normal whole-genome sequencing (WGS). The isolated cells were frozen and stored at −80°C. In addition, tissue was snap frozen for bulk DNA extraction and tumor WGS. Tissue was also subjected to formalin fixation and paraffin-embedding (FFPE) for histologic, immunohistochemical and multiplex immunophenotypic characterization.

#### Sample processing

We profiled patient samples using five different experimental assays:

1. Viably frozen single-cell suspensions were derived from fresh tissue samples and processed for single-cell whole-genome sequencing (scWGS) of 65 sites from 40 patients (~815 cells per site, **Supp. Tab. 2**). CD45^-^ cells were flow-sorted in samples with low tumor purity.
2. CD45^+^ and CD45^-^ flow-sorted cells were previously reported fresh tissue samples and processed for single-cell RNA sequencing (scRNA-seq) of 123 sites from 32 patients (~6k cells per site, **Supp. Tab. 2**).
3. For each specimen with scRNA-seq, site-matched FFPE tissue sections adjacent to the H&E section were stained by multiplexed immunofluorescence (IF) for micronuclei and DNA sensing mechanisms (83 tissue samples from 37 patients).
4. FDA-approved clinical sequencing of 468 cancer genes (MSK-IMPACT) was obtained on DNA extracted from FFPE tumor and matched normal blood specimens for each patient (**Extended Data Fig. 1B**).
5. Snap-frozen tissues were processed to obtain matched tumor-normal bulk whole-genome sequencing (WGS) on a single representative site of 33 out of 40 patients with scWGS, scRNA-seq and IF, to derive mutational processes from genome-wide single nucleotide and structural variants.

### Single-cell DNA sequencing

#### Tissue dissociation

Tumor tissue was immediately processed for tissue dissociation. Fresh tissue was cut into 1 mm pieces and dissociated at 37°C using the Human Tumor Dissociation Kit (Miltenyi Biotec) on a gentleMACS Octo Dissociator. After dissociation, single-cell suspensions were filtered and washed with Ammonium-Chloride-Potassium (ACK) Lysing Buffer. Cells were stained with Trypan Blue and cell counts and viability were assessed using the Countess II Automated Cell Counter (ThermoFisher). For detailed protocol see Bykov et al., 2020^45^. Freshly dissociated cells were processed for scRNA-seq as described in Vázquez-García et al., 2022^14^. Viably frozen dissociated cells were stored for scWGS.

#### Cell sorting

Viably frozen dissociated cells used for scWGS were thawed and then stained with a mixture of GhostRed780 live/dead marker (TonBo Biosciences) and Human TruStain FcX™ Fc Receptor Blocking Solution (BioLegend). For samples with low tumor purity, the stained samples were then optionally incubated and stained with Alexa Fluor® 700 anti-human CD45 Antibody (BioLegend). Post staining, they were washed and resuspended in RPMI + 2% FCS and submitted for cell sorting. The cells were sorted into CD45 positive and negative fractions by fluorescence assisted cell sorting (FACS) on a BD FACSAria™ III flow cytometer (BD Biosciences). Positive and negative controls were prepared and used to set up compensations on the flow cytometer. Cells were sorted into tubes containing RPMI + 2% FCS for sequencing.

#### Library preparation and sequencing

Single-cell whole-genome library preparation was carried out as described in Laks et al., 2019^16^. Briefly, single cells were dispensed into nanowells with protease (Qiagen) and DirectPCR Cell lysis reagent (Viagen). After overnight incubation cells are subjected to heat lysis and protease inactivation followed by tagmentation in a tagmentation mix (14.335 nL TD Buffer, 3.5 nL TDE1, and 0.165 nL 10% Tween-20) at 55°C for 10 minutes. Once the tagmentation reaction was neutralized, 8 cycles of PCR followed. The indexed single-cell libraries were recovered from the nanowells by centrifugation into a pool and sequenced on Illumina NovaSeq 6000.

### Immunofluorescence

#### Overview

We profiled matched FFPE tissues with cGAS and DAPI immunofluorescence to quantify the rate of micronuclei formation in tumors. The immunofluorescence detection of cGas was performed at the Molecular Cytology Core Facility of Memorial Sloan Kettering Cancer Center using Discovery XT processor (Ventana Medical Systems.Roche-AZ). Antigen retrieval was performed using ULTRA Cell Conditioning (Ventana Medical Systems, 950-224). The tissue sections were blocked first for 30 minutes in Background Blocking reagent (Innovex, catalog#: NB306).

#### Tissue staining

For the cGAS staining, a mouse monoclonal cGAS antibody (LSBio, LS-C757990) was used in 1:200 dilution. The incubation with the primary antibody is done for 5 hours followed by biotinylated mouse secondary (Vector Labs, MOM Kit BMK-2202) in 5.75μg/mL. Blocker D, Streptavidin-HRP and TSA Alexa594 (Life Tech, cat#B40957) was applied for 16 minutes.

All slides were counterstained in 5μg/mL DAPI (Sigma D9542) for 5 minutes at room temperature, mounted with anti-fade mounting medium Mowiol.

### RPE1 cell line experiments

We explored the phenotypic effects of chromosomal instability and WGD in *TP53*-knockout RPE1 cells. *TP53*-knockout RPE-1 was a gift from the Maciejowski laboratory at the Memorial Sloan Kettering Cancer Center (MSKCC). RPE-1 cells were cultured in DMEM (Corning) supplemented with 10% fetal bovine serum (Sigma-Aldrich), 1% penicillin-streptomycin (Thermo Fisher) at 37°C and 5% CO2. All cells were periodically tested for mycoplasma contamination.

*TP53*-/- RPE1 cells were treated with nocodazole, reversine and DMSO control to induce varying levels of chromosomal instability, then subject to both 10X multiome sequencing and DLP+ scWGS. For nocodazole treatment, RPE-1 cells were seeded at 20% confluence at the time of nocadazole addition. Cells were treated with 100 ng/ml nocodazole (Sigma-Aldrich) or DMSO for 8hrs. After 8hrs, cells were treated with three washes with phosphate buffered saline to remove the drug. After 48hrs the cells were collected. For reversine (Cayman Chemical Company) treatment, cells were treated at a concentration of 0.5 µM reversine for 48hrs. After 48hrs, cells were washed with three washes with phosphate buffered saline to remove the drug. Cells were collected after 12hrs. 10,000 cells per condition were collected for 10X Chromium Single Cell Multiome ATAC+Gene Expression according to the manufacturer’s protocol. Library preparation and sequencing were performed in MSKCC Integrated Genomics Core. 1M cells per condition were subject to DLP+ as described above.

A spontaneously arising WGD subclone was observed as a minor population of the *TP53*-knockout RPE1 cells (**Extended Data Fig. 6E**). After 30 additional passages (sample RPE-WGD), the WGD subclone, as measured by DLP+, comprised 37% of the population, presenting the opportunity to explore phenotypic differences between WGD and non-WGD cells. Sample RPE-WGD was subject to DLP+ scWGS and 10X scRNA.

| Sample | Treatment | Sequencing | Description |
| --- | --- | --- | --- |
| RPE-D | DMSO | 10X Multiome/DLP+ | non-WGD, low CIN control |
| RPE-Noco | Nocodazole | 10X Multiome/DLP+ | non-WGD, medium CIN |
| RPE-Rev | Reversine | 10X Multiome/DLP+ | non-WGD, high CIN |
| RPE-WGD | None | 10X scRNA/DLP+ | WGD/non-WGD mixed population |

## Computational methods

Computational analyses of multi-modal datasets were enabled by the Isabl platform^46^.

### Single-cell DNA sequencing

#### Overview

The single-cell DNA analysis pipeline is a suite of workflows for analyzing the single-cell data generated by the DLP+ platform^16^. The workflow takes dual-indexed reads from Illumina paired-end sequencing data as the input and performs various alignment and postprocessing tasks. The pipeline is publicly available on GitHub (https://github.com/mondrian-scwgs/mondrian), which we run within the Isabl framework^46^.

#### Alignment

We use Trim Galore to remove adapters and FastQC to generate QC reports before running alignment. The reads are then aligned with bwa-mem (with support for bwa-aln). The pipeline can also perform local indel realignment with GATK’s IndelAligner if required. PCR duplicates are marked using Picard MarkDuplicates and alignment metrics are computed for each cell with Picard tools CollectWgsMetrics and CollectInsertSizeMetrics. The pipeline also generates plots for each alignment metric for a quick overview.

#### Copy number segmentation

Reads are tabulated for non-overlapping 500 kb regions. A modal regression normalization^16^ is performed to reduce GC bias. The pipeline then runs HMMcopy with 6 different ploidy settings and the best fit is chosen automatically^47^. The pipeline also generates heatmaps with cell clustering, per-cell copy-number profile and the modal regression fit for visualization.

#### Quality control

scWGS was first subjected to quality control and filtering to remove non-cancer cells, S-phase replicating cells, low quality cells, and doublets, resulting in 29,481 high-quality cancer cell genomes (**Extended Data Fig. 2C-D**). The quality control pipeline compiles the results from the total copy number analysis and alignment, and we then use a random forest classifier to predict the quality of each cell based on the alignment and HMMcopy metrics^16^. We then inferred allele-specific copy number for each of these cells using `SIGNALS`^11^. Patient level average ploidy ranged from 1.6 to 4.5, and LOH ranged from 0.2 to 0.8. Ploidy and LOH estimates were concordant with matching bulk WGS and clinical panel sequencing by MSK-IMPACT, and losses and gains from scWGS coincided with known drivers of HGSOC (**Extended Data Fig. 2E-G**). Thus at a pseudobulk level, the genomic characteristics of our scWGS cohort matched those of both whole-genome and targeted bulk data.

#### Haplotype-specific copy number

In a matched normal sample we measure reference and alternate allele counts for SNPs from the 1000 Genomes Phase 2 reference panel. We use a binomial exact test to filter for SNPs heterozygous in the normal sample. Using SHAPEIT^48^ and the 1000 Genomes Phase 2 reference panel, we compute haplotype blocks. Next we measure per-cell reference and alternate allele counts for heterozygous SNPs in the tumor scWGS data.

#### Cell filtering

We established stringent filters to maximize the removal of problematic cells without sacrificing sensitivity to rare interesting populations including those representing cell specific WGD.

#### Removal of low-quality cells

We removed cells with quality score <0.75. The quality score was computed using the classifier presented in Laks et al., 2019^16^.

#### Removal of suspect high-ploidy cells

We restricted analysis to cells with high confidence ploidy calls. Absolute ploidy is unidentifiable from the sequencing data of an individual cell, thus we take a parsimony approach and assume the true ploidy to be the lowest ploidy value that provides a reasonable fit to the data. One failure mode in the automatic determination of ploidy by HMMCopy occurs when HMMCopy converges on a solution with double the true ploidy driven by the overfitting of isolated outlier bins. Such cells are characterized by mostly even copy number states except for isolated bins with odd copy number. To remove such potential artifacts we required there to be at least one segment >10MB in length with copy number 1, 3 or 5. Cells with no segment >10MB in length with copy number 1, 3, or 5 were removed from further analysis.

#### Removal of doublets

We applied several orthogonal approaches to remove doublets from the DLP data. First, under the assumption that chromosome 17 LOH should be clonal in ovarian cancer, we removed tumor cells that lacked LOH of chromosome 17. Then, we used a combination of mutation-based features to manually identify tumor-normal doublets, including LOH (much lower than typical tumor cells), proportion of SNVs with alternate reads (higher than typical normal cells), and copy-number profiles that were similar to tumor cells with the addition of 2 copies across the genome. Finally, 2 raters separately reviewed the brightfield image of each cell in the clear microfluidic nozzle prior to deposition in the microwell array for sequencing, and flagged any images that appeared to contain more than 1 cell. Any cell whose image was flagged by at least 1 reviewer was removed from analysis. Additional details on these approaches are described in the Supplementary Note.

#### Removal of S-phase cells

It is necessary to remove S-phase cells before downstream analysis as the observed HMMcopy profiles of these cells reflect a mixture of both somatic (heritable) copy number and transient doubling of replicated genomic loci. We nominated S-phase cells through a combination of features known to correlate with S-phase cells. As we aimed to isolate the high-quality G1/2-phase cells for downstream analysis, we did not need to distinguish between S-phase cells and low quality cells (i.e. noisy HMMcopy profiles due to other factors such as under-tagmentation prior to sequencing or incomplete cell lysis).

We first computed the following three features for each cell:

**1)** The Spearman correlation between HMMcopy state profile for a cell-of-interest and the RepliSeq replication timing profile from MCF-7 cells. S-phase cells will have higher correlations than G1/2- phase cells.
**2)** The number of HMMcopy breakpoints per cell (number of adjacent loci with different integer copy number state). S-phase cells have more breakpoints than G1/2-phase cells.
**3)** The median breakpoint prevalence across all HMMcopy breakpoints. This statistic is calculated by first computing the mean prevalence of each breakpoint across all cells belonging to said patient. Then, for each cell-of-interest, we subset to only the genomic loci with detected breakpoints in that cell and calculate the median of the mean breakpoint prevalences for said loci. S-phase cells have low median breakpoint frequency scores as they have lots of rare breakpoints.

All three features varied widely across patients due to each patient’s unique number, positioning, and heterogeneity of somatic copy number alteration. Thus we employed a strategy of examining each feature’s distribution across all cells in a patient, manually inspecting outlier cells, and selecting custom thresholds for each patient. We employ a filtering approach whereby cells are called as S-phase if any two of the three features are beyond the threshold. This conservative strategy ensures that all remaining cells are truly in G1/2-phase and therefore have HMMcopy profiles that accurately reflect somatic copy number.

#### Removal of normal cells

After copy number calling, we identified normal cells as those cells with copy number state average between 1.95 and 2.05 and standard deviation less than 0.5. We removed these normal cells from further analysis. We also manually inspected cells with aneuploidy slightly outside this range but much less than tumor cells in the same sample, and removed these “aberrant normal” cells (see Supplementary Note for examples). These cells typically did not share SNVs with the tumor cells and may correspond to other epithelial cells affected by field cancerization^49^ or immune/stromal cells with rare chromosomal aberrations.

#### Comparison with bulk copy number

We use WGS copy number inferred by *ReMixT*^50^ to validate the average ploidy in the MSK SPECTRUM cohort. Similarly, we use IMPACT copy number inferred by FACETS^51^ for additional orthogonal validation.

#### Detecting WGD in single cells using allele-specific copy number

WGD events were identified in single cells based on the allele-specific copy number state previously described for bulk WGS^18^. We computed two metrics from `SIGNALS` results: fraction of the genome with ≥2 copies for the major allele (*FM2*), and fraction of the genome ≥3 copies for the major allele (*FM3*). Similar to results in bulk WGS, a clear separation can be seen between subpopulations using each metric (**Extended Data Fig. 3H,I**). We classified any cell with FM2 > 0.5 as having undergone at least 1 WGD, and any cell with FM3 > 0.5 as having undergone at least 2 WGD.

#### Patient level WGD classifications

Patients were classified as Prevalent WGD if the fraction of cells classified as 1WGD≥1 exceeded 50% of the cells sequenced for that patient. The remaining patients were classified as Rare WGD.

#### Subclonal WGD classification

We classified cells within each patient as comprising a subclonal WGD subpopulation if they were predicted to have 1 more WGD than the number of WGDs for the majority population. However, for patients 081 and 125, a significant fraction of cells were predicted to be 0×WGD (>25%), with the remaining cells 1×WGD. For these patients, we considered the 1×WGD to be the subclonal WGD population.

#### Variant calling

##### SNV calling

Since the low per-cell coverage in scWGS is insufficient to resolve variants at nucleotide resolution, we merge all the single cells together to create a pseudo-bulk genome for each library. We run the Mutect2 variant caller^52^ on the merged data across all libraries from each patient. We compute the reference and alternate counts for each cell at variant loci that are detected by either caller over all libraries.

##### SV calling

We employed a similar approach for breakpoint calling by creating pseudo-bulk libraries, then running deStruct^53^ and Lumpy^54^ on each library. Only consensus SVs detected by both methods were retained, where an SV from both methods were considered consensus if their coordinates were within 200 bp and their orientations matched. The SV calls were further post-processed as described in a previous study^55^.

#### Filtering somatic variant calls using evolutionary constraints

Standard variant callers can produce artifactual calls on scWGS data, since its low insert sizes can result in incorrect alignments that appear to represent somatic variants. To address this, we developed a label propagation classifier to identify artifacts based on read-level features. To train this classifier, we leveraged the principle that distinct copy-number clones should not share subclonal variants to annotate high-confidence true and high-confidence artifact variants in a subset of samples. We then applied this classifier and trained it on high-confidence correct and artifactual calls based on manually labeled clones from a subset of patients, then applied it to all variants from all patients.

#### SBMClone

We applied SBMClone^20^ to the filtered somatic variants for each patient. SBMClone was run 10 times on each patient with different random initializations, and the solution with the highest likelihood was kept.

#### Evolutionary histories of SNV clones using doubleTime

We developed doubleTime, a method for computing evolutionary histories of the SNV clones in each patient, including accurate placement of WGD events in the clonal phylogeny of each patient. Our approach involved three major steps. First, we constructed a clonal phylogeny relating the clones identified by SBMClone. Second, we assigned WGD events to branches in the clonal phylogeny. For each pair of WGD clones, we assessed whether those clones arised from a single common WGD or two independent WGD. Given this information we were able to unambiguously assign WGD events to branches throughout each patient’s clonal phylogeny. Third, we used a probabilistic model to assign SNVs to branches of the clonal phylogeny, including assignment before and after WGD events on WGD branches. We describe each of the three steps in additional detail below.

#### Perfect phylogenies of SNV clones

We reconstructed phylogenetic trees with SBMClone clones as leaves using a binarized version of the implicit block structure inferred by SBMClone. We first computed a density matrix D where each row corresponds to a clone (i.e., cell block), each column corresponds to an SNV cluster (i.e., SNV block) and each entry *D*_*i,j*_ contains the number of pairs *(a,b)* in which cell *a* in clone *i* has at least one alternate read covering SNV *b* in cluster *j*, divided by the total number of possible pairs (i.e., the size of clone *i* times the size of cluster *j*). We then computed a binary matrix B by rounding up those entries of D that exceeded a density of 0.01, removed empty columns, and attempted to infer a phylogenetic tree by applying the perfect phylogeny algorithm. Matrices B that did not permit a perfect phylogeny were manually modified with the minimum number of changes required to permit a perfect phylogeny – this typically occurred when mutations shared between two or more clones had been lost due to a deletion in a subset of the clones.

#### Discerning independent from shared WGD

To identify cases in which sequenced WGD cells arose from distinct WGD events, we analyzed SNVs from the single-cell DNA sequencing data. Specifically, for each patient, we focused exclusively on those regions that exhibited copy-neutral loss of heterozygosity (cnLOH; i.e., major copy number 2 and minor copy number 0) among nearly all (≥ 90%) tumor cells with a single WGD. Given a candidate bipartition of the 1 WGD cells, under the infinite sites assumption, each cnLOH SNV then fits into one of the following categories:

- 2 mutant copies in both clones (shared pre-WGD and pre-divergence)
- 1 mutant copy in one clone (private post-divergence)
- 0 mutant copies (false positive variant)
- 1 mutant copy in both clones (shared post-WGD and pre-divergence)
- 2 mutant copies in both clones (post-WGD and post-divergence)

The last two categories of SNVs present evidence for or against multiple independent WGD events. SNVs that are shared at 1 variant copy (VAF ~0.5) would suggest that the two sets of cells underwent the same ancestral WGD event, as they share mutations that must have followed the WGD. Conversely, SNVs that are private at two variant copies (VAF ~1) would suggest that the two sets of cells underwent distinct WGD events, as they have private mutations that preceded the WGD. Specifically, we considered the following hypotheses:

1. Single-WGD: Shared 1-copy SNVs are allowed, but private 2-copy SNVs are not allowed.
2. Multiple-WGD: Shared 1-copy SNVs are not allowed, but private 2-copy SNVs are allowed.

To evaluate the relative strength of these hypotheses, we developed a likelihood ratio test that compared the probability of observing the given variant counts for cnLOH SNVs under these two hypotheses: for each patient, we evaluated P(Multiple-WGD)/P(Single-WGD) using a simple binomial model of read counts. We then tested the significance of this likelihood ratio by generating an empirical distribution: we fixed the SNV read counts and their best-fitting variant copy numbers under the Single-WGD hypothesis and resampled alternate counts.

#### Assigning SNVs to branches and estimating branch lengths

From the previous steps, we are given a tree relating the clones detected by SBMclone. We place WGD events on branches such that all Prevalent WGD patients had a WGD event placed on the root of the tree, except those in which independent WGD events had been identified (patients 025 and 045) or WGD only affected a subset of clones (patient 081), in which case those specific events were placed further down the tree. We used a probabilistic model to assign SNVs to branches and estimate branch lengths based on read count evidence for SNVs in each clone. For WGD branches, the model assigns SNVs as occurring before or after the WGD, and estimates the length of the branch before and after the WGD. This strategy effectively splits each branch with a WGD event into two unique positions in the tree, meaning that the total number of positions in the tree to which an SNV can be assigned is equal to the number of branches determined by SBMclone + the number of branches with WGD events.

For this analysis, we considered only those SNVs in regions where for each SBMClone clone, over 80% of cells shared the same copy-number state. We further restricted analysis to SNVs in regions with allele-specific copy-number states whose multiplicity (i.e., variant copy number, or the number of copies of the genome containing the SNV) and thus expected VAF could be uniquely determined by the combination of tree placement and WGD status (i.e., whether or not the clone was affected by an ancestral WGD event). Specifically, we analyzed regions with the following copy-number states across all clones:

- 1:0 in both WGD and non-WGD clones
- 1:1 in both WGD and non-WGD clones
- 2:0 in WGD clones, 1:0 in non-WGD clones
- 2:1 in WGD clones, 1:1 in non-WGD clones
- 2:2 in WGD clones, 1:1 in non-WGD clones

In each of these scenarios, we assume that the WGD and copy-number events immediately following the WGD account for the differences in copy number between WGD and non-WGD clones. Note that the only patient in the cohort with different WGD status for different leaves was patient 081, so for nearly all patients we analyzed only those SNVs with clonal copy-number states (matching the above listed states depending on WGD status). The multiplicity for an SNV on a particular allele placed on a particular branch of the tree was as follows:

- 0, if the corresponding allele had 0 copies
- Equal to the allele-specific copy number of the allele in the clone, if the SNV occurred pre-WGD and the leaf was affected by WGD
- Equal to 1 otherwise

Each SNV is assigned to a tree position by fitting the observed total and alternative counts of said SNV to the expected VAFs for all clones. SNVs are assigned to positions in the tree using a Dirichlet-Categorical distribution, and a Beta-Binomial emission model is used to relate observed SNV counts to expected VAFs. The model is implemented in Pyro and fit using black box variational inference^56^. Note that when computing branch lengths, we only use C>T SNVs at CpG sites as these SNVs have been reported to correspond most closely to chronological age^57^.

To account for the differences in genome size and copy-number heterogeneity between different patients with varying amounts of aneuploidy, we normalize the number of C>T CpG SNVs on each branch by the number of bases being considered. First, we computed the effective genome length of each clone as the total size of the bins considered to be “clonal” for a valid copy-number state as defined above, with each bin weighted by its total copy number. Then, for the internal nodes of the tree, we assumed that the only copy-number changes to these bins were directly coupled to WGD events. Thus, for post-WGD branches, the genome length was identical to that of the leaves; and for pre-WGD branches, the genome length was computed using the correspondence described above between pre- and post-WGD copy numbers.

#### Estimating pre- and post-WGD changes in WGD subpopulations

We use a maximum parsimony based method to estimate pre- and post-WGD changes from estimated ancestral and descendent copy-number profiles. We proceed independently for each bin. Let *x* be the ancestral copy number state and *y* the descendent copy number state, and assume *y* is produced by a combination of pre-WGD CN change followed by WGD followed by post-WGD CN change. We can relate *x* and *y* using,

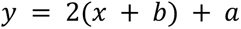

where *b* represents pre-WGD CN change and a post-WGD CN change. Let the *cost* of any given *a* and *b* be |*a*| + |*b*|. Conveniently, every combination of *x* and *y* results in a unique *a* and *b* that minimize this cost. Thus, for each *x* and *y* we compute the associated *b* and *a* as the pre- and post-WGD changes and |*a*| + |*b*| as the cost of those changes.

#### Measures of diversity and heterogeneity

We compute the “percent genome different” for a pair of cells as follows. First, we compute the bin level difference in total copy number and identify consecutive segments of changed and unchanged bins. We then remove segments less than or equal to 2 MB in size (i.e., affecting fewer than four consecutive 500-kb bins). Finally, we count the number of bins for which the two genomes have different total copy number and divide by the total number of bins considered.

#### Classification of divergent cells

We call divergent cells as outliers of the nearest neighbor distance (NND), using “percent genome different” as the distance metric. For each index cell we compute its nearest neighbor as the other cell in the population within minimum percent genome different. The nearest neighbor distance for each cell is thus the percent genome different with respect to its neighbor cell. We then fit a beta distribution to the NND values of all cells in the cohort, and call divergent cells as those cells that have NND values in the 99th percentile of the beta distribution fit to the data.

#### Cell phylogenies using MEDICC2

We derived estimates of chromosome missegregation rates per cell for each patient from copy-number phylogenies inferred with MEDICC2^58^. First, we refined single-cell haplotype-specific copy-number profiles by applying the dynamic programming formulation from asmultipcf^59^ on GC-corrected read counts and phased B-allele frequencies for each bin. Using this method, we identified segment boundaries across all cells for each patient and then summarized the number of copies of each segment and allele in each cell by rounding. Next, we ran MEDICC2^58^ on these refined haplotype-specific single-cell copy numbers, which infers a tree (with single cells corresponding to leaves), copy-number profiles for the ancestral internal nodes of the tree, and copy-number events for each branch of the tree. We used the –wgd-x2 flag for MEDICC2 which represents WGD as an actual doubling of all copy-number segments in the genome, rather than the default behavior of adding 1 to all segments. We then computed missegregation rates by counting the number of inferred chromosome-/arm-level gains and losses on the terminal branches of the tree (i.e., the number of cell-specific events) and dividing by the total number of cells in the tree.

#### Reconstruction of ancestral copy number

To infer the ancestral haplotype-specific copy-number profiles associated with internal nodes of the cell phylogeny, we use a maximum parsimony approach that treats each bin independently and aims to minimize the total number of changes on the tree. Specifically, the parsimony score for each branch is the sum across bins and across both haplotypes of the absolute difference in copy number between the parent and the child. Transitions from 0 to any other copy number are given a score of infinity to prevent gain from 0 copies. The score for a WGD branch (assumed known from MEDICC2) is the sum of two parsimony scores: the score for copy-number changes between the parent and an intermediate genome, and the score for copy-number changes between a doubled version of the intermediate genome and the child (this intermediate genome is described above in *Estimating pre- and post-WGD changes in WGD subpopulations*). The state of each bin at each branch in the tree is chosen to minimize this parsimony score using the Sankoff algorithm^60,61^. We assume that the MEDICC2 placement of WGD on branches of the phylogeny is correct in all but two patients: for patients 025 and 045, we adjusted WGD placement to be concordant with SNV evidence suggesting independent clonal origin of multiple WGD clones.

#### Classifying event from CN differences

Given a phylogenetic tree where both leaves and internal nodes are labeled by haplotype-specific copy-number profiles, we identify the copy-number events on each branch using a greedy approach. First, we identify the differences between the parent haplotype-specific copy-number profile and the child copy-number profile. Then, for each chromosome and haplotype, we aim to explain the copy-number differences between parent and child using events that are as large as possible:

1. If more than 90% of bins in the chromosome are altered in the same direction, we call a chromosome gain or loss that accounts for a change of one copy for all bins in the chromosome.
2. If no chromosome gain or loss is found, but 90% of the bins in one of the two arms is altered in the same direction, we call an arm-level gain or loss that accounts for a change of 1 copy for all bins in the chromosome arm.
3. If no chromosome- or arm-level gain or loss is found, we call a gain or loss of the largest contiguous segment that has a change in the same direction.

We then adjust the copy number difference by the selected event, and repeat until all copy-number changes between parent and child have been accounted for. Note that if all but a few of the bins of a chromosome are gained (or lost), our method will first predict a chromosome gain (or loss), then an additional small segment loss (or gain) to account for the few bins that were predicted as unchanged. We have selected this approach as we consider a whole chromosome (or arm) change to be more parsimonious if most of a chromosome’s (or arm’s) bins are altered. Our approach is also more robust to bin level noise than a strategy that requires 100% of the bins to be altered.

For branches with WGD, we compute the intermediate pre-doubling profile that would result in the fewest copy-number changes (see *Estimating pre- and post-WGD changes in WGD subpopulations* above). Using our bin-independent parsimony model, we can compute the optimal intermediate profile analytically. We then perform the event calling procedure described above twice: once on the differences between the parent and the intermediate pre-WGD profile, and again between the doubled intermediate profile and the child.

#### Estimating rates of cell specific events

We explored controlling for the “opportunity” for each cell to mis-segregate by dividing the number of copy-number events for each cell by the number of chromosomes (for chromosome-level missegregations) or arms (for arm-level missegregations) in the inferred parent node of each cell in the tree (i.e., the source of the terminal branch). This yields a rate of missegregation events per cell and per parental copy. For shorter “segmental” copy-number events, we divided the number of events in each cell by its parent’s genome length to control for opportunity. While the resulting rate is not comparable to segment- and arm-level rates, it makes the cell-specific segmental rates more comparable between cells and across patients.

#### Detection of focal high-level amplifications in single cells

To detect focal high-level amplification in single cells, we used a two-stage approach compiling a set of potential amplified segments, then re-called amplification of those segments in individual cells. We first identified all contiguous segments with copy number exceeding 3✕ ploidy per cell. We then merged per cell segments to generate a set of amplified segments for the patient tumor cell population as a whole, and merged adjacent amplified segments if the boundaries of those segments were closer than 2MB. Only amplified segments larger than 500kb (1 bin) were considered further. Given a set of amplification segments predicted per patient, we then computed the average copy number for each cell within each segment, as well as the average copy number for the 5MB on either side of each segment. A focal high-level amplification was called in an individual cell if the average copy number of the amplification segment was greater than 3✕ ploidy and greater than 3✕ the average copy number in the boundary segments.

#### Enumerating events on ancestral branches

We computed gains and losses of chromosomes and chromosome arms for three classes of event timing. Events were classified as non-WGD if they were predicted to occur on the root branch of a Rare WGD patient, pre-WGD if they were predicted to occur prior to the WGD event on the root branch of a Prevalent WGD patient, and post-WGD if they were predicted to occur prior to the WGD event on the root branch of a Prevalent WGD patient. Patients 025, 045, and 081 were omitted from this analysis as their WGD history precludes this categorization of copy-number events.

#### Calculating post-WGD changes in WGD clones

We cataloged all high confidence WGD clones detected in our cohort. This included all predicted WGD clades with at least 20 cells in the MEDICC2 phylogenies. In addition, we included two small WGD clones from patient 006 and 031 (**Extended Data Fig. 3E-F**). Counts of post-WGD events were calculated from ancestral reconstruction on MEDICC2 trees as described above (see section *Reconstruction of ancestral copy number*).

### Single-cell RNA sequencing

#### Cell type assignment

Using scRNA-seq of CD45^+/-^ sorted cells we assigned major cell types using supervised clustering with CellAssign^62^, as described in Vázquez-García et al., 2022^14^.

#### InferCNV and scRNA-seq derived copy number clonal decomposition

InferCNV (version 1.3.5) was used for identifying large-scale copy number alterations in ovarian cancer cells identified by CellAssign^63,64^. For each patient, 3,200 non-cancer cells annotated by CellAssign were randomly sampled from the cohort and used as the set of reference “normal” cells. After subtracting out reference expressions in non-cancer cells, chromosome-level smoothing, and de-noising, we derived a processed expression matrix which represents copy number signals. Cancer cell subclusters are identified by ward.D2 hierarchical clustering and “random_trees” partition method using *p*-value < 0.05.

#### WGD classification

Identification of WGD cells from scRNA data is technically challenging, as inferred copy number from expression data is typically noisy, allele-specific markers are sparse, and as shown in our scWGS analysis, the prevalence of non WGD cells in Prevalent WGD cases, and WGD cells in Rare WGD cases is generally low, confounding identification of non-clonal ploidy populations within samples. We reasoned that due to the high concordance between scWGS and scRNA derived copy number, even between non site-matched patient samples (**Extended Data Fig. 5A**), and the typically clonal nature of WGD, WGD status could be propagated to all available patient matched scRNA samples for the purposes of transcriptional phenotyping analysis. Furthermore, within-sample absolute normalization of UMI counts between tumor and non-tumor cells showed a significant increase in overall transcript counts per cell in Prevalent versus Rare WGD patients (**Extended Data Fig. 5B**), which was highly concordant with established estimates of transcriptional changes in WGD versus non-WGD samples in bulk RNA^65^. Thus, we concluded that site-matched scRNA data effectively captures WGD transcriptional phenotypes. Any analyses correlating scWGS derived missegregation rates to transcriptional phenotypes were restricted to site matched samples with at least 20 cells in both DLP and scRNA.

#### Cell cycle analysis

We identified circular trajectories linked to cell cycle progression in cancer cells using Cyclum^28^. Across the cohort, 10,000 cancer cells annotated by CellAssign were randomly sampled across tumors and used for cell cycle trajectory inference. Pseudotime inference was run on the scaled cell-by-gene matrix, limiting genes to cell cycle markers included in cell cycle GO terms (GO:0007049). Discretization of the continuous pseudotime trajectories was accomplished using a three-component Gaussian mixture model. Discrete cell cycle phase information was computed using Seurat’s CellCycleScoring function, excluding samples with fewer than 20 malignant cells. Smoothed pseudotime trajectories of cell cycle-related genes previously reported in the literature^66^ were then evaluated to interpret phase-specific gene activity and phase transitions as a function of pseudotime (**Fig. 5D**).

#### Differential gene and pathway activity

Pathways were curated from the single-cell hallmark metaprograms^67^, the 50 hallmark pathways^68^, or CIN-associated gene signatures manually curated from literature, including inflammatory signaling and ER stress^31,38^, and scored in single cells using Seurat’s ‘AddModuleScore’ function. Due to the hierarchical nature of the data, with multiple samples from patients, we used generalized estimating equations (GEE) on sample mean gene or pathway expression levels, adding tumor site (adnexa vs non-adnexa) as a covariate in the model, and restricting analysis to samples with at least 20 cells in order to compare WGD states. *P*-values were adjusted for multiple testing using FDR. In parallel, we also performed differential expression analysis using a pseudobulked generalized linear mixed model (DREAMLET^69^), accounting for random patient and fixed tumor site effects, and performed gene set enrichment analysis (GSEA) with the same set of pathways.

#### Differential cell type abundance

To determine cell populations that were differentially abundant between rare WGD and prevalent WGD samples we utilized miloR v1.8.1^70^, setting ‘prop’ to 0.2, and using ‘tumor_megasite’ (adnexa vs non-adnexa) as a contrast in the differential abundance testing. To obtain significance values for each cell population, we ran permutation tests by swapping the sample WGD status labels 1,000 times, and computing the proportion of tests in which the resulting non-permuted median log2-fold change was more extreme than the permuted median values for each cell type.

### Immunofluorescence

#### Regions of interest

We defined regions of interest (ROIs) containing tumor on IF images by delineating regions with tumor foci, and contrasting these with images of the IF-adjacent H&E section. ROI annotations were drawn in QuPath. To ensure that complex tissue regions within ROIs used for analysis only included tumor, we classified regions of tumor, stroma, vasculature and glass within each ROI. We trained a pixel classifier with examples of tumor, stroma, vasculature and glass from each of the ROIs and slides using the IF-adjacent H&E section.

#### Segmentation of primary nuclei and micronuclei

Whole-slide IF images stained with cGAS, ENPP1 and DAPI were analyzed to characterize primary nuclei (PN) and micronuclei (MN) within ROIs. Segmentation of PN was carried out in QuPath v0.3.0 using the StarDist algorithm on the DAPI channel^71^. We used a segmentation model pre-trained on single-channel DAPI images (“dsb2018_heavy_augment.pb”). Applying the PN segmentation model across all ROIs yielded 1,779,351 PN in tumor regions. Segmented PN ranged between 5 μm^2^ and 100 μm^2^ in size, with a minimum fluorescence intensity of 1 a.u. The cell membrane for each PN was approximated using a cell expansion of 3 μm of the nuclear boundary.

Micronuclei were detected by StarDist segmentation of cGAS spots. We trained a new segmentation model on single-channel cGAS images using a U-Net architecture. We manually annotated cGAS^+^ MN in a set of 256px x 256px tiles encompassing tumor regions across all slides. We created training and test sets using a 70:30 split, resulting in a training set of 70 tiles and a test set of 30 tiles. To ensure that the model generalized across patients and samples, we applied augmentation to the training set by applying random rotations, flips, and intensity changes. We monitored the loss function during model training and saved the trained model with frozen weights.

This allows for whole slide quantification and cell-level annotation of PN and MN. Nuclear segmentation was also carried out using StarDist on the DAPI channel. Each MN was assigned to the closest PN. MN were excluded if they were >10μm from the centroid of the closest nucleus, had area >10μm^2^ or probability <0.75.

#### Validation of micronuclei segmentation

We have evaluated our method on a test dataset with held-out MN labels, showing good performance of predicted MN segmentations with high average precision and F1 scores (IoU < 0.5). We quantitatively evaluated the segmentation performance on the test data by considering cGAS^+^ MN objects in the ground truth to be correctly matched if there are predicted objects with overlap. We used the intersection-over-union (IoU) as an overlap criterion, demonstrating good performance with a chosen IoU threshold > 0.5.

#### Micronuclei rates

Micronucleus rupture rates were estimated based on the number of cGAS^+^ MN and PN segmented within tumor ROIs. The rate of micronuclei rupture was estimated by localization of cGAS^+^ MN neighboring PN. MN rate was calculated as the fraction of PN with 1 or more MN. Applying the MN segmentation model across all ROIs yielded 83,352 cGAS^+^ MN in tumor regions, with a mean MN area of 2 μm^2^, ranging between 1 μm^2^ and 10 μm^2^, and a minimum object probability of 0.75. To overcome batch effects, we used within-batch MN rate Z-score for downstream comparisons.

#### Statistical comparisons of micronuclei rates

For comparing MN rate between prevalent and rare WGD, we used generalized estimating equations (GEE). We used binary Prevalent vs Rare WGD as the dependent variable with binomial distribution and Z-score MN rate as the independent variable, adding patient as a group variable in the model. Reported effect size of WGD was calculated from the coefficient of Z-score MN rate in the learned model. For correlation between gain and loss rates and MN rate, we used a mixed linear model with Z-score MN rate as the dependent variable, gain or loss rate as the independent variable, and patient as a group variable.

### Multi-modal sample matching

For integrative genotype-phenotype analyses, we utilized scRNA-seq data patient-matched with scWGS to profile cell type-specific abundance and gene/pathway activity changes in the context of WGD (**Figure 6**). Given the clonally dominant nature of each sample’s WGD status, we reasoned that tumor cells identified in scRNA-seq within each patient would likewise be mostly clonal WGD or not, allowing for direct comparisons across all tumor cells in each patient. Indeed, site-matched scWGS and scRNA-seq derived estimates of copy number were highly concordant (**Extended Data Fig. 6A**), with UMI count ratios between tumor and normal cells being significantly elevated in Prevalent WGD compared to Rare WGD cases as expected (**Extended Data Fig. 6B**).

### Mutational signatures

We analyzed mutational signatures by integrating SNVs and structural variations detected by either bulk WGS or scWGS in a unified probabilistic approach called multi-modal correlated topic models (MMCTM)^15^.

For bulk WGS samples, we obtained signature labels in the MSK SPECTRUM cohort (*n*=40) using MMCTM, as presented in Vázquez-García et al., 2022^14^. Mutational signatures for cases without bulk WGS data were assigned based on mutational signatures inferred from scWGS. For scWGS samples, we obtained signature labels in the MSK SPECTRUM cohort (*n*=40) using a ridge classifier with default regularization strength (α=1.0). This classifier was trained on the integrated SNV and SV signature probabilities, which were obtained using MMCTM^11^ from HGSOC bulk whole genomes (*n*=170)^11^.

Consensus mutational signatures were preferentially derived based on: (i) MMCTM signatures derived from bulk WGS, and (ii) MMCTM signatures from scWGS. Mutational signatures for cases without bulk WGS data (006, 044, 046, 071) or inconclusive bulk WGS assignments (004, 045, 080, 081) were resolved based on scWGS.

### Analysis of RPE1 cell line experiments

#### 10X scRNA pre-processing

Raw 10X sequencing data were aligned using CellRanger (version 7.0.0), which also performed barcode filtering and unique molecular identifier (UMI) gene counting using the 10X GRCh38 reference transcriptome.

#### 10X Multiome pre-processing

Raw 10X sequencing data were aligned to the 10X GRCh38 reference transcriptome using CellRanger ARC (version 2.0.2). CellRanger ARC also performed barcode filtering and unique molecular identifier (UMI) gene counting to generate feature-barcode matrices for both RNA and ATAC modalities.

#### scATAC copy number analysis

Copy number was inferred from the scATAC component of the 10X multiome data for RPE-D, RPE-Noco and RPE-Rev samples. Blacklist filtered fragments were first counted in 10MB genome bins. Bins with GC content of less than 30% were removed prior to performing GC correction using modal regression^16^. Cells with more than 5% of their bins containing NA values after GC modal correction were removed from subsequent analysis. GC corrected counts were smoothed using the DNACopy R package (v1.73.0) ‘smooth.CNA’ function, setting ‘smooth.region’=4. Smoothed counts were mean-normalized per cell prior to clustering using Seurat (v5)^72^. For visualization, mean-normalized and smoothed counts were scaled binwise to emphasize copy differences between clusters.

#### scRNA copy number analysis

Copy number was inferred from 10X scRNA for the RPE-WGD sample using Numbat (v1.4.0)^73^ to preprocess and smooth expression counts. Smoothed counts were then rebinned to 500Kb, bins, reduced to 50 dimensions by PCA, and then clustered using Leiden clustering at 1.0 resolution on a SNN graph.

#### Identification of WGD subclones

A spontaneously arising WGD subclone was observed in all DLP+ samples, characterized by gain of 1p and loss of 1q, 2q, 4q and 21 (**Extended Data Fig. 6E**). The same WGD clone was evident copy number inferred from both scATAC for RPE-D, RPE-Noco and RPE-Rev (**Extended Data Fig. 6D**) and scRNA for RPE-WGD (**Extended Data Fig. 6I**). For RPE-D, RPE-Noco and RPE-Rev, our aim was to characterize the phenotypic impact of CIN in non-WGD cells. Thus we excluded scRNA cells in the scATAC inferred WGD cluster from further analysis. For RPE-WGD we aimed to characterize the phenotypic differences between WGD and non-WGD cells. We thus used the scRNA based copy number clusters to label cells as either WGD or non-WGD in that sample.

#### Estimating rates of cell specific events from DLP+

We inferred cell-specific rates of copy number change from DLP+ data using similar methods to those applied to the patient data. We first removed low quality and cycling cells as described above. For RPE-D, RPE-Noco and RPE-Rev we removed cells with ploidy > 2.5, thereby removing the WGD clone and other WGD cells. We then used MEDICC2 to infer a phylogeny independently for each sample, computed cell specific changes and classified those changes into chromosome, arm, and segment as described above.

## FIGURES

**Extended Data Figure 1.**
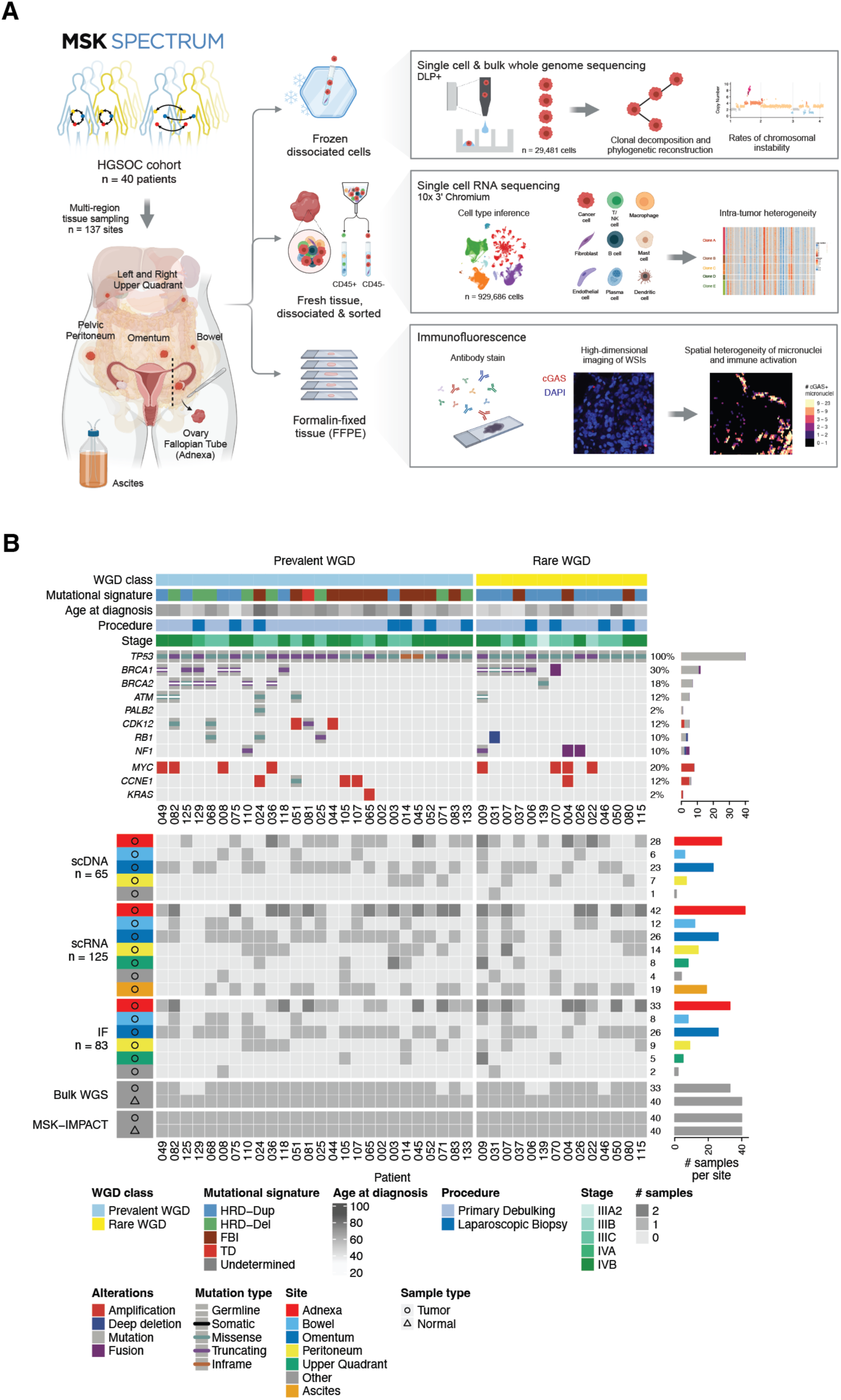
Study and cohort overview. **A.** Schematic of the MSK SPECTRUM specimen collection workflow including primary debulking surgery or laparoscopic biopsy, single-cell suspensions for scWGS and scRNA-seq, and biobanking of snap-frozen and FFPE tissue samples. **B.** Cohort overview. Top panel: Oncoprint of selected somatic and germline mutations per patient and cohort-wide prevalence. Single nucleotide variants (SNVs), indels, and fusions shown are detected by targeted panel sequencing (MSK-IMPACT). Focal amplifications and deletions are detected by single-cell whole genome sequencing (scWGS). Patient data include WGD class, mutational signature subtype, patient age, staging following FIGO Ovarian Cancer Staging guidelines, and type of surgical procedure. Bottom panel: Sample and data inventory indicating number of co-registered multi-site datasets: single-cell whole genome sequencing, single-cell RNA sequencing, H&E whole-slide images, immunofluorescence, bulk WGS and bulk MSK-IMPACT.

**Extended Data Figure 2.**
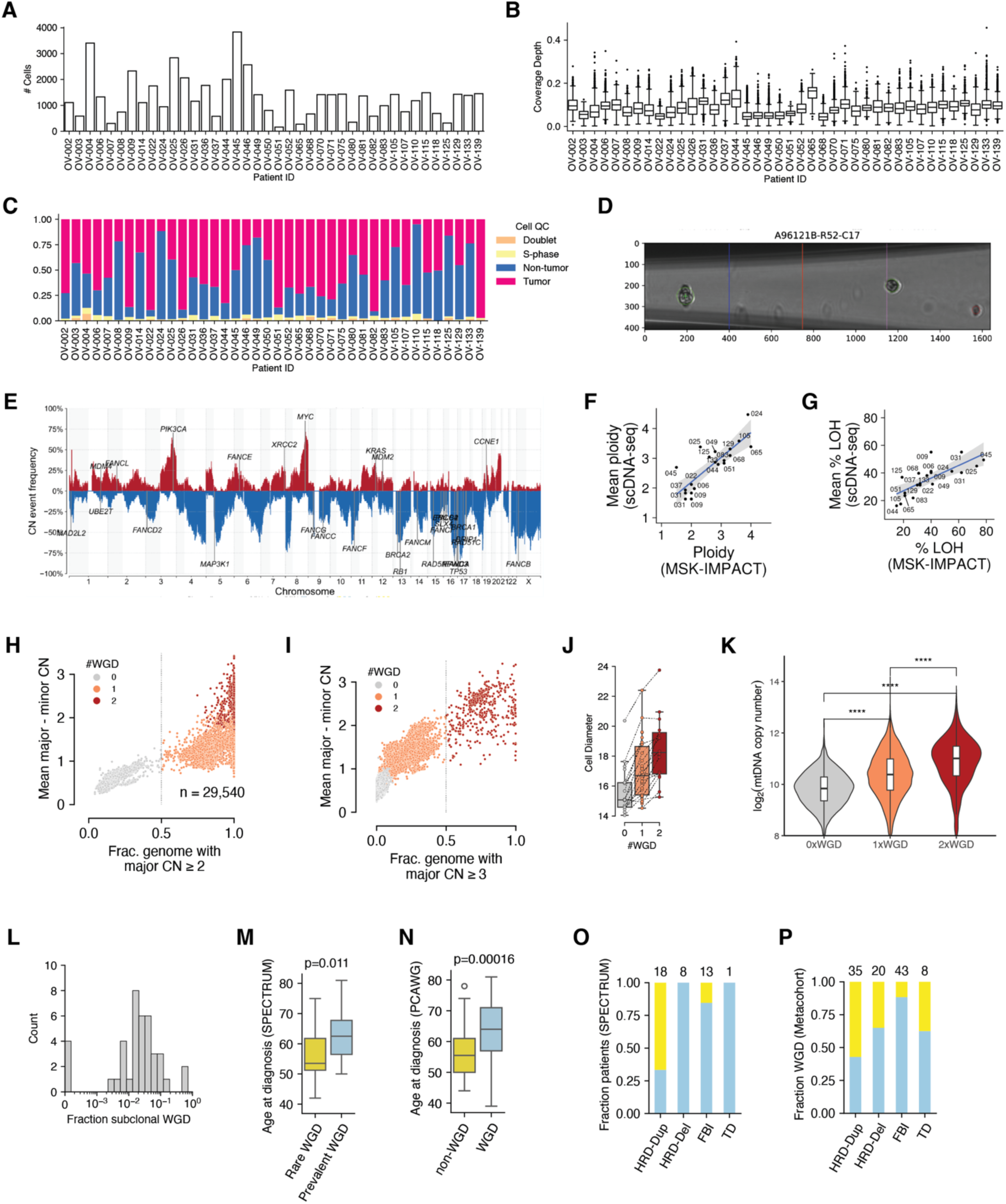
Quality control of scWGS data and WGD inference. **A.** Number of high-quality cells generated per patient. **B.** Distributions of per-cell coverage depth per patient. **C.** Fraction of cells called as tumor, non-tumor, doublet, and S-phase for each patient. **D.** Example doublet identified from an image taken during DLP+ sequencing. **E.** Frequency of gains (red, above the horizontal) and losses (blue, below the horizontal) among all single-cell genomes in the cohort, with known drivers genes annotated. **F.** Ploidy (mean copy number) for each patient in the SPECTRUM cohort as measured by MSK IMPACT (x-axis) and scWGS (y-axis). **G.** Fraction of the genome with loss of heterozygosity (LOH) for each patient in the SPECTRUM cohort as measured by MSK IMPACT (x-axis) and scWGS (y-axis). **H.** Shown for all quality-filtered cells in the cohort is the mean difference between major and minor copy number (y-axis) versus the fraction of the genome with major copy number ≥ 2 (x-axis), with cells colored by #WGD state. The dashed line at 0.5 denotes the decision boundary for 0 vs 1 WGDs. **I.** Shown for all quality filtered cells in the cohort is the mean difference between major and minor copy number (y-axis) versus the fraction of the genome with major copy number ≥ 3 (x-axis), with cells colored by #WGD state. The dashed line at 0.5 denotes the decision boundary for 1 vs 2 WGDs. **J.** Cell diameter measured from DLP+ images. Each point is the mean cell diameter within a given patient for 0×, 1× or 2×WGD cells. Points representing cells from the same patient are connected by dashed lines. Boxplots show the distribution of means for each WGD state. **K.** Distribution of mitochondrial DNA copy number (log2) inferred from scWGS in 0×, 1×, and 2×WGD cells. **L.** Distribution over patients of the fraction of cells within each patient with subclonal WGD, i.e., 1 more WGD than the dominant population for that patient. **M.** Age at diagnosis for patients in the SPECTRUM cohort split by Prevalent vs Rare WGD. **N.** Age at diagnosis for patients in the PCAWG ovarian cohort split by WGD vs non-WGD. **O.** Fraction of Prevalent and Rare WGD patients in the SPECTRUM cohort for each mutational signature. **P.** Fraction non-WGD and WGD patients in the Ovarian Metacohort for each mutation signature.

**Extended Data Figure 3.**
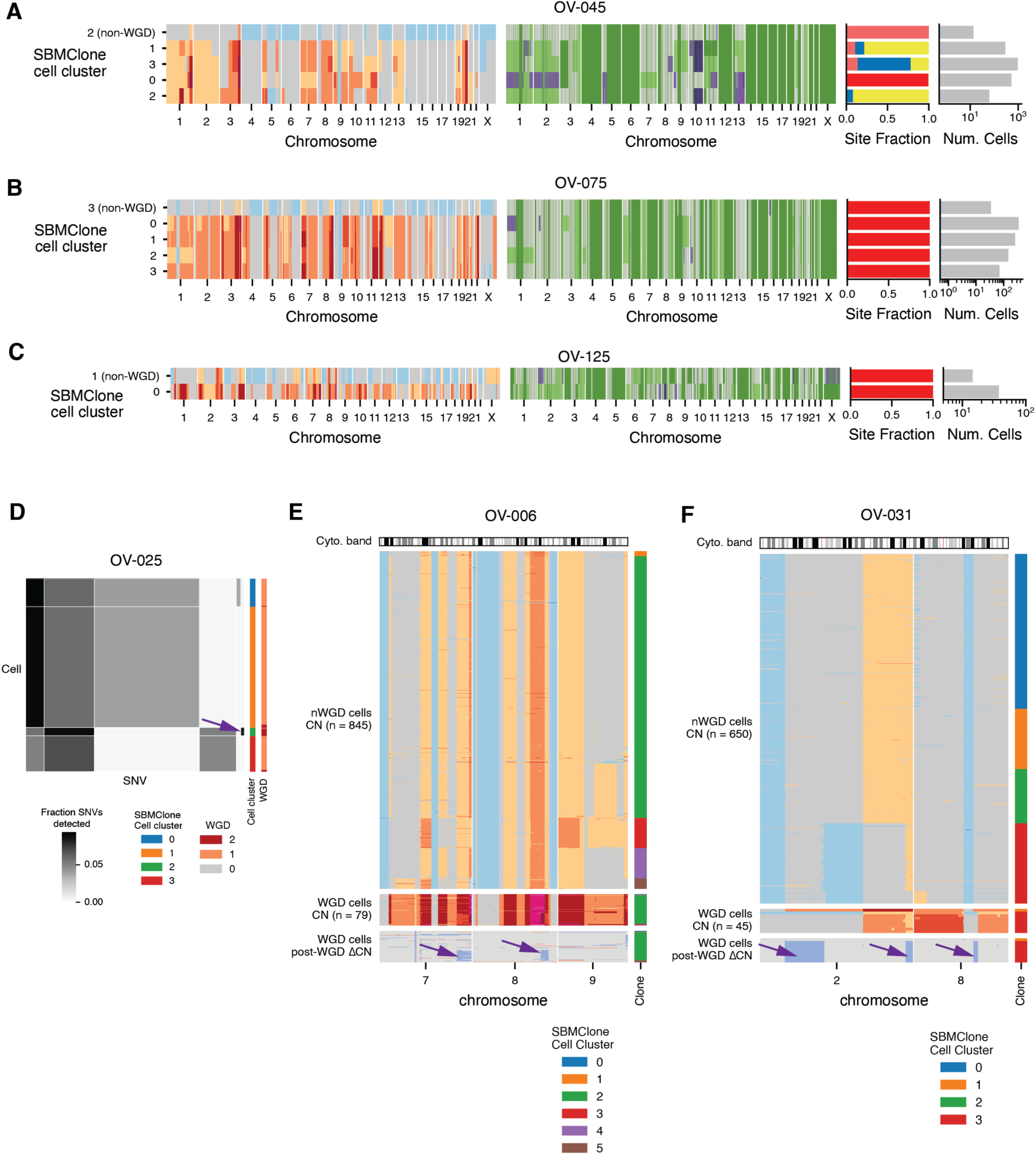
Non-WGD subclones and subclonal WGD. **A-C.** 0×WGD subpopulations in patients 045 (A), 075 (B) and 125 (C). Shown for each patient is the total (left) and allele specific (middle) copy number for each clone (y-axis). At right are the fraction of cells from that clone found in each anatomic site (left) and the number of cells for each clone (right). **D.** SBMClone block density matrix for patient 025 showing the proportion of SNVs detected for each clone (y-axis) and SNV block (x-axis). The SBMClone cluster and WGD status of each cell are shown on the right. The 2×WGD clone in patient 025 is distinguished by clone-specific SNVs (arrow). **E.** Copy number for chromosomes 7, 8, and 9 for cells in patient 006, separated into non-WGD cells (top), WGD cells (middle), and inferred post-WGD changes in WGD cells (bottom). The cell order is the same for the middle and bottom plots. Arrows indicate shared post-WGD changes that represent a WGD subclone. **F.** Copy number for chromosomes 2 and 8 for cells in patient 031, separated into non-WGD cells (top), WGD cells (middle), and inferred post-WGD changes in WGD cells (bottom). The cell order is the same for the middle and bottom plots. Arrows indicate shared post-WGD changes that represent a WGD subclone..

**Extended Data Figure 4.**
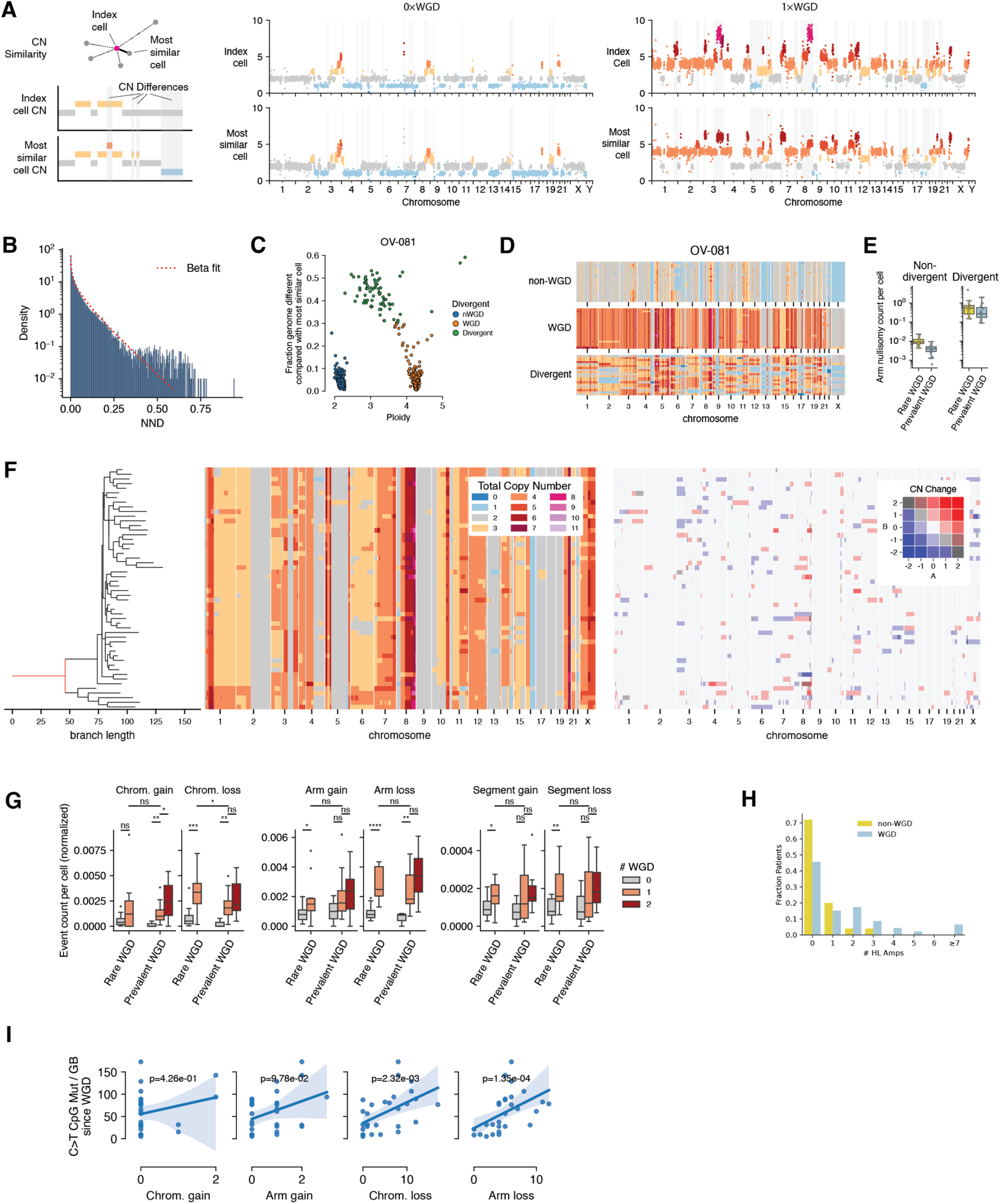
Single cell measurement of chromosomal instability. **A.** Schematic of nearest neighbor difference (NND) using fraction of the genome different as a distance measure (left). Shown are the nearest neighbors and regions of the genome that are different for a 0×WGD cell (middle) and a 1×WGD cell (right). **B.** Empirical distribution of NND for all cells, and beta distribution fit (red). **C.** NND (y-axis) by ploidy (x-axis) for cells from patient 081. Color indicates #WGD and divergent status. **D.** Copy-number profiles for example 0×WGD (top), 1×WGD (middle) and divergent (bottom) cells from patient 081. **E.** Arm nullisomy rates (counts per cell) for divergent and non-divergent cells in rare and prevalent WGD patients. Shown is the distribution of mean rates per population in each patient. **F.** MEDICC2 phylogeny (left) total copy number (center) and inferred cell specific copy number changes (right) for patient 110. **G.** Rates of chromosome, arm, and segment losses and gains (counts per cell) normalized for increased or decreased opportunity for an event based on genomic content in each cell’s ancestor. MWU significance is annotated as ‘ns’: 5.0×10^-2^ < *p* <= 1.0, ‘*’: 1.0×10^-2^ < *p* <= 5.0×10^-2^, ‘**’: 1.0×10^-3^ < *p* <= 1.0×10^-2^, ‘***’: 1.0×10^-4^ < *p* <= 1.0×10^-3^, ‘****’: *p* <= 1.0×10^-4^. **H** Number of focal high level amplifications per patient detected in the PCAWG ovarian cohort, split by WGD vs non-WGD. **I.** Number of post-WGD chromosome and arm gains and losses (x-axis) compared to the mutation time in C>T CpG counts (y-axis) measured since the WGD event.

**Extended Data Figure 5.**
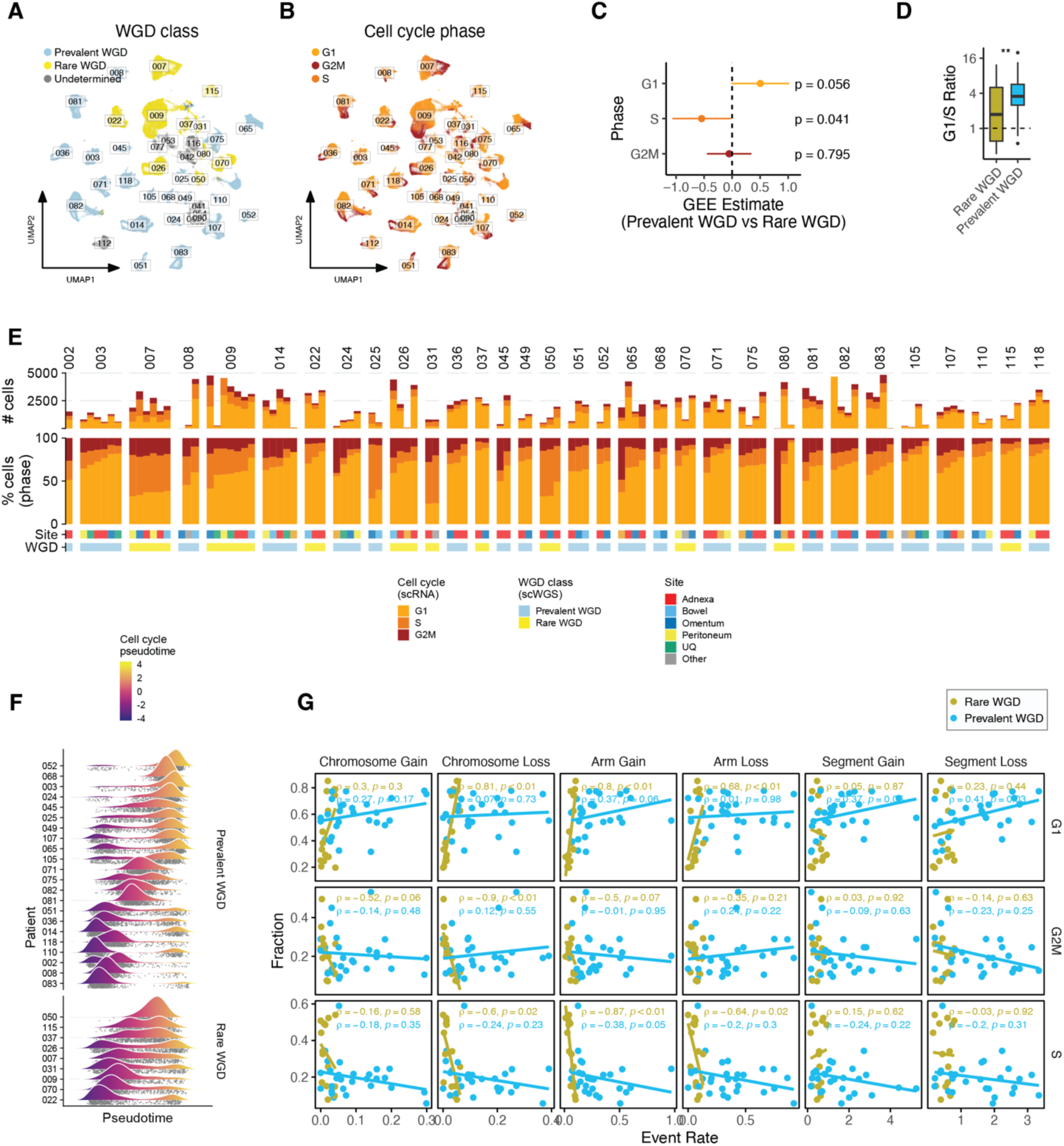
Cell cycle progression in the context of whole genome doubling. **A.** UMAP of cancer cells colored by Rare vs Prevalent WGD patient labels. **B.** UMAP of cancer cells colored by inferred cell cycle state. **C.** Coefficients (x-axis) of a Generalized Estimation Equation (GEE) fit to the difference in cancer cell cycle fractions between Rare and Prevalent WGD samples, corrected for patient effects. Significance of WGD effect on cell cycle fractions are shown at right. **D.** Distribution of G1/S cancer cell cycle ratios for Rare and Prevalent WGD samples. **E.** Absolute and relative compositions of cell cycle fractions in CD45^-^ sorted samples based on scRNA-seq. Samples are separated by patient and ordered by proportion of S-phase cells out of all cancer cells. **F.** Distribution of cell cycle pseudotime estimates over all cells for each patient, separated into Prevalent WGD (top) and Rare WGD (bottom). **G.** Correlation between the fraction of cancer cells in G1, S and G2M phase (y-axis) and rates (counts per cell) of chromosome, arm, and segment losses and gains (x-axes).

**Extended Data Figure 6.**
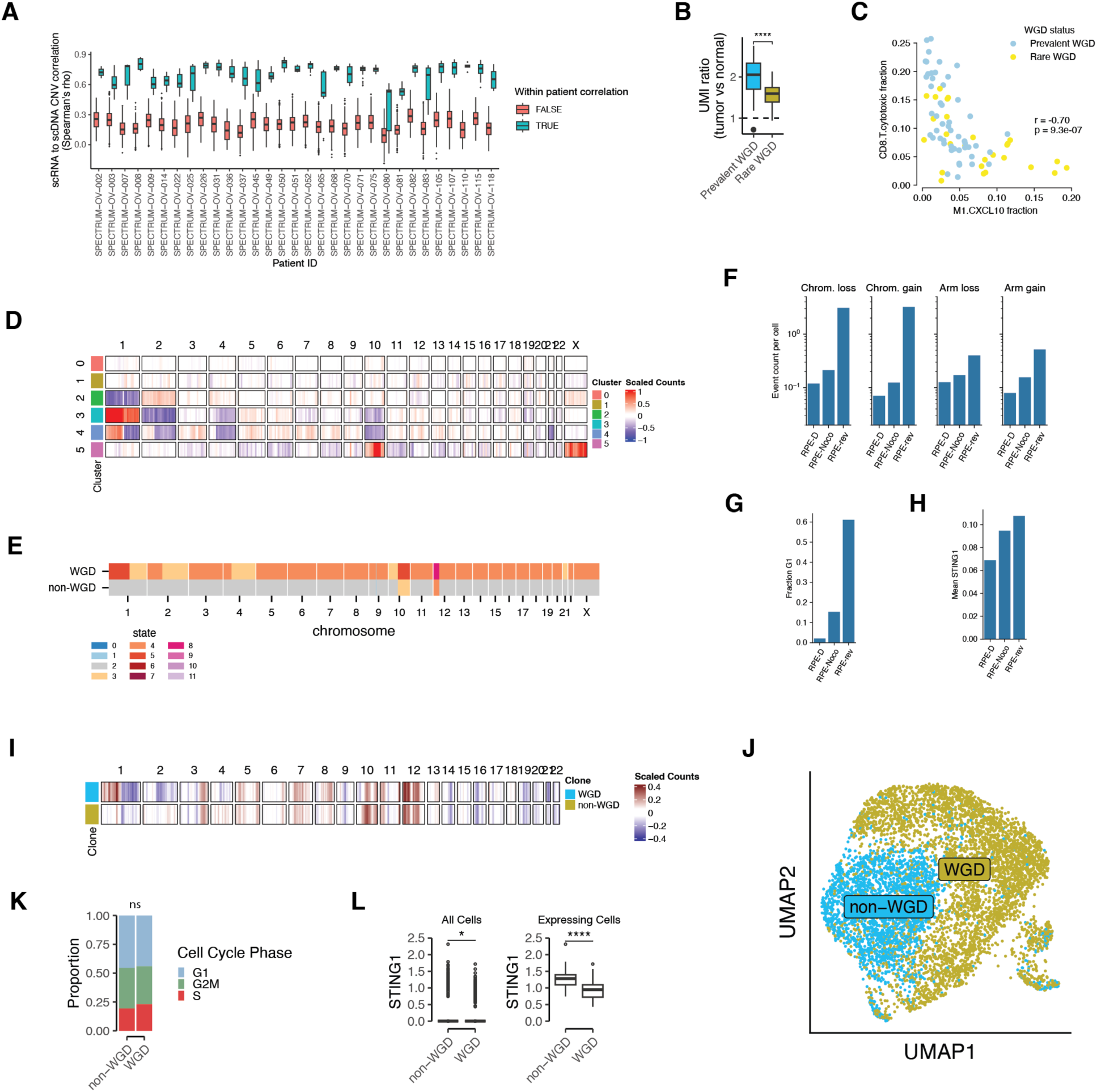
Tumor cell phenotypes and microenvironment remodeling in the context of whole genome doubling. **A.** Correlation between DLP+ and scRNA based copy number. Data points for box plots are scRNA inferCNV copy number clusters. The y-axis shows correlation between each DLP+ sbmclone cluster and each scRNA copy number cluster from the same patient (blue). As a comparator we show the same correlation computed with each DLP+ SBMClone cluster from any other patient (red). **B.** Ratio of cancer cell UMI counts to fibroblast and endothelial cell UMI counts, averaged within each patient. Patients are grouped by Rare vs Prevalent WGD. **C.** Cytotoxic CD8^+^ T cells (y-axis) and *CXCL10*^+^*CD274*^+^ Macrophages (x-axis) as fractions of CD45^+^ cells across CD45^+^ samples. Points are colored by the WGD class of the patient from which the sample originated. **D.** Copy number inferred from scATAC for RPE1 cells across treatment conditions. **E.** Clone copy number inferred from DLP for RPE1 cells across treatment conditions. Two clones were identified: one WGD and one non-WGD. **F.** Chromosome and arm loss and gain events per cell for non-WGD RPE1 cells treated with DMSO control (RPE-D), nocodazole (RPE-noco) and reversine (RPE-rev). **G.** Fraction of non-WGD RPE1 cells within G1 phase (y-axis) for each treatment condition: DMSO control (RPE-D), nocodazole (RPE-noco) and reversine (RPE-rev). **H.** Average STING1 expression (y-axis) for non-WGD RPE1 cells by treatment condition (x-axis): DMSO control (RPE-D), nocodazole (RPE-noco) and reversine (RPE-rev). **I.** WGD and non-WGD copy number clones inferred from scRNA-seq of sample RPE-WGD. **J.** Expression UMAP from scRNA-seq of sample RPE-WGD with cells colored by assignment to the WGD and non-WGD clones. **K.** Cell cycle fractions for WGD and non-WGD clones in the RPE-WGD sample. **L.** Expression of STING1 across all cells (left) and in cells with positive expression (right) in the RPE-WGD sample.

## TABLES

**Supplementary Table 1**

Clinical overview of the MSK SPECTRUM patient cohort. Data include patient age at diagnosis, staging following FIGO Ovarian Cancer Staging guidelines, type of surgical procedure, WGD class, and mutational signature subtype.

**Supplementary Table 2**

Sample inventory. Metadata associated with scWGS, scRNA-seq, H&E, IF, bulk tumor and normal WGS, and tumor and normal MSK-IMPACT datasets.

## Notes

### Competing Interest Statement

B.W. reports grant funding by Repare Therapeutics paid to the instiution, outside the submitted work, and employment of a direct family member at AstraZeneca. C.F. reports research funding to the institution from Merck, Astra Zeneca, Genentech/Roche, Bristol Myer Squibb, and Daiichi; uncompensated membership of a scientific advisory board for Merck and Genentech; and is a consultant for OncLive, Aptitude Health, Bristol Myers Squibb and Seagen, all outside the scope of this manuscript.
D.C. Medical Advisory Board: Verthermia Acquio Inc, Biom'up; Paid Speaker: Astra Zeneca 2021-present; Stock Holder: Doximity 2021-present; Moderna 2021; BioNTech 2021.
D.Z. reports institutional grants from Merck, Genentech, AstraZeneca, Plexxikon, and Synthekine, and personal fees from AstraZeneca, Xencor, Memgen, Takeda, Astellas, Immunos, Tessa Therapeutics, Miltenyi, and Calidi Biotherapeutics. D.Z. own a patent on use of oncolytic Newcastle Disease Virus for cancer therapy.
N.A. Grants to institution from Stryker/Novadaq and GRAIL, outside the submitted work.
R.N.G. reports funding from GSK, Novartis, Mateon Therapeutics, Corcept, Regeneron, Clovis, Context Therapeutics, EMD Serono, MCM Education, OncLive, Aptitude Health and Prime Oncology, outside this work.
S.F.B. owns equity in, receives compensation from, and serves as a consultant and the Scientific Advisory Board and Board of Directors of Volastra Therapeutics Inc. He also serves on the SAB of Meliora Therapeutics Inc.
S.P.S. reports research funding from AstraZeneca and Bristol Myers Squibb, outside the scope of this work; SPS is a consultant and shareholder of Canexia Health Inc.
Y.L.L. reports research funding from AstraZeneca, GSK/Tesaro, Artios Pharma, and Tesaro Therapeutics outside this work.
Y.L. reports serving as a consultant for Calyx Clinical Trial Solutions outside this work.

### Summary of Updates

Authors have been re-ordered to be consistent with the intended author order as shown in the document. Corrected figure 1D and 1E that were erroneously swapped.

