## Supplementary Note for "Ongoing genome doubling promotes evolvability and immune dysregulation in ovarian cancer"

### Supplementary Information for “Ongoing genome doubling promotes evolvability and immune dysregulation in ovarian cancer”

#### Table of contents

1. scWGS cell filtering evidence and examples 3
 1.1. Supporting evidence for ploidy calls from odd copy-number states 3
 1.2. Supporting evidence for S-phase filtering 3
 1.3. Example cells by filtering classification 4
  1.3.1. Example normal cells 6
  1.3.2. Example aberrant normal cells 8
  1.3.3. Example S-phase cells 10
  1.3.4. Example doublets (identified from manual review of spotter images) 12
  1.3.5. Example multipolar cells (model-based classification) 14
  1.3.6. Example multiplets (model-based classification) 16
  1.3.7. Example filter-pass cells 18
2. Discerning independent WGD from shared WGD using SNVs 20
3. Benchmarking estimation of chromosomal instability using simulations 23
 3.1. Generation of simulated data 23
 3.1.2. Results on simulated data 25
4. Extended SBMClone results and validation 26
 4.1. SBMClone block density matrices 26
 4.2. Confidence in SBMClone clone assignments 28
 4.3. Consistency between SBMClone clusters and copy-number states 30
5. Assignment of SNVs to clone tree branches 32
References 35

#### 1. scWGS cell filtering evidence and examples

##### 1.1. Supporting evidence for ploidy calls from odd copy-number states

For each scWGS cell in the cohort, we computed the proportion of the genome assigned to copy numbers 1, 3, or 5 as well as the longest contiguous segment assigned to copy number 1, 3, or 5. Among cells with haplotype-specific copy number profiles that suggest 1 or 2 WGDs, those with only a small proportion of the genome and without any long segments in these odd states are likely to be spurious WGD calls. As such, we remove any cells with a longest-1/3/5-segment shorter than 10 Mb from analysis.

##### 1.2. Supporting evidence for S-phase filtering

Here we show an example of the 3 statistics used to identify S-phase cells for patient 002 (**Supplementary Figure 1-2**). The thresholds for each statistic were tuned manually for each patient to account for the varying aneuploidy in typical tumor cells. A cell was called S-phase if it was outside the thresholds for at least 2 out of the 3 statistics.

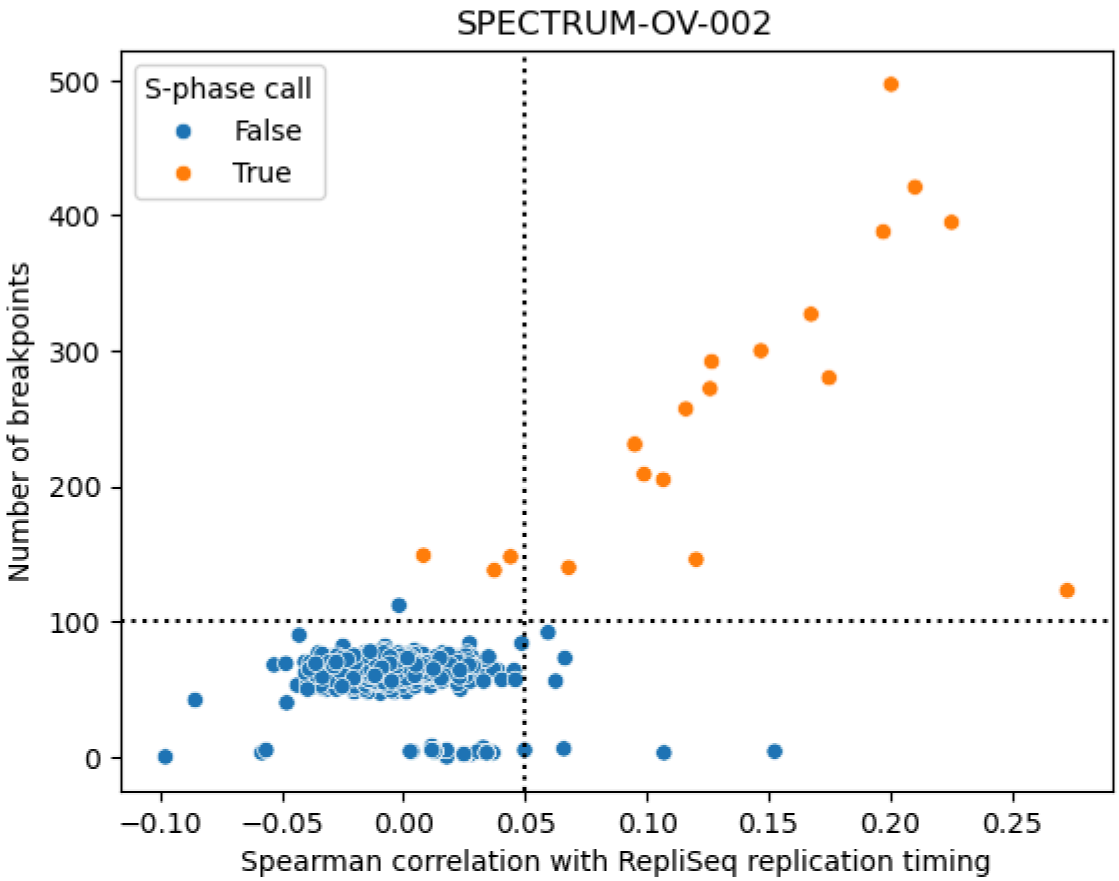


**Supplementary Figure 1. Replication timing and total breakpoints for cells from patient 002.** This figure shows two S-phase statistics for cells from patient 002: Spearman correlation with replication timing (x-axis) and number of breakpoints (y-axis). Cells are colored by the ultimate S-phase call, which requires that 2 of the 3 S-phase statistics exceed the manually determined patient-specific thresholds.

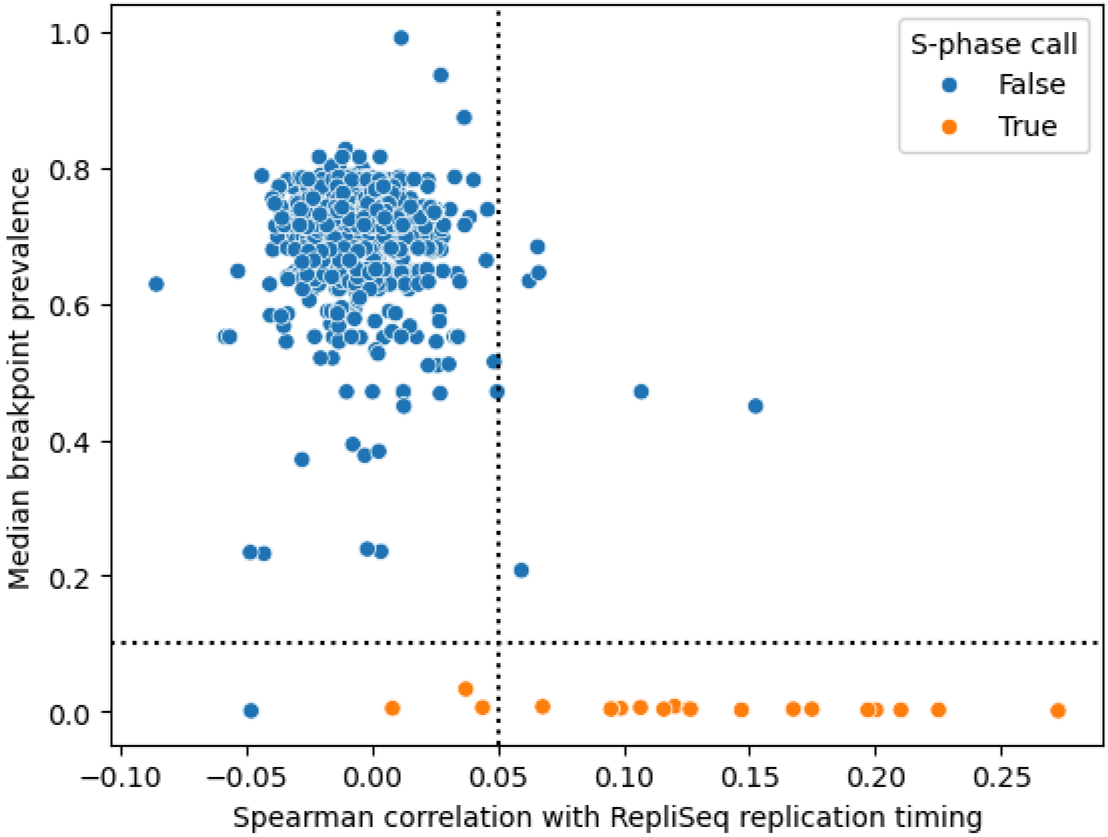


**Supplementary Figure 2. Replication timing and median breakpoint prevalence for cells from patient 002.** This figure shows Spearman correlation with replication timing (x-axis) and the median prevalence of copy-number breakpoints present in the cell (y-axis). Cells are colored by the ultimate S-phase call, which requires that 2 of the 3 S-phase statistics exceed the manually determined patient-specific thresholds.

##### 1.3. Example cells by filtering classification

For each filtering step, we show 4 example cells (**Supplementary Figures 3-14**) that were identified as violating the filter (or passing all filters, for the "filter-pass" section: **Supplementary Figures 15-16**). For a description of the filtering steps used to identify these cells for removal, see Methods.

For each cell, we show 3 views:

1. GC-corrected and ploidy-scaled read counts across the genome (left, "copy") colored by the total copy-number state assigned by HMMCopy.
2. Phased BAF across the genome (right, "BAF") inferred by signals colored by the minor copy number inferred by signals.

3. Image from the DLP+ cell spotter. The ejection nozzle is to the left of the region shown in this image. The left vertical line indicates the rightmost edge of the ejection zone (fluid that is guaranteed to be ejected), and the right vertical line indicates the rightmost edge of the sedimentary zone (fluid that may or may not be ejected). Ejection into a well occurs only if exactly 1 cell is identified in either of these regions. The green cross marks the object that was identified as a cell by the automated imaging pipeline.

*1.3.1. Example normal cells*

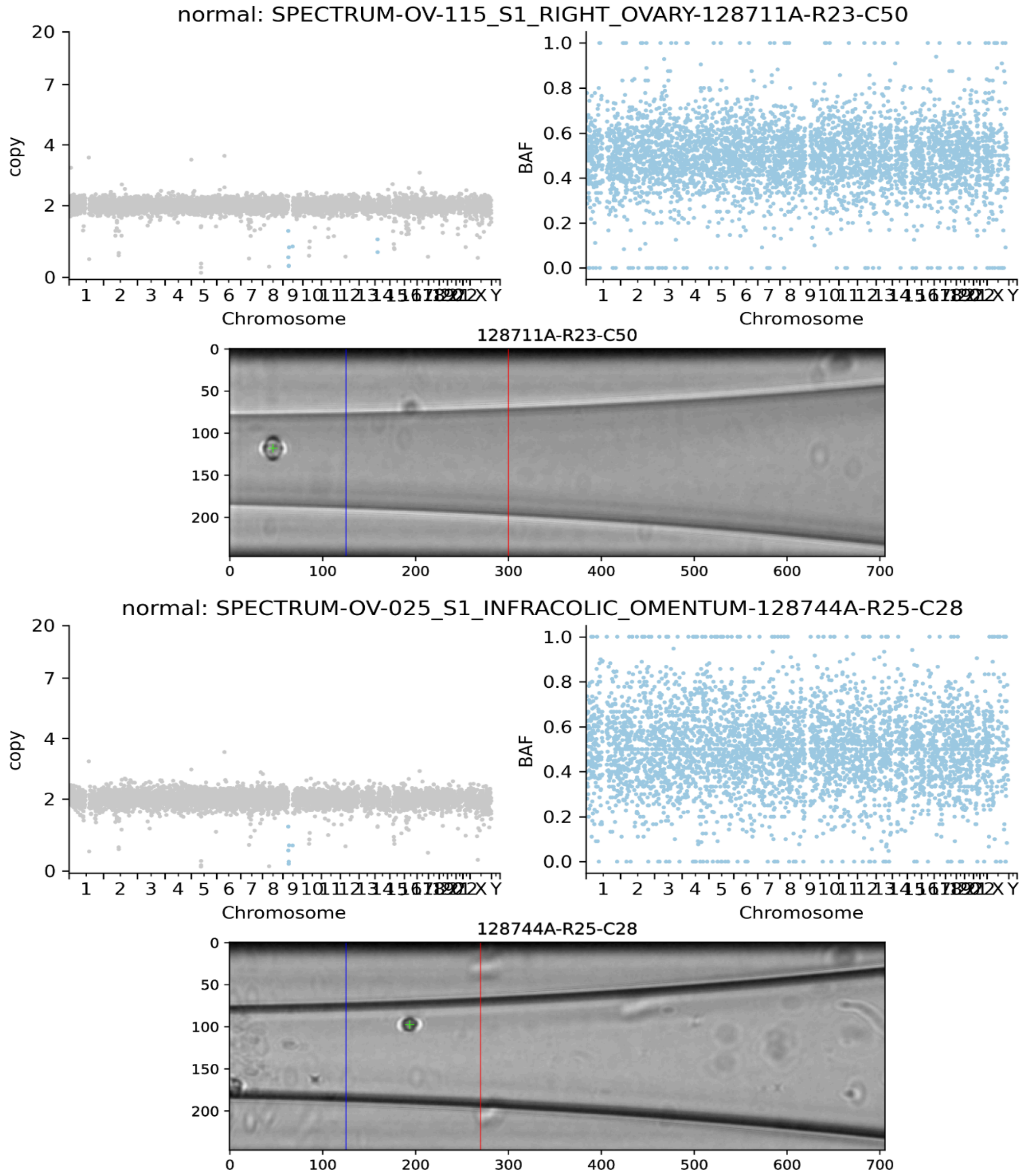


**Supplementary Figure 3. Two example cells classified as normal by the QC pipeline.** Scatterplots show the normalized and ploidy-scaled read-depth of each bin colored by HMMCopy-inferred copy-number state (left) and B-allele frequency of each bin colored by signals-inferred minor copy-number state (right). Images for each cell (center) show the image taken through the DLP+ nozzle where a green cross indicates the detected cell object, the left vertical line indicates the rightmost edge of the ejection zone, and the right vertical line indicates the rightmost edge of the sedimentary zone.

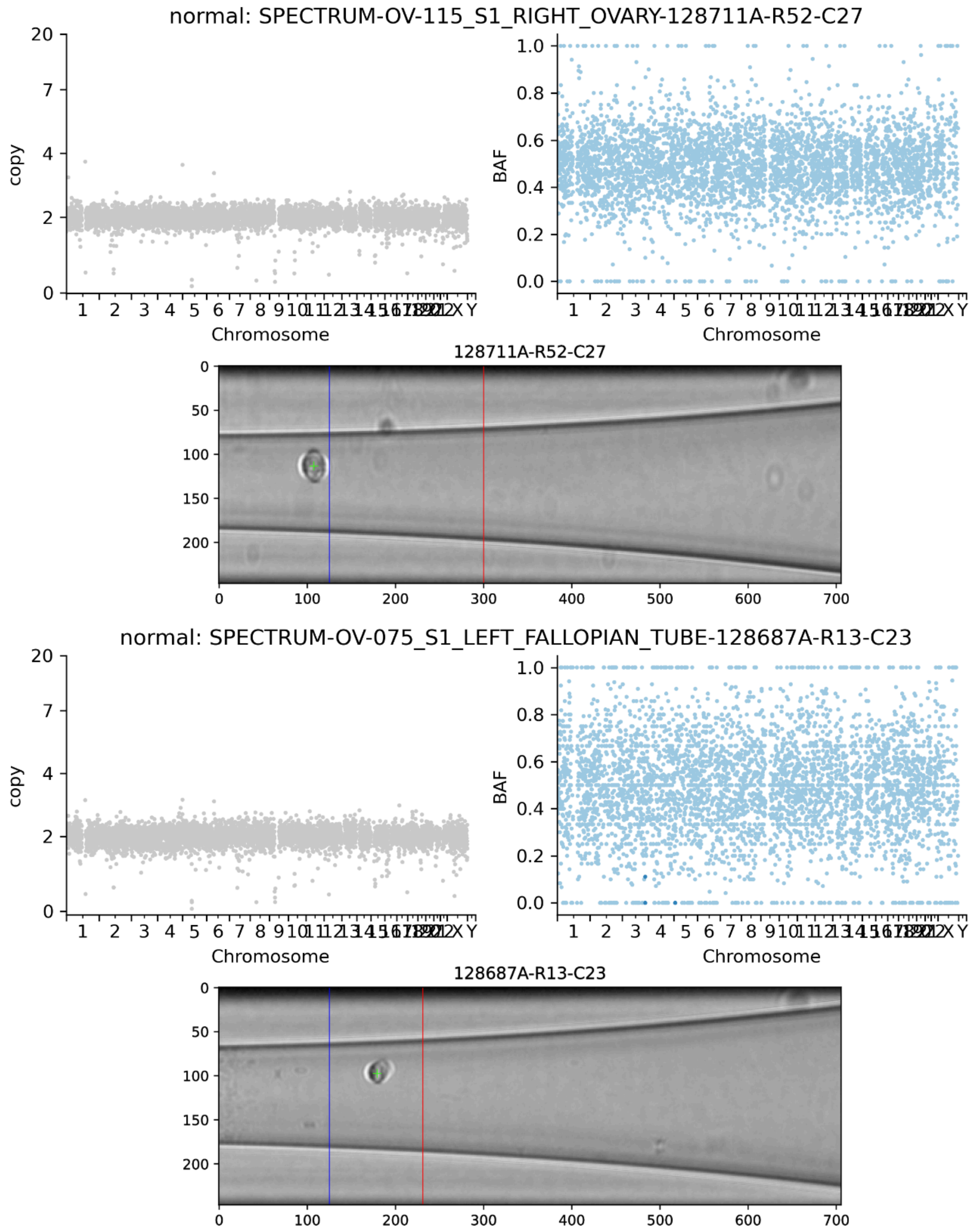


**Supplementary Figure 4. Two example cells classified as normal by the QC pipeline.** Scatterplots show the normalized and ploidy-scaled read-depth of each bin colored by HMMCopy-inferred copy-number state (left) and B-allele frequency of each bin colored by signals-inferred minor copy-number state (right). Images for each cell (center) show the image taken through the DLP+ nozzle where a green cross indicates the detected cell object, the left vertical line indicates the rightmost edge of the ejection zone, and the right vertical line indicates the rightmost edge of the sedimentary zone.

*1.3.2. Example aberrant normal cells*

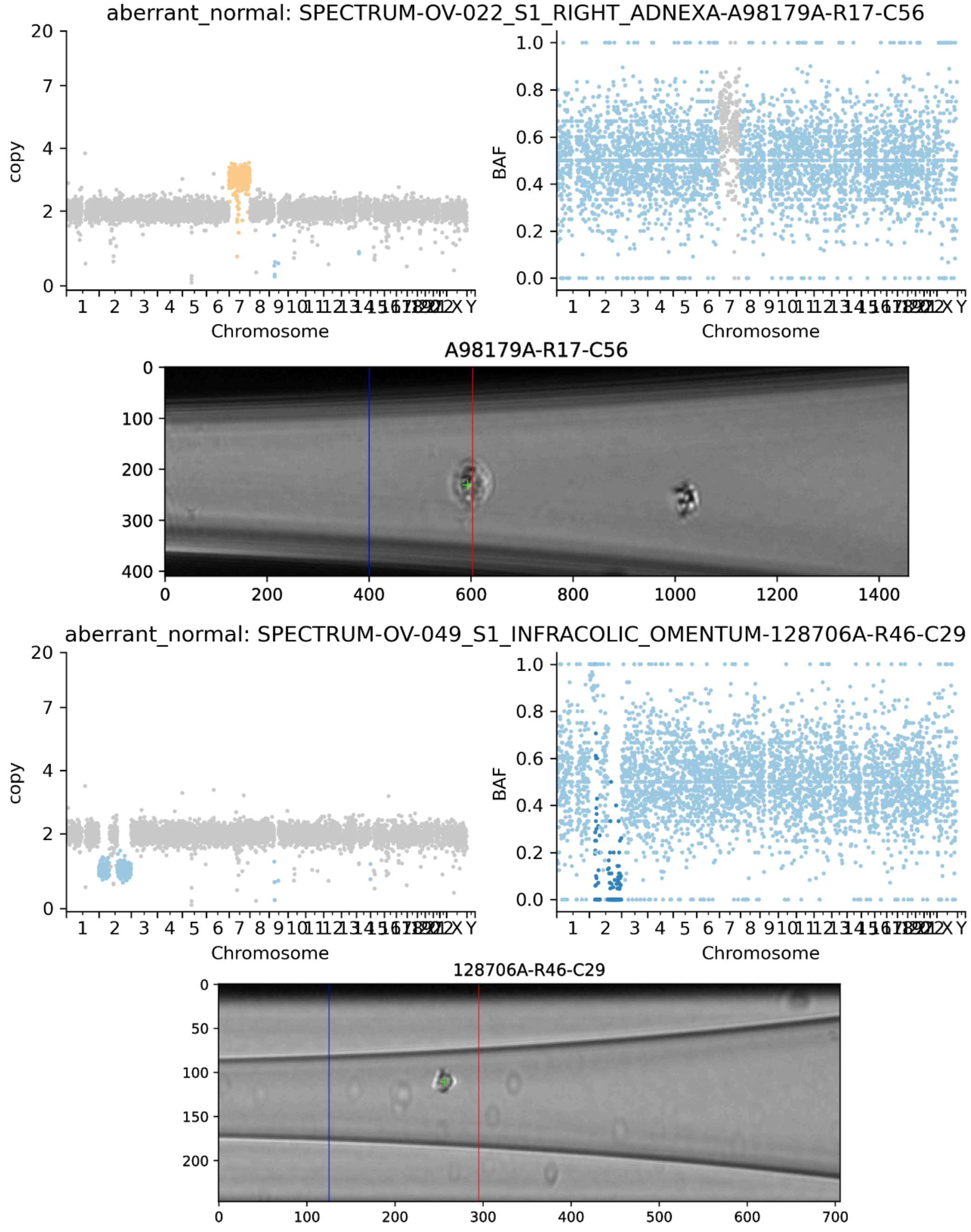


**Supplementary Figure 5. Two example cells classified as aberrant normal by the QC pipeline.** Scatterplots show the normalized and ploidy-scaled read-depth of each bin colored by HMMCopy-inferred copy-number state (left) and B-allele frequency of each bin colored by signals-inferred minor copy-number state (right). Images for each cell (center) show the image taken through the DLP+ nozzle where a green cross indicates the detected cell object, the left vertical line indicates the rightmost edge of the ejection zone, and the right vertical line indicates the rightmost edge of the sedimentary zone.

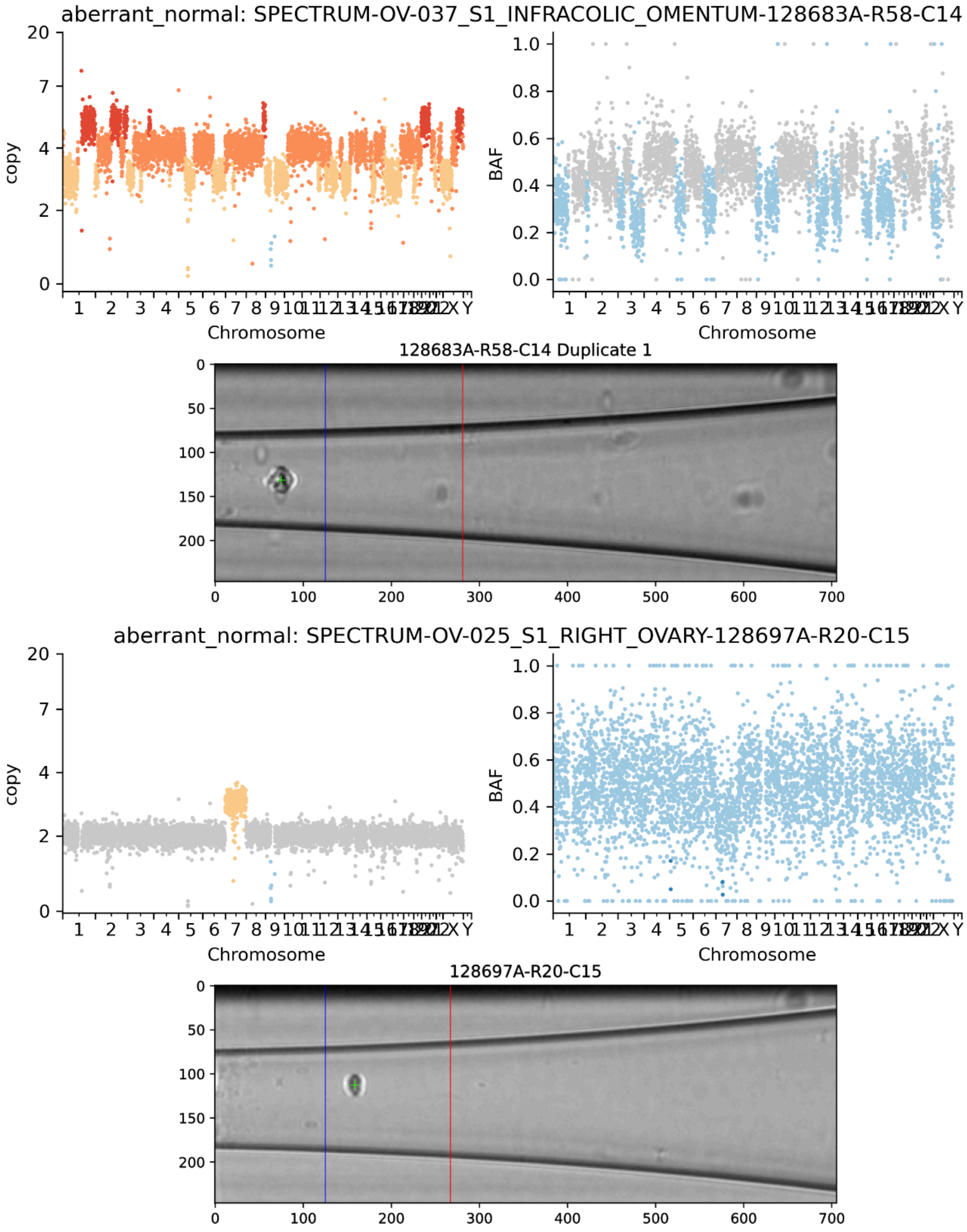


**Supplementary Figure 6. Two example cells classified as aberrant normal by the QC pipeline.** Scatterplots show the normalized and ploidy-scaled read-depth of each bin colored by HMMCopy-inferred copy-number state (left) and B-allele frequency of each bin colored by signals-inferred minor copy-number state (right). Images for each cell (center) show the image taken through the DLP+ nozzle where a green cross indicates the detected cell object, the left vertical line indicates the rightmost edge of the ejection zone, and the right vertical line indicates the rightmost edge of the sedimentary zone.

*1.3.3. Example S-phase cells*

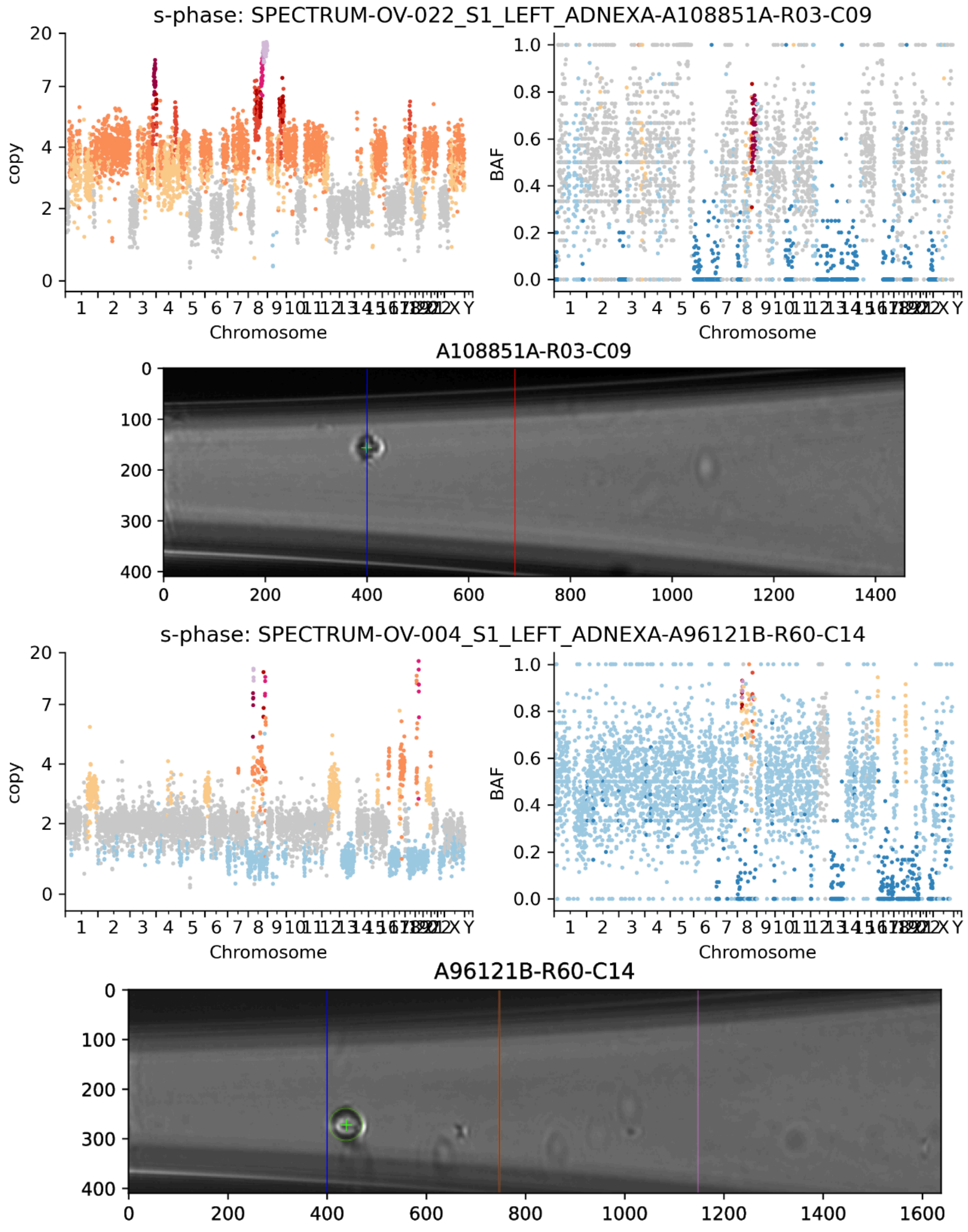


**Supplementary Figure 7. Two example cells classified as S-phase by the QC pipeline.** Scatterplots show the normalized and ploidy-scaled read-depth of each bin colored by HMMCopy-inferred copy-number state (left) and B-allele frequency of each bin colored by signals-inferred minor copy-number state (right). Images for each cell (center) show the image taken through the DLP+ nozzle where a green cross indicates the detected cell object, the left vertical line indicates the rightmost edge of the ejection zone, and the right vertical line indicates the rightmost edge of the sedimentary zone.

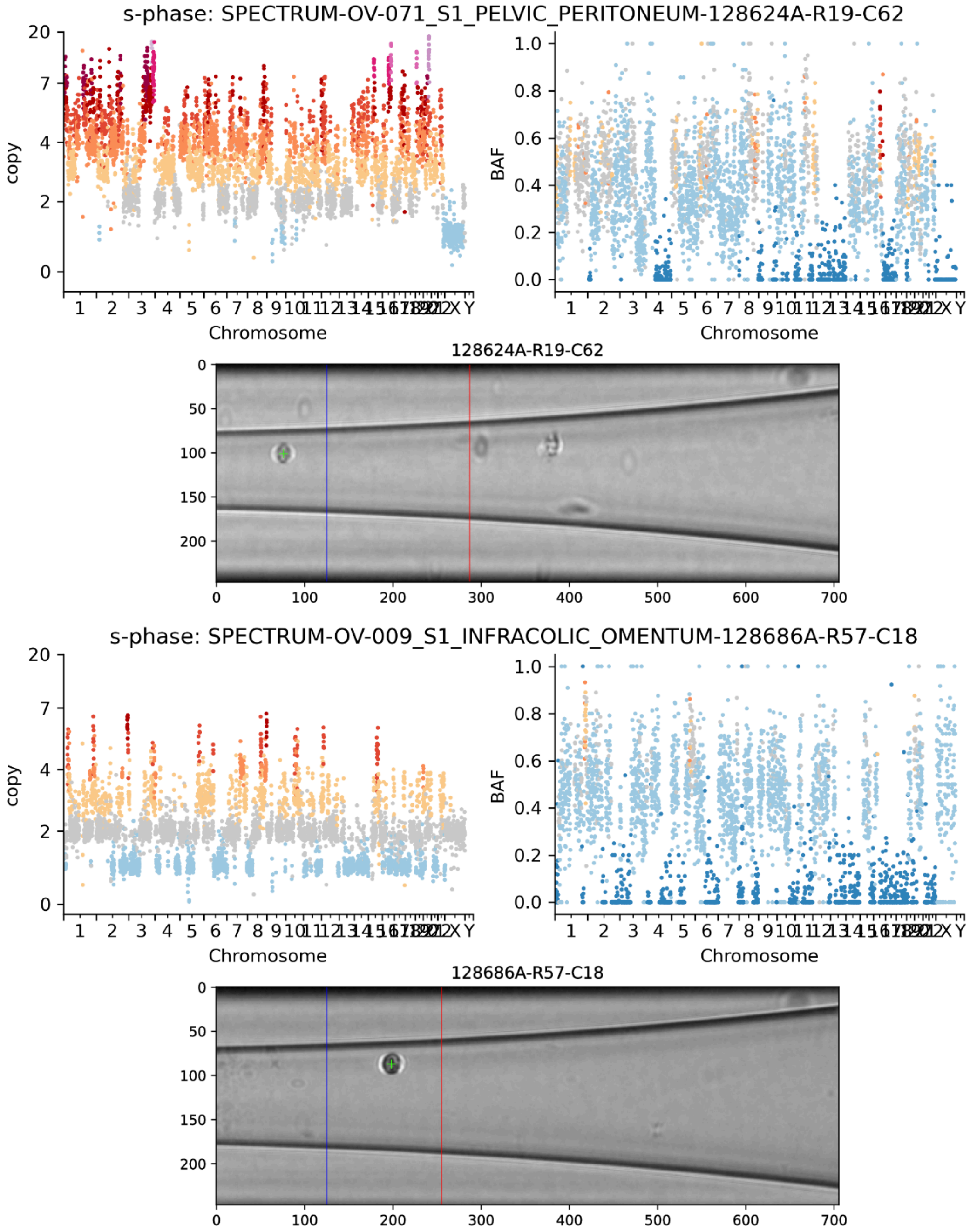


**Supplementary Figure 8. Two example cells classified as S-phase by the QC pipeline.** Scatterplots show the normalized and ploidy-scaled read-depth of each bin colored by HMMCopy-inferred copy-number state (left) and B-allele frequency of each bin colored by signals-inferred minor copy-number state (right). Images for each cell (center) show the image taken through the DLP+ nozzle where a green cross indicates the detected cell object, the left vertical line indicates the rightmost edge of the ejection zone, and the right vertical line indicates the rightmost edge of the sedimentary zone.

*1.3.4. Example doublets (identified from manual review of spotter images)*

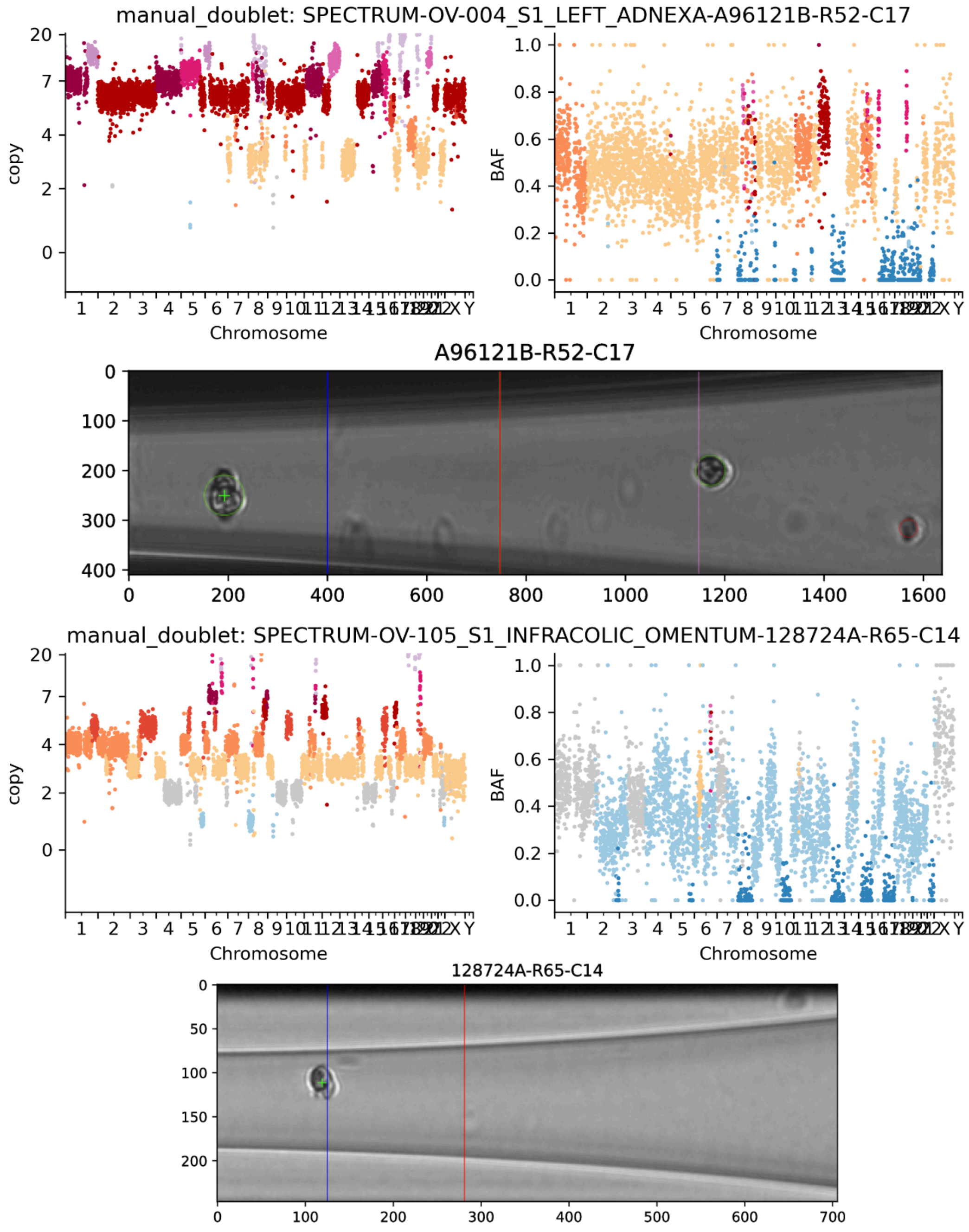


**Supplementary Figure 9. Two example cells classified as doublets based on manual review of the spotter images.** Scatterplots show the normalized and ploidy-scaled read-depth of each bin colored by HMMCopy-inferred copy-number state (left) and B-allele frequency of each bin colored by signals-inferred minor copy-number state (right). Images for each cell (center) show the image taken through the DLP+ nozzle where a green cross indicates the detected cell object, the left vertical line indicates the rightmost edge of the ejection zone, and the right vertical line indicates the rightmost edge of the sedimentary zone.

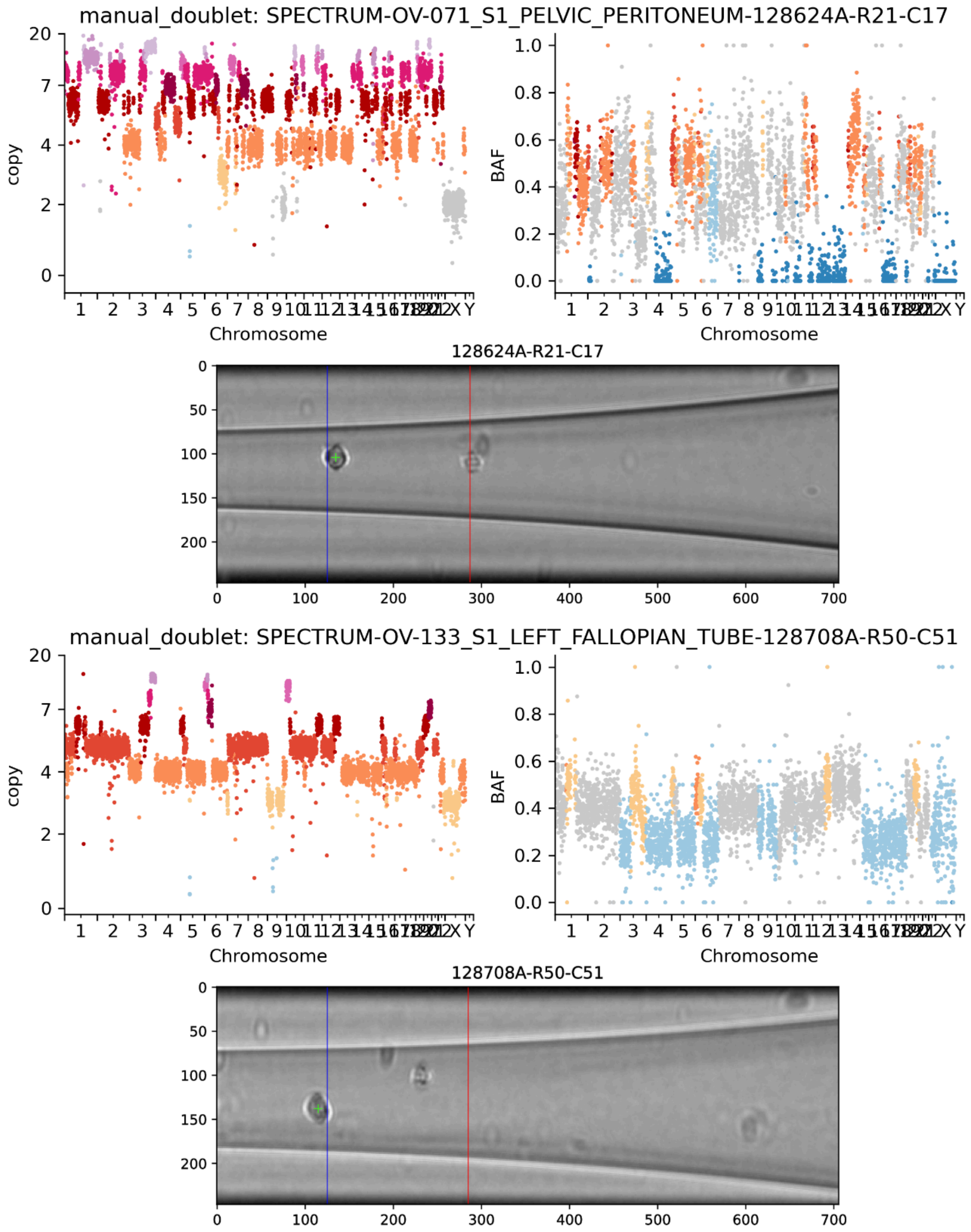


**Supplementary Figure 10. Two example cells classified as doublets based on manual review of the spotter images.** Scatterplots show the normalized and ploidy-scaled read-depth of each bin colored by HMMCopy-inferred copy-number state (left) and B-allele frequency of each bin colored by signals-inferred minor copy-number state (right). Images for each cell (center) show the image taken through the DLP+ nozzle where a green cross indicates the detected cell object, the left vertical line indicates the rightmost edge of the ejection zone, and the right vertical line indicates the rightmost edge of the sedimentary zone.

*1.3.5. Example multipolar cells (model-based classification)*

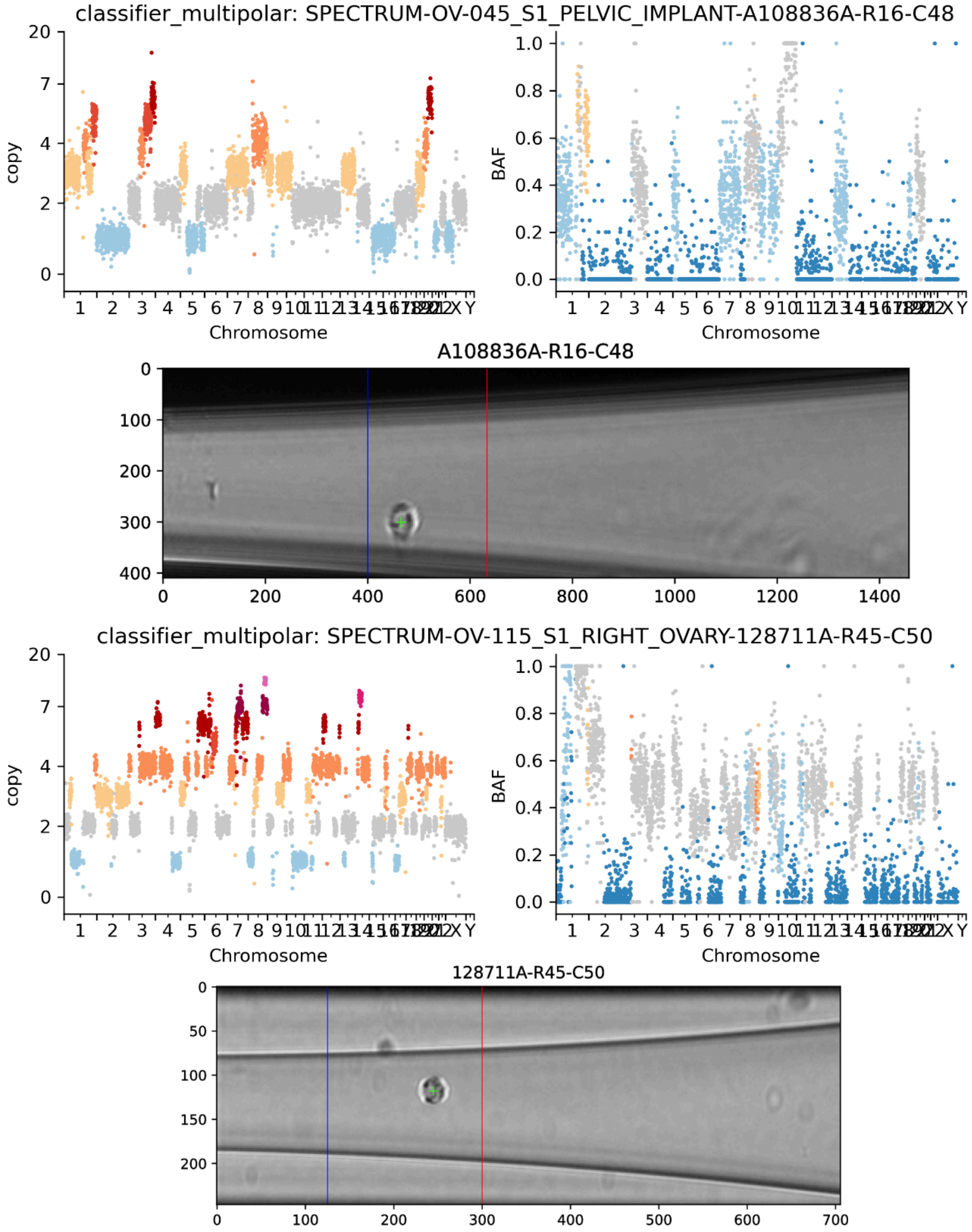


**Supplementary Figure 11. Two example cells classified as multipolar by the model of copy-number event rates.** Scatterplots show the normalized and ploidy-scaled read-depth of each bin colored by HMMCopy-inferred copy-number state (left) and B-allele frequency of each bin colored by signals-inferred minor copy-number state (right). Images for each cell (center) show the image taken through the DLP+ nozzle where a green cross indicates the detected cell object, the left vertical line indicates the rightmost edge of the ejection zone, and the right vertical line indicates the rightmost edge of the sedimentary zone.

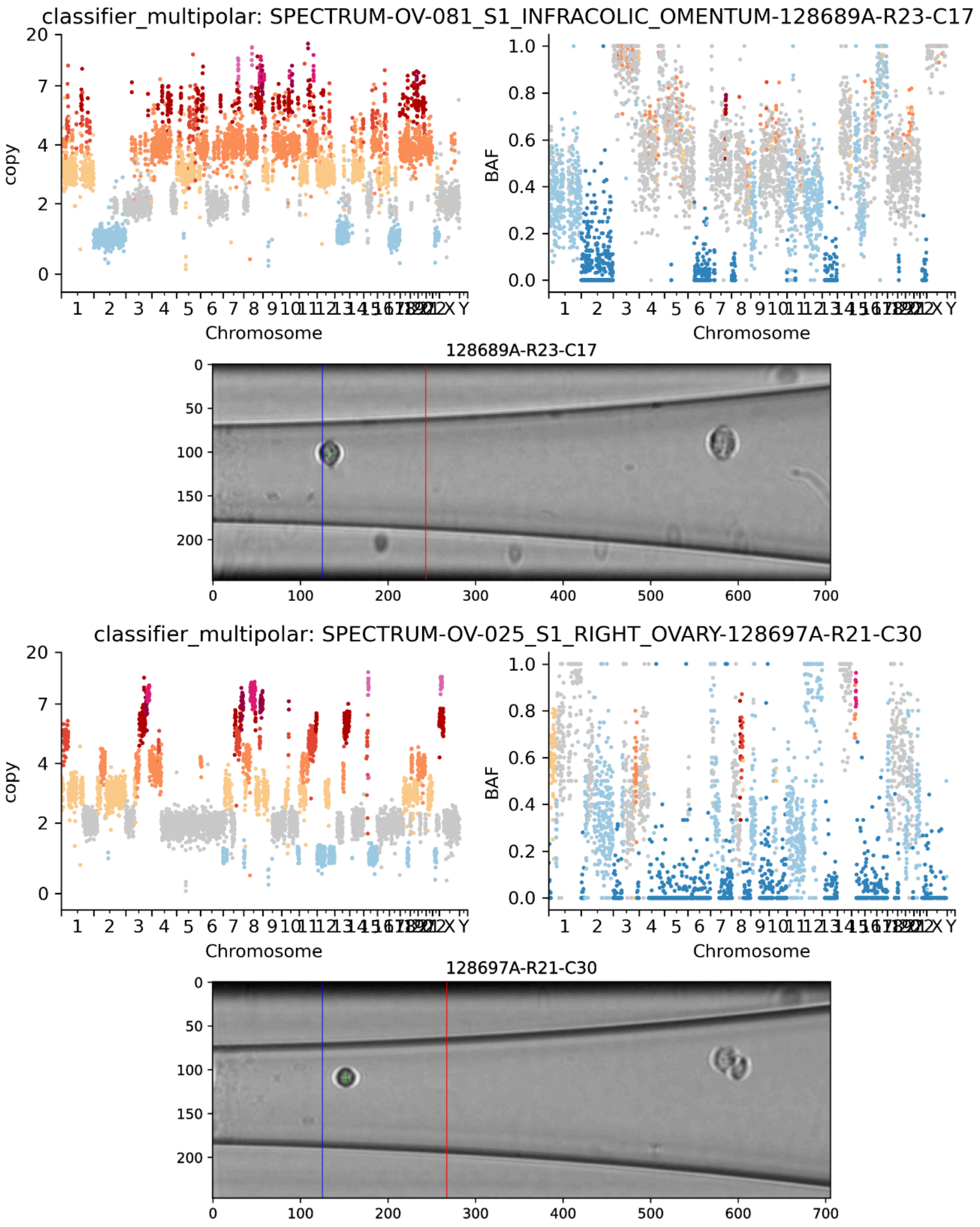


**Supplementary Figure 12. Two example cells classified as multipolar by the model of copy-number event rates.** Scatterplots show the normalized and ploidy-scaled read-depth of each bin colored by HMMCopy-inferred copy-number state (left) and B-allele frequency of each bin colored by signals-inferred minor copy-number state (right). Images for each cell (center) show the image taken through the DLP+ nozzle where a green cross indicates the detected cell object, the left vertical line indicates the rightmost edge of the ejection zone, and the right vertical line indicates the rightmost edge of the sedimentary zone.

*1.3.6. Example multiplets (model-based classification)*

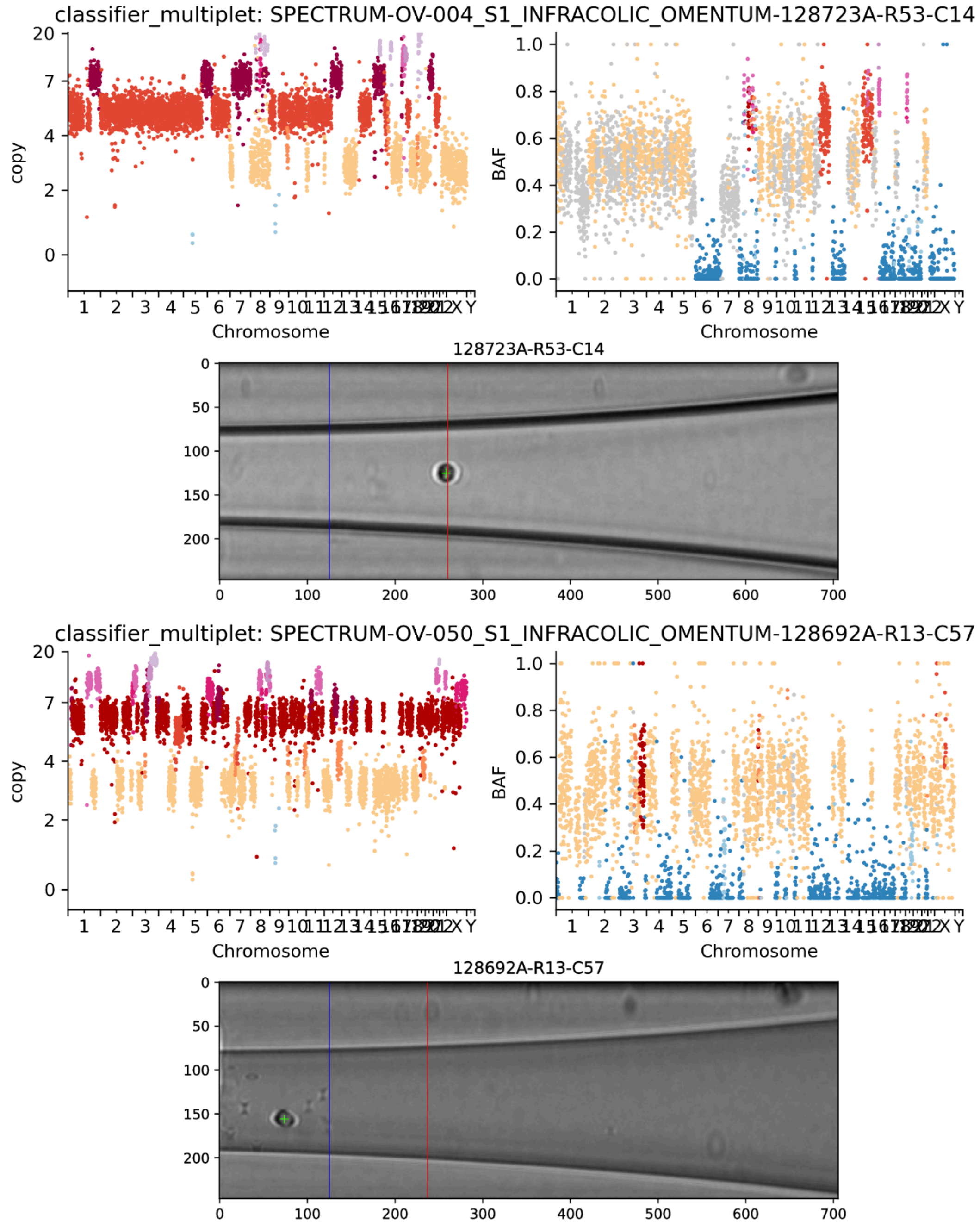


**Supplementary Figure 13. Two example cells classified as multiplets by the model of copy-number event rates.** Scatterplots show the normalized and ploidy-scaled read-depth of each bin colored by HMMCopy-inferred copy-number state (left) and B-allele frequency of each bin colored by signals-inferred minor copy-number state (right). Images for each cell (center) show the image taken through the DLP+ nozzle where a green cross indicates the detected cell object, the left vertical line indicates the rightmost edge of the ejection zone, and the right vertical line indicates the rightmost edge of the sedimentary zone.

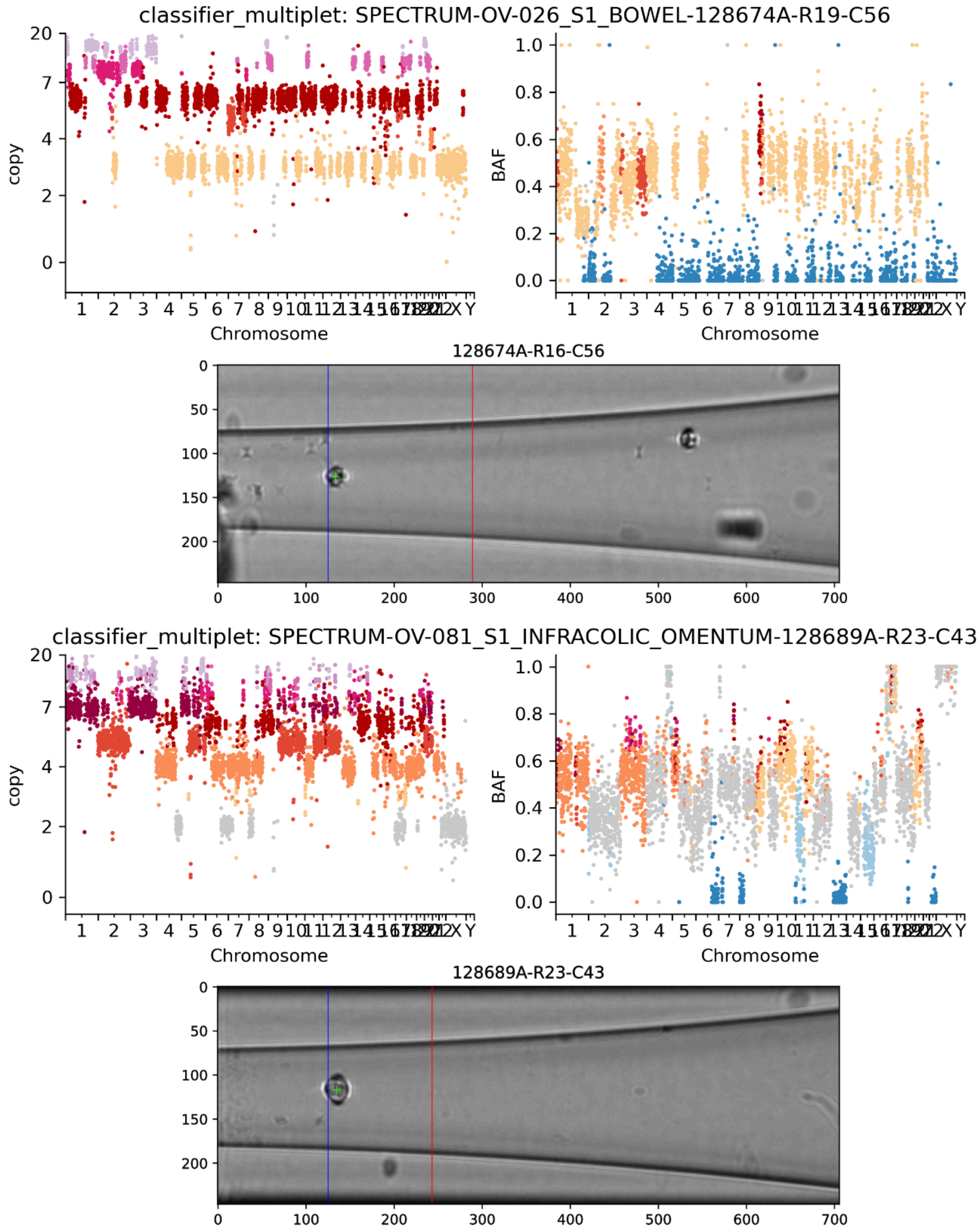


**Supplementary Figure 14. Two example cells classified as multiplets by the model of copy-number event rates.** Scatterplots show the normalized and ploidy-scaled read-depth of each bin colored by HMMCopy-inferred copy-number state (left) and B-allele frequency of each bin colored by signals-inferred minor copy-number state (right). Images for each cell (center) show the image taken through the DLP+ nozzle where a green cross indicates the detected cell object, the left vertical line indicates the rightmost edge of the ejection zone, and the right vertical line indicates the rightmost edge of the sedimentary zone.

*1.3.7. Example filter-pass cells*

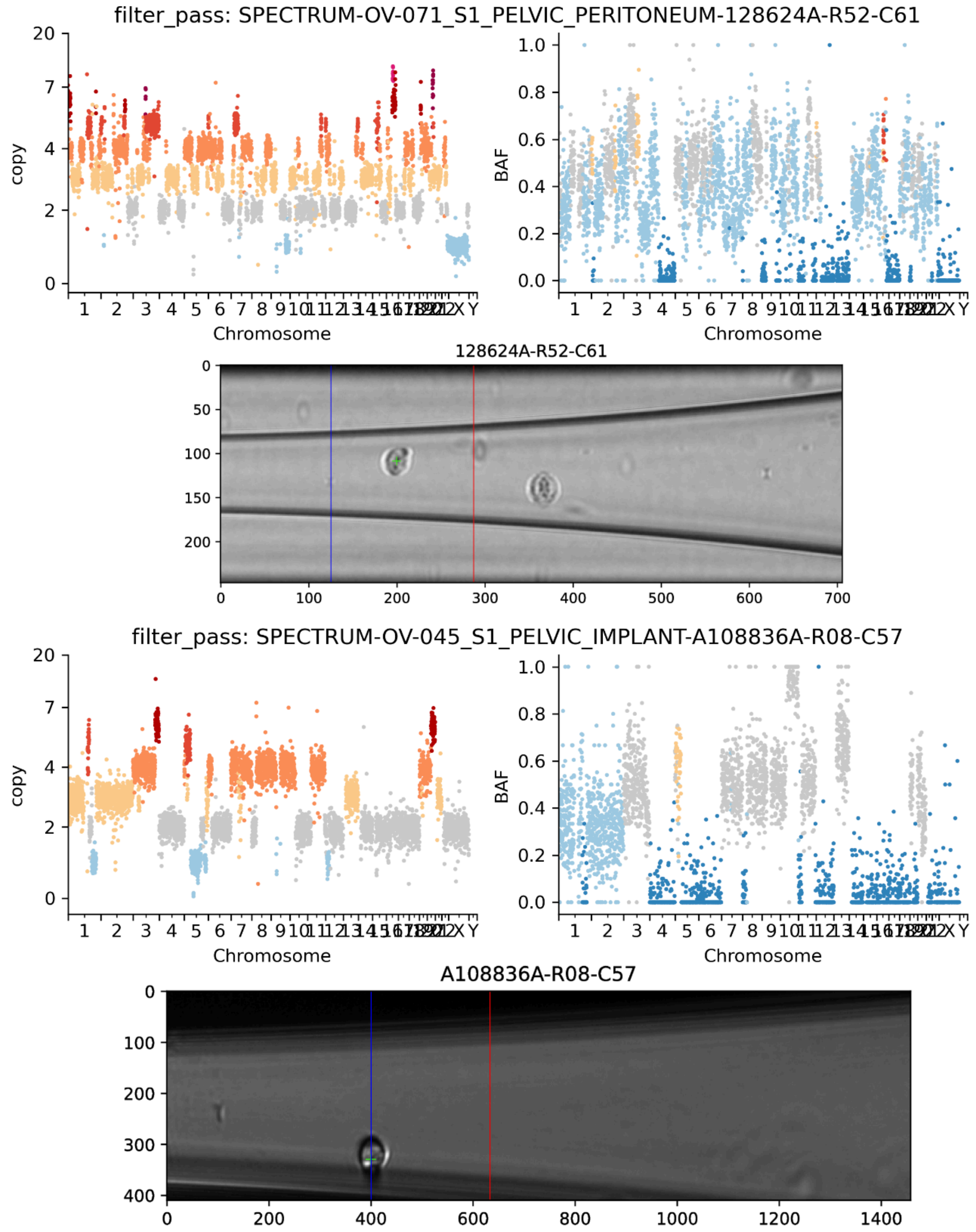


**Supplementary Figure 15. Two example cells that passed all QC filters.** Scatterplots show the normalized and ploidy-scaled read-depth of each bin colored by HMMCopy-inferred copy-number state (left) and B-allele frequency of each bin colored by signals-inferred minor copy-number state (right). Images for each cell (center) show the image taken through the DLP+ nozzle where a green cross indicates the detected cell object, the left vertical line indicates the rightmost edge of the ejection zone, and the right vertical line indicates the rightmost edge of the sedimentary zone.

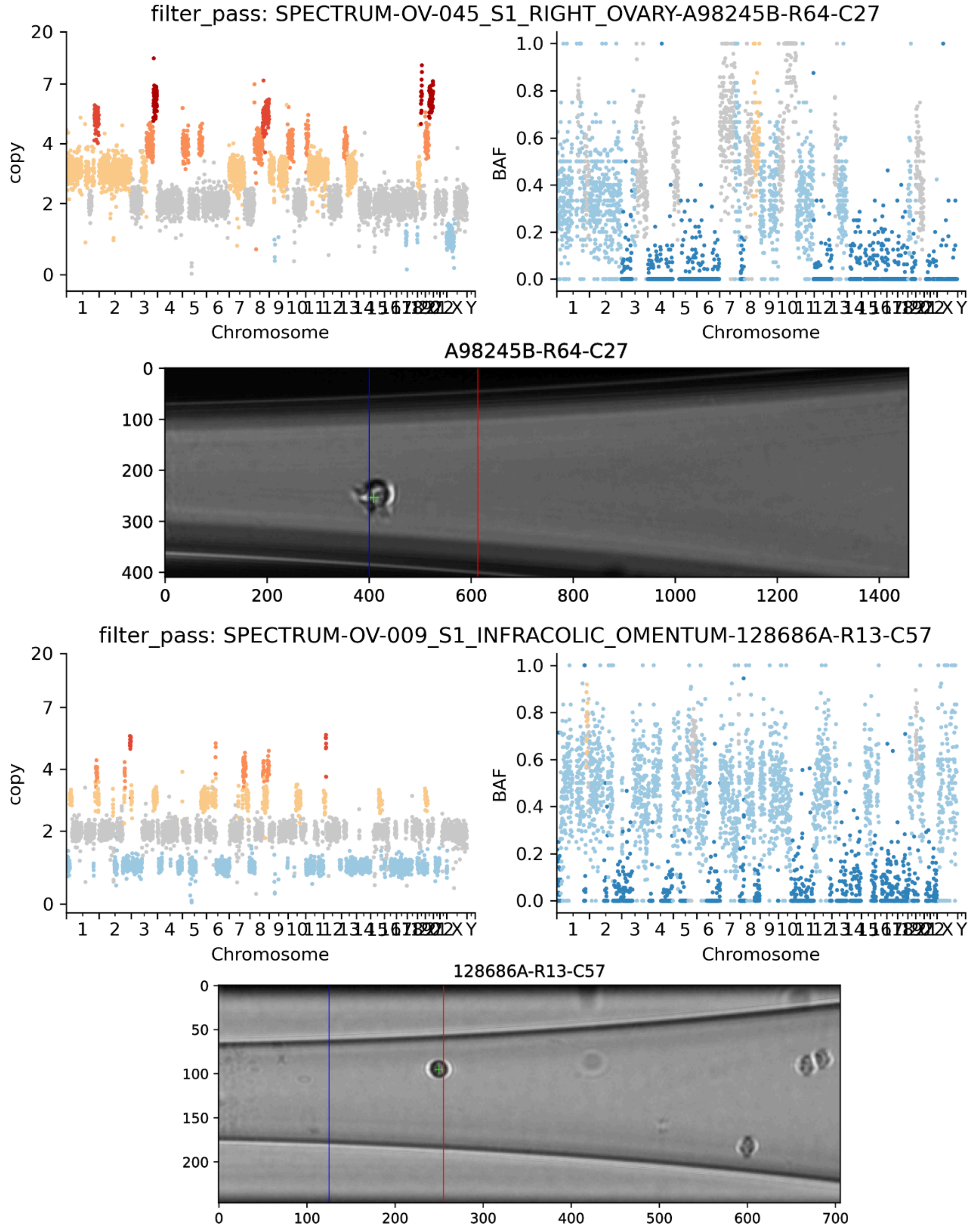


**Supplementary Figure 16. Two example cells that passed all QC filters.** Scatterplots show the normalized and ploidy-scaled read-depth of each bin colored by HMMCopy-inferred copy-number state (left) and B-allele frequency of each bin colored by signals-inferred minor copy-number state (right). Images for each cell (center) show the image taken through the DLP+ nozzle where a green cross indicates the detected cell object, the left vertical line indicates the rightmost edge of the ejection zone, and the right vertical line indicates the rightmost edge of the sedimentary zone.

#### 2. Discerning independent WGD from shared WGD using SNVs

Given two clones (i.e., sets of cells from the same patient) where all cells in both clones appear to have undergone WGD, we used SNVs in 2-copy LOH regions shared among nearly all cells to distinguish whether the cells likely underwent the same WGD event or distinct, independent WGD events. This distinction is equivalent to the question of whether clonal divergence preceded WGD (requiring two independent WGD events; “independent WGD”) or followed WGD (supporting a single shared WGD event; “shared WGD”). Under the assumption that the 2-copy LOH regions common to all cells result from pre-WGD LOH that was followed by doubling to reach 2 copies, the two scenarios can be distinguished by the variant copy numbers of SNVs in these regions (**Supplementary Figure 17**): if the WGD event is shared, then there should be some SNVs present at 1 variant copy in both clones, which were acquired after the WGD but before clonal divergence. However, if there are two distinct WGD events, then each clone should have exclusive mutations at 2 variant copies that were acquired after clonal divergence but before their respective WGDs (note that these may be vastly different in number between the two clones depending on the relative timing of the WGDs in each clones’ ancestry post-divergence). SNVs with other combinations of variant multiplicities do not distinguish between the two scenarios. SNVs at 1 variant copy in one clone and 2 variant copies in the other are discordant with both scenarios, and may suggest that our assumption about the origin of the LOH containing these SNVs is false.

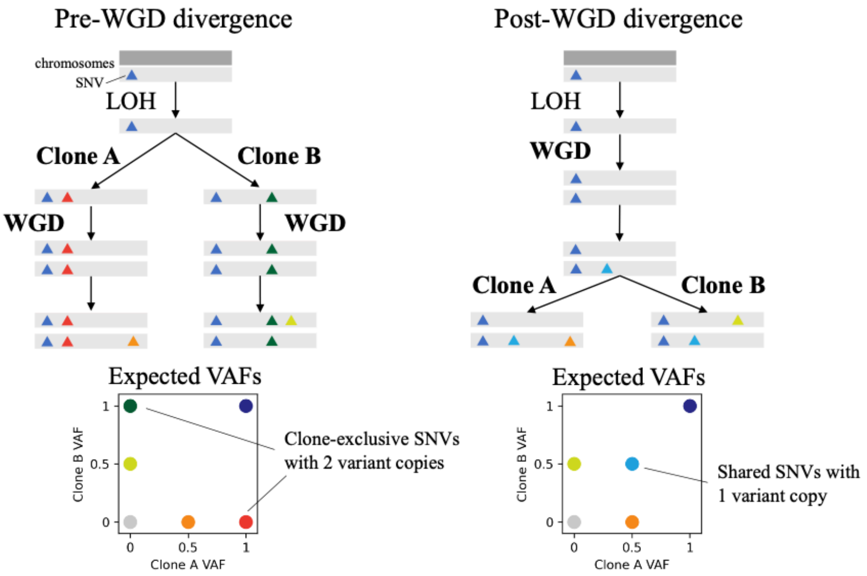


**Supplementary Figure 17. Model for using SNVs in snLOH regions to distinguish between pre-WGD divergence of two clones (left) and post-WGD divergence (right).** Rectangles indicate chromosomes and colored triangles indicate SNVs. The red and dark green SNVs that are each exclusive to a single clone at VAF=1 (2/2 copies) are not possible in the right model, whereas the blue SNVs that are present in both clones at VAF=0.5 (1/2 copies) are not possible in the left model.

For each pair of SBMClone clones inferred for each patient, using a simple binomial model of alternate read counts, we estimated the likelihood of each scenario by inferring the most likely variant copy numbers of each SNV using the respective scenario's permitted combinations of variant copy numbers for the two clones: both scenarios include SNVs with copies 2/2 (shared pre-WGD), 1/0 or 0/1 (exclusive post-WGD), 0/0 (rare or absent); shared WGD includes 1/1 (shared post-WGD pre-divergence), and independent WGD includes 2/0 and 0/2 (exclusive post-divergence post-WGD). We tested the relative fit of these two scenarios using a likelihood ratio test. To evaluate how significantly the observed counts different from the shared-WGD scenario, we generated a null distribution by resampling alternate counts given the observed total read counts under the assumption that the best-fitting variant copy number states under the shared-WGD hypothesis were correct. In **Supplementary Figure 18**, we show these two statistics for the best-scoring bipartition of SBMClone clones for each patient (16 low-WGD patients and 3 patients in which SBMClone did not identify multiple SNV clones were excluded). The two points at the leftmost extreme of the plot correspond to patients patient 071 and patient 133, which each have relatively few (<200) cells in one of the two partitions, leading to relatively few counts and thus a significant permutation test result despite no exclusive VAF-1 SNVs.

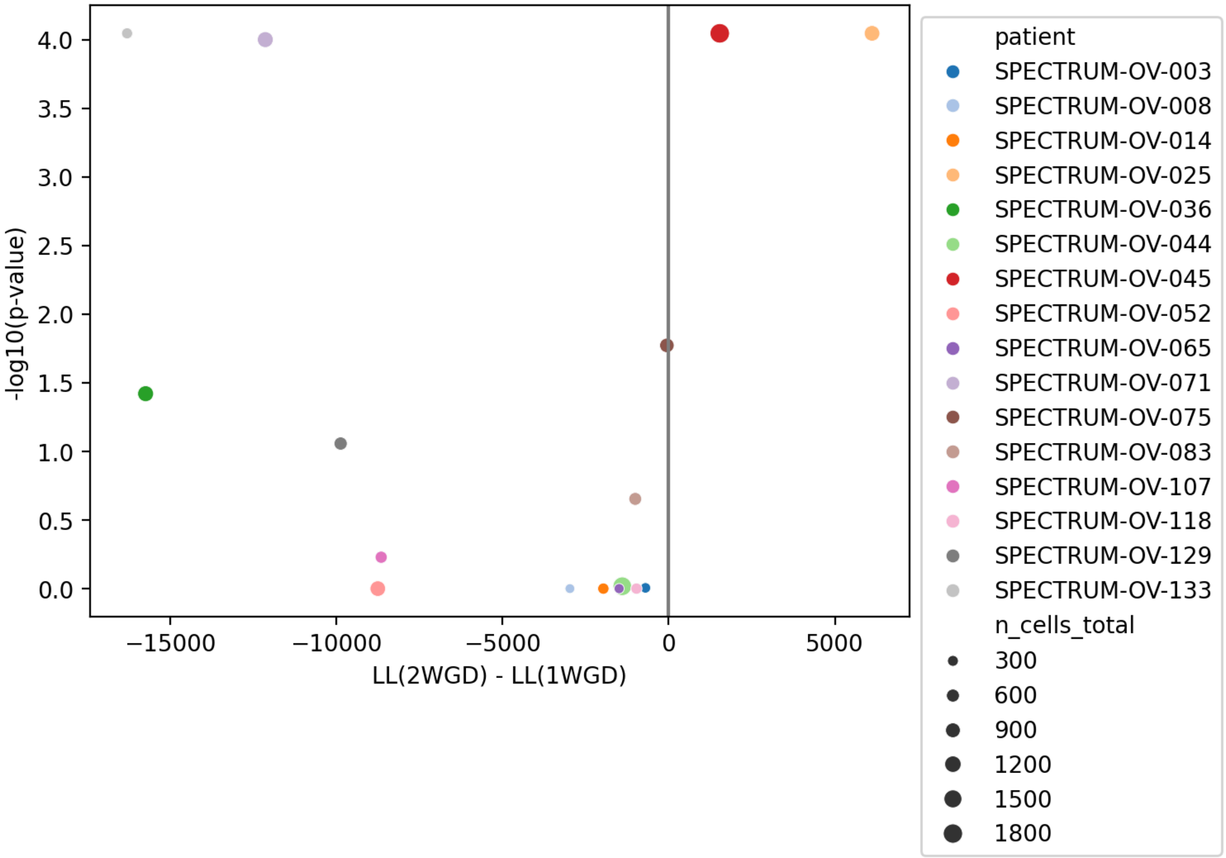


**Supplementary Figure 18. Results from applying likelihood ratio test to SNVs in cnLOH regions in Prevalent WGD patients.** The x-axis shows the difference between the log-likelihood of the independent-WGD scenario ("2-WGD", i.e., pre-WGD divergence) and the log-likelihood of the shared-WGD scenario ("1-WGD", i.e., post-WGD divergence), and the y-axis shows the permutation test p-value assessing the significance of the difference.

We found that two patients, patient 025 and patient 045, showed evidence for multiple WGD events. When we tested all pairs of clones for these two patients to better resolve the WGD events, we found that patient 045 showed evidence for 3 independent WGD events.

#### 3. Benchmarking estimation of chromosomal instability using simulations

##### 3.1. Generation of simulated data

We use CINner[1], a simulation framework for single-cell copy number evolution, to benchmark our methods for estimation of rates of chromosomal instability. Every CINner simulation starts with 1,000 diploid cells. Cell lifetimes are exponentially distributed with the same rate. Cell $i$ has fitness $f^i$ and either divides with probability $p(div)^i$ or dies with probability $1 - p(div)^i$ at the end of its life. The formulae for $f^i$ and $p(div)^i$ follow below.

- Formula for $f^i$ for cell $i$ with CN profile $\{c_k\}$ (where $c_k$ is the count of chromosome $k$) and ploidy $p$ (average of $\{c_k\}$ weighted for chromosomes' lengths):
    - $f^i = 0$ if any of the following conditions is satisfied: (viability checkpoints)
        - $c_k = 0$ for any $k$.
        - $c_k > 8$ for any $k$.
        - $p > 4.5$.
    - Otherwise, $f^i = \prod_k s_k^{2c_k/p}$.
- Formula for $p(div)^i$:
    - $p(div)^i = \frac{N}{1000+N} \cdot \frac{f^i \cdot N}{\sum_{j=1,\cdots,N} f^i}$ where $N$ is the total cell count.
    - The first fraction is to ensure the total population does not diverge from 1,000.
    - The second fraction models the selection.
- Variables:
    - $p_{WGD}$ = probability that a WGD event occurs in a cell division.
    - $p_{misseg}$= probability that a selective missegregation event occurs in a cell division. If the event happens, a chromosome chosen at random (with uniform distribution) is gained in one daughter cell and lost in the other. These events are "selective" because they change the cell's $f^i$ and therefore its $p(div)^i$, detailed below.
    - $p_{neu-misseg}$= probability that a neutral missegregation event occurs in a cell division.
    - $s$ = selection rate.
- Role of selection rate $s$:
    - Every chromosome $k$ is assigned a selection rate $s_k$.
    - For a random half of the chromosomes ("OG chromosomes"), $s_k = 1 + s$. Gains of these chromosomes increase $f^i$ (therefore rendering the cell more selective), and their losses decrease $f^i$.

- For the other chromosomes ("TSG chromosomes"), $s_k = \frac{1}{1+2\cdot s}$. Losses and gains of TSG chromosomes increase and decrease $f^i$ respectively.
- Note that losses of TSG chromosomes are "twice as selective" compared to gains of OG chromosomes (although all events are equally likely to occur). Therefore, TSG chromosome losses are more likely to be fixed, i.e. becoming clonal.

- Simulation process:
  - Step 1 – clonal evolution: Clonal populations are simulated in forward time. At any step, a given clone might give rise to new subclones at given rates of $p_{WGD}$ and $p_{misseg}$. Note that here only the dynamics of clonal cell counts is simulated. Also recorded is the division count in each clone during each time step.
  - Step 2 – sample phylogeny: Once the clonal populations at the final time are known, their phylogeny is simulated in backward time. At each time step, nodes belonging to the same clone are sampled using the clone's division count from step 1 for coalescence.
  - Step 3 – neutral variations: Once the phylogeny of the final cell population is finished, the algorithm simulates neutral missegregations in forward time from the MRCA node to the leaves. At each edge, the count of neutral missegregations is simulated with rate $p_{neu-misseg}$ and count of cell divisions along the edge. Neutral missegregations do not have any role in $f^i$ and therefore do not affect selection forces (already simulated in step 1). However, they are conditioned to not violate the 3 viability checkpoints, or alter the CN in any chromosome that will be changed in a selective event further down the branch in the phylogeny. If no chromosome qualifies, then no neutral missegregations are allowed on the edge.
  - Step 4 – readcounts: Let $\{n_j\}$ be the vectorized CN profile of a given cell, where $n_j$ = CN in bin $j$, and $\{g_j\}$ be the vector of GC content per bin and $r$ be the total read count per cell ($\{g_j\}$ and $r$ are the same for all cells). Then with variables $s$ and $\sigma$, the readcounts per bin of the cell $\{r_j\}$ are simulated as below:
    - $\{n_j'\} = s \cdot \{n_j\} \otimes \{g_j\}$, where $\otimes$ denotes element-wise product
    - $\{n_j''\} \sim Gamma\left(shape = \frac{\{n_j'\}}{\sigma}, scale = \sigma\right)$
    - $\{r_j\} \sim Multinomial\left(size = r, probabilities = \frac{\{n_j''\}}{\Sigma|n_j''|}\right)$

    In the simulations, we use $s = 3906632$ and $\sigma = 0.02642392$ (from previous fitting for OV2295 sample).

Simulation ends after an average of 300 generations, and cell population is expected to stay constant throughout. All cells at the end are sampled, and their CN profiles are recorded.

##### 3.1.2. Results on simulated data

To assess the accuracy of our approach for estimating the rate of chromosome-level copy-number changes from a population of cells, we generated 1,000 simulated datasets varying the missegregation rate and WGD rate (**Supplementary Figure 19**). We applied the same approach for inferring chromosomal instability rates as was used on real data in the main text: we reconstructed phylogenetic trees using MEDICC2, then applied the Sankoff algorithm to reconstruct ancestral copy-number profiles using an independent-bin maximum-parsimony model, and finally called events on each branch using a greedy approach (see Methods for details).

Both the inferred chromosome event rate and the inferred WGD rate were strongly correlated with the corresponding simulation parameters (Pearson correlation $p < 10^{-63}$), although the coefficient of determination for chromosome events ($R^2 = 0.852$) was much higher than that for WGD ($R^2 = 0.496$ for WGD), likely because not all WGD events are detectable or distinguishable from chromosome-level events in sample cells.

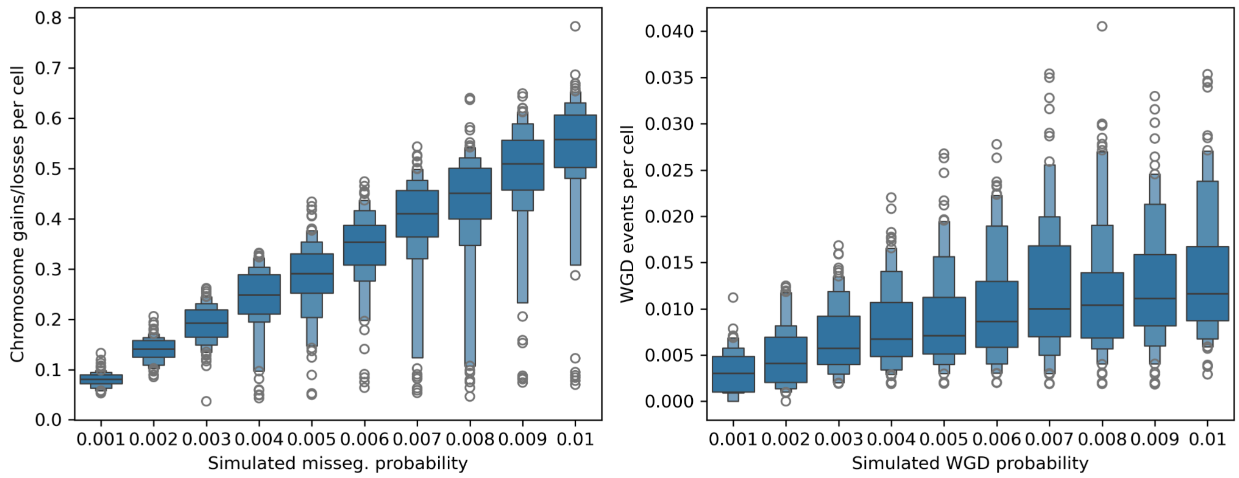


**Supplementary Figure 19: Chromosomal instability rate inference benchmarking results.** Benchmarking inferred chromosome gain/loss rates (y-axis, left) and WGD rates (y-axis, right) using simulated data with varying missegregation probability (x-axis, left) and varying WGD probability (x-axis, right).

#### 4. Extended SBMClone results and validation

##### 4.1. SBMClone block density matrices

SBMClone takes as input a matrix of cells by SNVs and groups cells into clones (y-axis) and SNVs into SNV blocks (x-axis). In **Supplementary Figure 20**, for each patient, each entry is the percentage of nonzero values in the submatrix whose rows consist of the cells assigned to the corresponding clone and whose columns consist of the SNVs assigned to the corresponding SNV block. Only the 31 patients for which SBMClone identified more than 1 clone are shown.

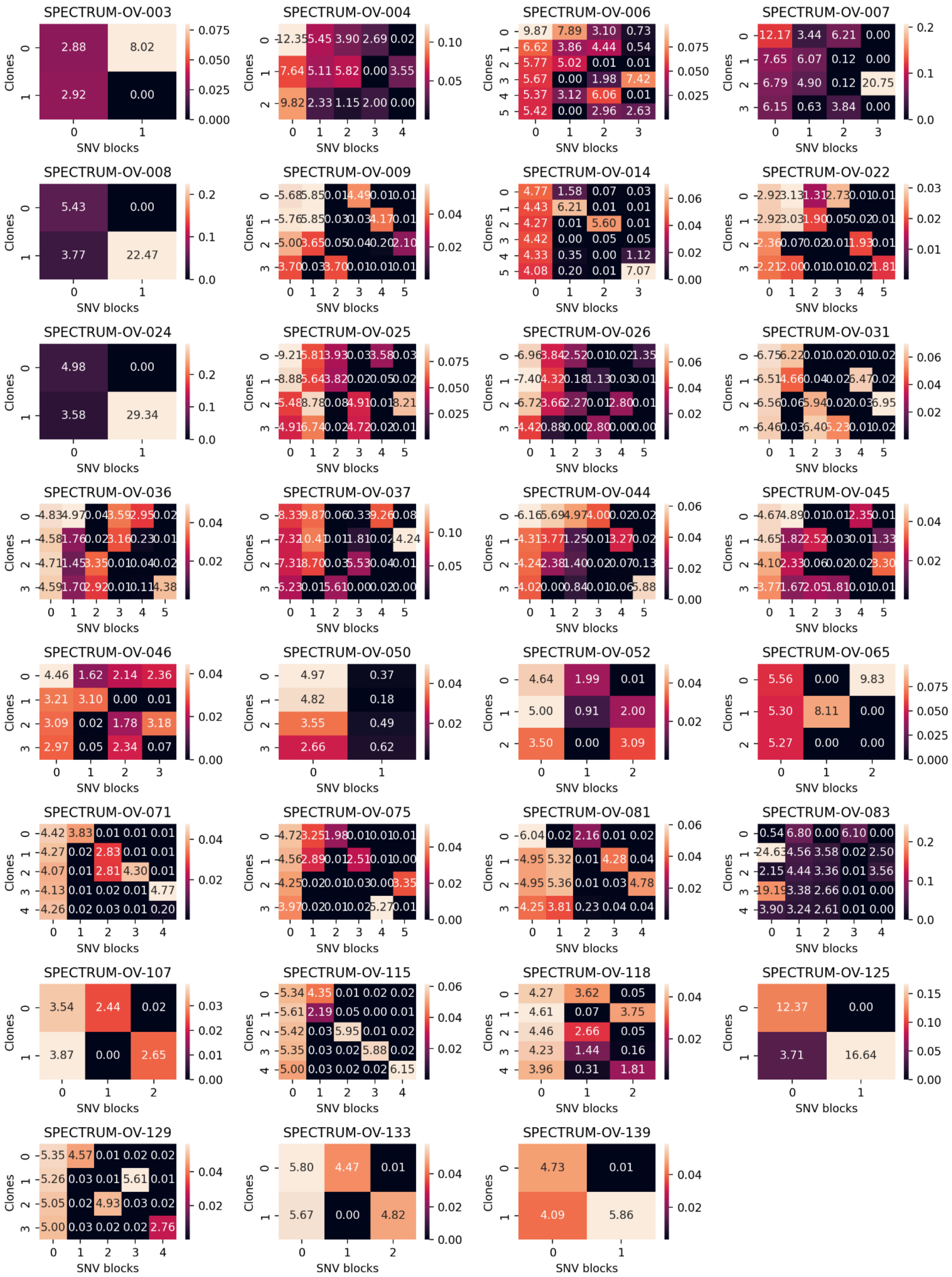
SPECTRUM-OV-003
SPECTRUM-OV-004
SPECTRUM-OV-006
SPECTRUM-OV-007
SPECTRUM-OV-008
SPECTRUM-OV-009
SPECTRUM-OV-014
SPECTRUM-OV-022
SPECTRUM-OV-024
SPECTRUM-OV-025
SPECTRUM-OV-026
SPECTRUM-OV-031
SPECTRUM-OV-036
SPECTRUM-OV-037
SPECTRUM-OV-044
SPECTRUM-OV-045
SPECTRUM-OV-046
SPECTRUM-OV-050
SPECTRUM-OV-052
SPECTRUM-OV-065
SPECTRUM-OV-071
SPECTRUM-OV-075
SPECTRUM-OV-081
SPECTRUM-OV-083
SPECTRUM-OV-107
SPECTRUM-OV-115
SPECTRUM-OV-118
SPECTRUM-OV-125
SPECTRUM-OV-129
SPECTRUM-OV-133
SPECTRUM-OV-139
Clones
SNV blocks

**Supplementary Figure 20: SBMClone results for all patients.** For each clone (set of cells) and SNV block (set of SNVs) inferred by SBMClone, we show the percentage of positive entries in the submatrix whose rows correspond to cells in the clone and whose columns correspond to SNVs in the SNV block.

##### 4.2. Confidence in SBMClone clone assignments

To assess the strength of the assignment of each cell to its SBMClone clone, we evaluated the silhouette score of the SBMClone clusters in SNV space as follows (**Supplementary Figure 21**). For each cell, we computed the proportion of SNVs in each SBMClone SNV block for which it had alternate reads – i.e., we computed the density of each SNV block in each cell. Then, using these density vectors to represent each cell, we computed the silhouette score of the SBMClone clone assignment (with Euclidean distance). Each cell (y-axis) is colored and labeled by the SBMClone block to which it was assigned. Positive silhouette coefficients correspond to cells that are nearby others in the same clone and distant from cells in different clones, whereas negative values represent cells that are no more likely than by chance to be placed in their respective clusters.

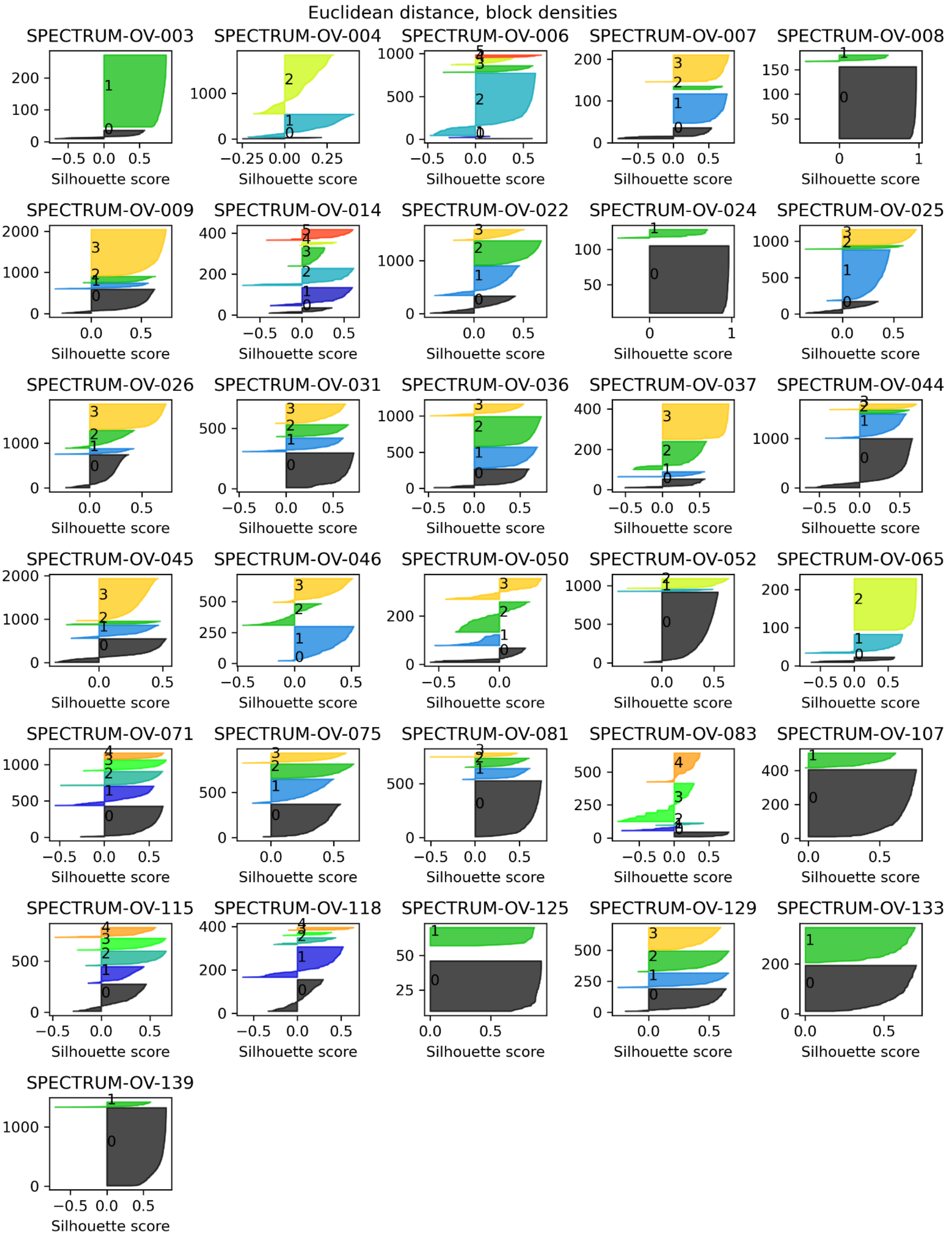


**Supplementary Figure 21: Silhouette scores evaluating the consistency between the SBMClone clones and the cells' SNV block density profiles.**

##### 4.3. Consistency between SBMClone clusters and copy-number states

To evaluate the consistency between SNV-based SBMClone clones and cell copy numbers, we examined the copy-number profiles for the 31 patients in which SBMClone inferred more than 1 clone. Specifically, for each of these patients, we computed the silhouette score of the SBMClone cluster labels using Euclidean distance on the cells' haplotype-specific copy-number profiles (**Supplementary Figure 22**). Positive silhouette scores indicate cells whose SBMClone clone is dense and well-separated in copy-number space, and negative scores indicate cells that are not associated with a well-defined cluster in copy-number space. Overall, most clusters for most patients are fairly consistent between these orthogonal mutation signals, and exceptions often correspond to patients without obvious copy-number subclones (e.g., patient 007).

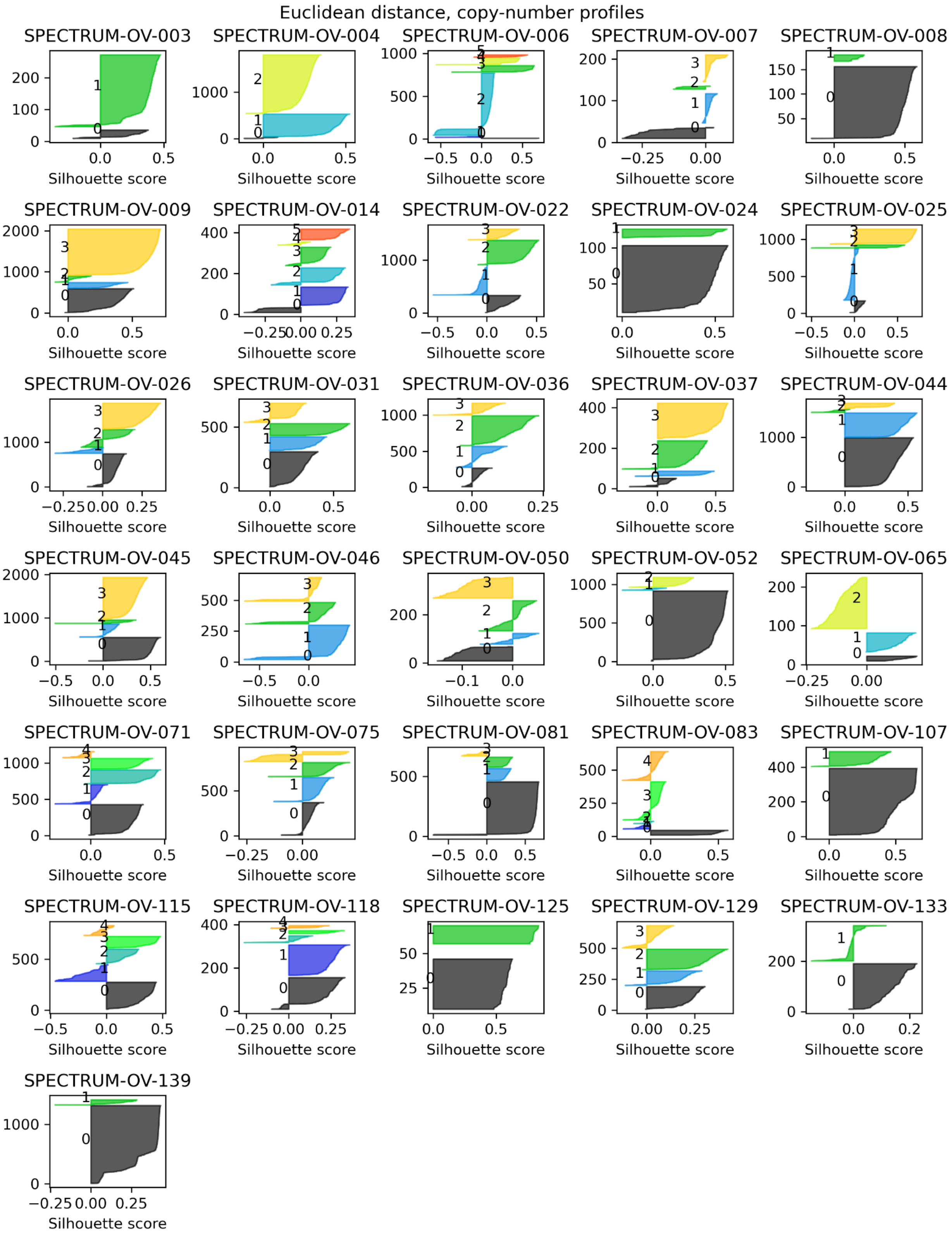


**Supplementary Figure 22: Silhouette scores evaluating the consistency between the SBMClone clones and the cells' haplotype-specific copy-number profiles.**

#### 5. Assignment of SNVs to clone tree branches

To assess how well SNVs were assigned to clone tree branches (see Methods for details), we grouped all (SNV, clone) pairs for each patient into those in which the SNV was inferred to be present in the clone (i.e., the SNV was assigned to a branch ancestral to the clone) and those in which the SNV was inferred to be absent (i.e., the SNV was assigned elsewhere on the tree or inferred to be absent from all clones). **Supplementary Figure 23** shows the SNV VAFs in clones in which the corresponding SNV was inferred to be present, colored by the clone. **Supplementary Figure 24** shows VAFs for those (SNV, clone) pairs in which the SNV was inferred to be absent.

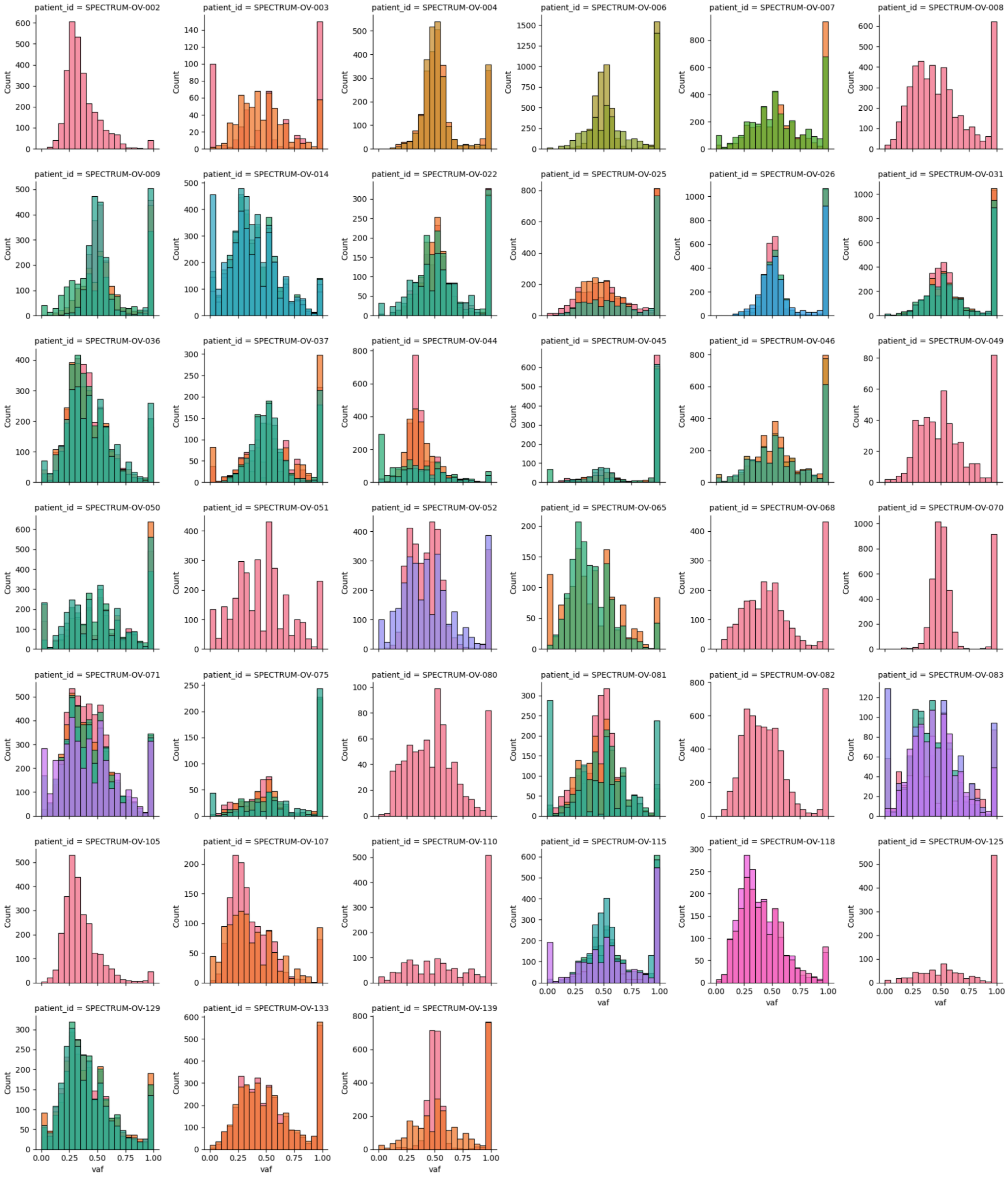


**Supplementary Figure 23: VAFs for SNVs inferred to be present in the corresponding clone for each patient.**

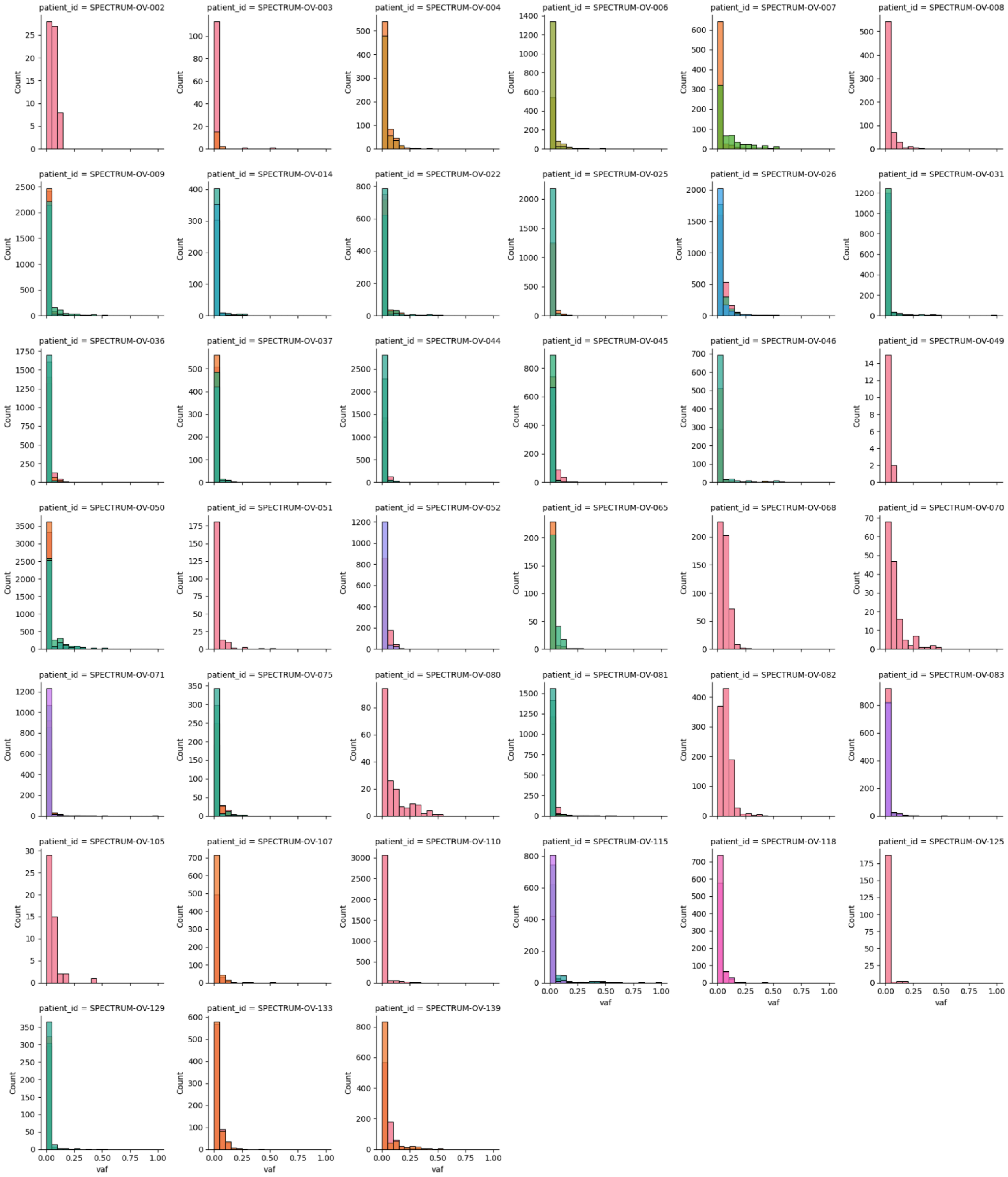


**Supplementary Figure 24: VAFs for SNVs inferred to be absent in the corresponding clone for each patient.**
